# Evolution of leaf-cutter behavior in bees (Hymenoptera: Megachilidae) as inferred from total-evidence tip-dating analyses

**DOI:** 10.1101/543082

**Authors:** Victor H. Gonzalez, Grey T. Gustafson, Michael S. Engel

**Affiliations:** Undergraduate Biology Program and Department of Ecology and Evolutionary Biology, Haworth Hall, 1200 Sunnyside Avenue, University of Kansas, Lawrence, Kansas 66045-7523, USA; Division of Entomology, Natural History Museum, 1501 Crestline Drive – Suite 140, University of Kansas, Lawrence, Kansas 66045-4415, USA; Department of Ecology and Evolutionary Biology, Haworth Hall, 1200 Sunnyside Avenue, University of Kansas, Lawrence, Kansas 66045-7523, USA; Division of Invertebrate Zoology, American Museum of Natural History, Central Park West at 79^th^ Street, New York, New York 10024-5192, USA

## Abstract

A unique feature among bees is the ability of some species of *Megachile s.l*. to cut and process fresh leaves for nest construction. The presence of razors between the female mandibular teeth (interdental laminae) to facilitate leaf-cutting (LC) is a morphological novelty that might have triggered a subsequent diversification in this group. However, we have a limited understanding of the evolutionary origins of this behavior and associated structures. Herein, we use total-evidence tip-dating analyses to infer the origin of LC bees and patterns of variation of interdental laminae. Our datasets included five nuclear genes, representatives of all fossil taxa, 80% of the extant generic-level diversity of Megachilidae, and the full range of generic and subgeneric diversity of Megachilini. Our analyses support the notion of a recent origin of LC bees (15–25 Ma), casting doubts on Eocene trace fossils attributed to these bees. We demonstrate that interdental laminae developed asynchronicaly from two different structures in the mandible (teeth or fimbrial ridge), and differ in their phenotypic plasticity. Based on the phylogenetic results, we propose robust classificatory solutions to long-standing challenges in the systematics of Megachilidae. We discuss the implications of our findings as a foundational framework to develop novel evolutionary, ecological, and functional hypotheses on this behavior.

## Introduction

Bees are a monophyletic group of hymenopteran insects that arose from the apoid wasps at least 125 Ma during the Cretaceous and diversified in conjunction with flowering plants (*e.g*., Engel, 2001; Michener, 2007; Michez *et al*., 2012). Bees are highly valuable to science and society because of their unique biology as pollinators of both wild and cultivated plants, which provide products and services, including social and cultural values (*e.g*., Gonzalez *et al*., 2013; Potts *et al*., 2016). There are more than 20,000 species worldwide, with the most widely accepted classification of bees recognizing seven extant families, more than 50 tribes, and about 500 genera (Michener, 2007). Bees are diverse in their morphologies and biologies. For example, most are robust, hairy, and forage during the day (*e.g*., bumble bees), but many are slender, nearly hairless (*e.g*., yellow-faced bees of the genus *Hylaeus* spp.), and visit flowers at night or at dusk (*e.g*., nocturnal sweat bees of the genus *Megalopta* spp.). Most bees do not live in colonies or make honey. The great majority are solitary and some are cleptoparasites on other bees. Females of solitary species build their own nests and do not have contact with their offspring, whereas cleptoparasitic species leave their eggs in the cells of their host bees (Michener, 2007). Unlike wasps, bees use pollen to feed their brood, but a few species use carcasses of dead animals as a source of protein or even tears from mammals in addition to, or in lieu of, pollen (*e.g*., Roubik, 1982; Bänziger *et al*., 2009).

A unique behavior among bees is the ability to cut and process fresh leaves for nest construction. Using their mandibles, the bees cut and collect circular to elliptical leaf pieces, leaving distinct excision patterns along the margin of leaves. The bees then use these leaf pieces to line and separate the brood cells, which they build in the ground or inside pre-existing cavities (*e.g*., Michener, 1953). This leaf-cutter (LC) behavior is exclusive to a group of solitary bees in the genus *Megachile* Latreille (Groups 1 and 3 of Michener, 2000, 2007), a megadiverse, cosmopolitan group that is both morphologically and biologically heterogeneous, which has resulted in a problematic taxonomy (Fig. 1). This genus includes a number of invasive species (*e.g*., Cane, 2004; Rasmussen *et al*., 2012), many highly promising pollinators, and *M.* (*Eutricharaea*) *rotundata* (Fabricius), the most intensively managed and produced solitary bee in the world for the production of alfalfa (Pitts-Singer and Cane, 2011).

**Fig 1.**
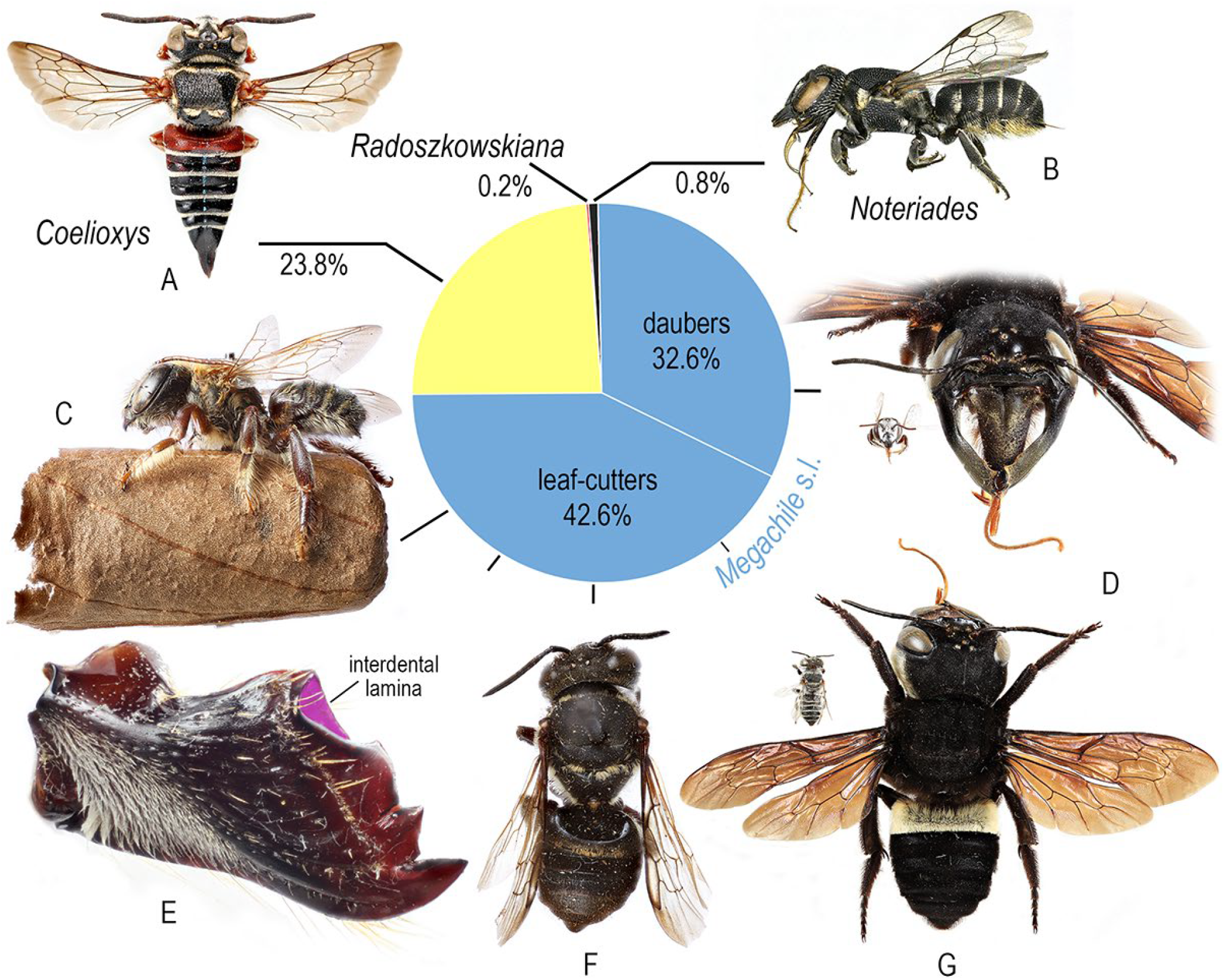
Species richness of currently recognized genera in the bee tribe Megachilini. **A**. Dorsal habitus of a female of *Coelioxys* sp. **B**. Lateral habitus of a female of *Noteriades spinosus* Griswold and Gonzalez. **C**. Male of *Megachile* (*Zonomegachile*) sp. on top of a brood cell built with leaf pieces. **D**. Facial habitus of leaf-cutter *M*. (*Eutricharaea*) *minutissima* Radoszkowski (left) and dauber bee *M*. (*Callomegachile*) *pluto* (Smith) (right). **E**. Outer surface of the female mandible of *M*. (*Leptorachis*) *laeta* Smith, a leaf-cutter bee, showing interdental lamina in pink. **F**. Dorsal habitus of *M*. (*Rhyssomegachile*) *kartaboensis* Mitchell. **G**. Dorsal views of *M*. *minutissima* (upper left) and *M*. *pluto* (right). Photographs are not at the same scale, except for the large and small species compared in figures D and G.

A good knowledge of LC behavior is essential to gain a better understanding of species’ biologies, predict species distributions, and improve current management practices for commercial and conservation purposes (Sinu and Bronstein, 2018). However, little information is available on which species of plants are used by LC bees and by which bee species. In addition, the majority of records are from common bees in urban or agricultural areas (MacIvor, 2016; Kambli *et al*., 2017; Sinu and Bronstein, 2018). Such limitations are surely a reflection of the challenges associated with finding nests, identifying plants from leaf fragments, and LC bees’ taxonomic problems (Michener, 2007; Gonzalez *et al*., 2013). Significantly, less information is yet available on the evolutionary history of this behavior and the mandibular structures involved in leaf cutting.

Unlike other bees, megachilids do not line their cells with hydrophobic secretions of the Dufour’s gland; instead, they rely on the physicochemical properties of the foreign material used for nesting (Williams *et al*., 1986). In the case of LC bees, certain phytochemicals (*e.g*., saponins) might increase larval mortality (Horne, 1995), while others (*e.g*., flavonoids, phenols, terpenoids) might decrease it by providing protection against microbes (MacIvor, 2016; Sinu and Bronstein, 2018). Therefore, it is not surprising that available data suggest that LC bees are highly selective in their plant and leaf choices, avoiding latex-producing plants, and preferring species with glabrous leaves, particularly in the families Fabaceae and Rosaceae (Michener, 1953; Kambli *et al*., 2017; Sinu and Bronstein, 2018).

The female mandible of LC bees varies considerably in its overall length and shape, as well as in the number and shape of its teeth (Fig. 2). It also has a distinct lamina in one or more spaces between the teeth, which authors have called ‘interdental lamina’ (Pasteels, 1965) or ‘cutting edges’ (Michener, 1962). Doubtless, this structure is an evolutionary novelty among bees because it is unique to this group and its appareance might have triggered a subsequent diversification within LC bees. Presently, the total number of species exhibiting LC behavior accounts for about 57% of the species in the tribe Megachilini (Michener, 2007).

**Fig 2.**
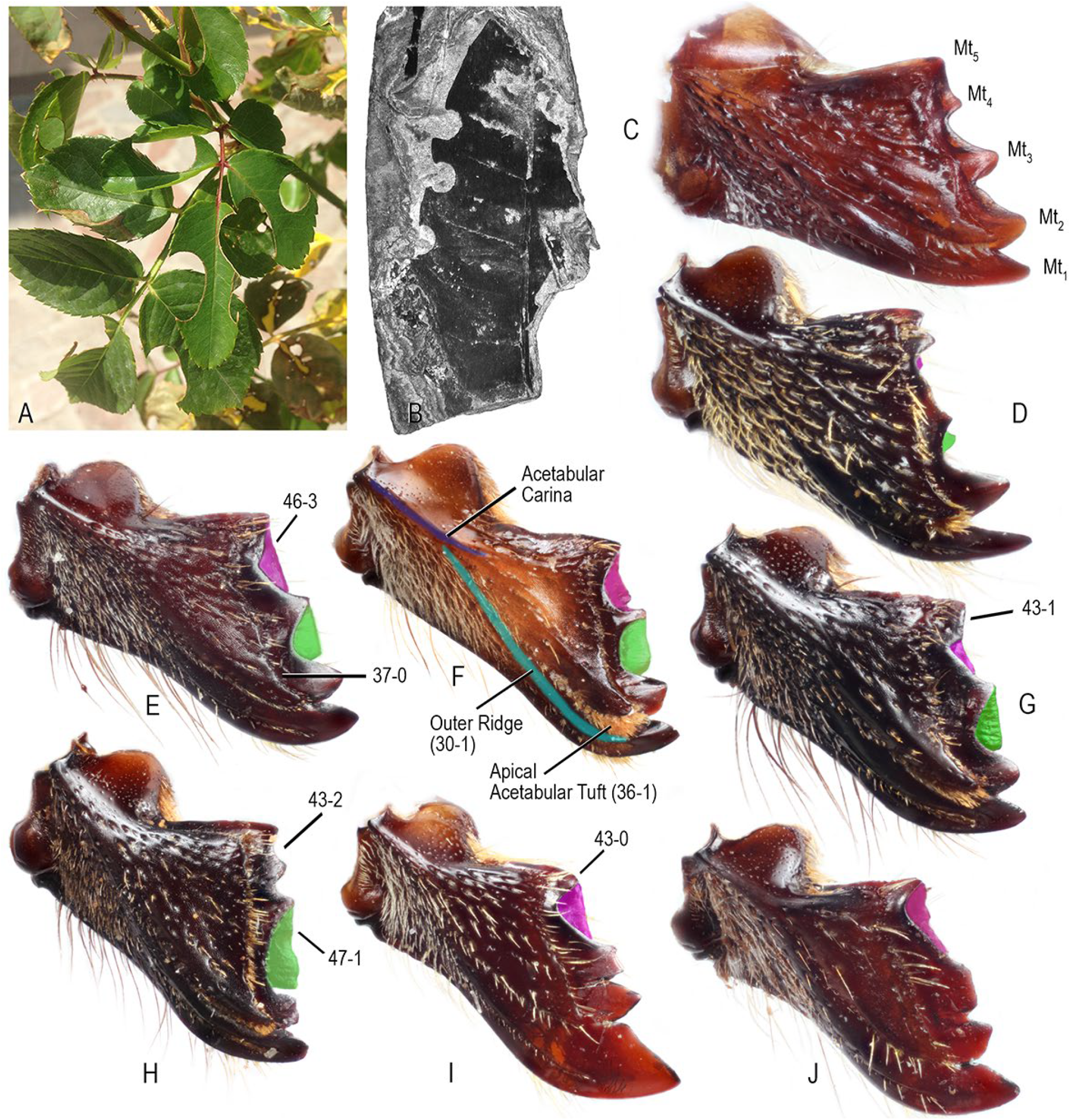
Leaves excisions and sample of the morphological diversity among the female mandible of leaf-cutter bees. **A.** Leaves of *Rosa* sp. (Rosaceae) from Lesvos, Greece. **B.** Fossil leaf cut (Fabaceae) from Eckfeld Maar, Germany (∼43 Ma). **C–J**. Outer view of the mandible showing interdental laminae in green (odontogenic) and pink (ctenogenic). **C**. *Megachile* (*Chrysosarus*) *parsonsiae* Schrottky. **D**. *M*. (*Rhyssomegachile*) *simillima* Smith. **E.** *M*. (*Pseudocentron*) *pruina* Smith. **F**. *M*. (*Zonomegachile*) sp. **G**. *M*. (*Moureapis*) *maculata* Smith. **H.** *M*. (*Melanosarus*) *xylocopoides* Smith. **I.** *M*. (*Acentron*) *albitarsis* Cresson. **J.** *M*. (*Leptorachis*) *petulans* Cresson. Interdental laminae highlighted in green (odontogenic) and pink (ctenogenic). Abbreviations: Mt= mandibular tooth.

The presence or absence of this interdental lamina, as well as its size and shape, varies among species and species groups. Such variations correlate with different modes of leaf-cutting behavior and have been useful in the taxonomy of the group. Species with a lamina that entirely fills the space between teeth generally exhibit extensive LC behavior; their brood cells are entirely made of smooth-margined leaf pieces. In contrast, species with incomplete lamina (not entirely filling spaces between teeth) or without it, have more limited LC behavior, with their brood cells made of a combination of mud and leaf or petal pieces, which are irregularly cut, often with serrate margins (Michener, 2007). The absence of this lamina in the mandible of some species that still exhibit LC behavior indicates that other structures are also involved in leaf cutting. Similarly, the morphological diversity of the mandible also suggests different mechanical solutions to diverse functional problems. However, no one has yet attempted to understand the origins and patterns of variations of these mandibular structures using a phylogenetic framework.

Several authors have recorded fossilized dicotyledonous leaves with distinctive cuts along their margins, similar to those caused by LC bees (Figs. 2A, B). Those trace fossils are from deposits in Europe, North and South America, and the oldest is approximately 60 Ma (*e.g*., Wedmann *et al*., 2009; Michez *et al*., 2012). Comparative analyses of the ellipse eccentricity between leaf discs of brood cells of living species and fossil excisions, support the attribution of these trace fossils to LC bees (Sarzetti *et al*., 2008). However, molecular analyses using a node-dating approach, which places the oldest fossil to the youngest internal node and thus imposes the age of the fossil as a minimum age constraint, suggest that LC bees originated around 20–25 Ma (Litman *et al*., 2011; Trunz *et al*., 2016). Other dating approaches might be useful for investigating this temporal discrepancy, such as Bayesian total-evidence tip dating, which utilizes morphological data to infer the placement of fossils within the phylogeny (as terminals or ‘tips’) in order to calibrate the tree. Therefore, tip-dating does not require the *a priori* constraint of taxa to nodes in order to generate age estimates, and allows the use of all available fossils within a group, extending age estimates beyond the minimum age for clades (Ronquist *et al.*, 2012a). (Ronquist *et al*., 2012a). Unfortunately, such analyses are not yet available for megachilids nor for any other group of bees.

Considering the biological importance of the LC behavior in the evolution and diversification of this group of pollinators, we set the following goals: First, to assess the origin of LC behavior using a Bayesian total-evidence tip-dating approach. Second, to determine the possible origins of the interdental lamina in the female mandible. Third, to explore possible patterns of variation of the interdental lamina. Fourth, to examine the implications of our phylogenetic results on the classification of Megachilidae and Megachilini. In the following sections, we provide an overview of the diversity and phylogeny of megachilids, highlighting outstanding problems in their classification.

### Diversity and phylogeny of Megachilidae

Megachilidae is the third largest bee family containing more than 4000 species worldwide (Michener, 2007; Ascher and Pickering, 2018). The nesting biology of megachilids is collectively more diverse than any other bee group, as these bees exploit a wide variety of nesting materials and substrates. For example, they use mud, petals, leaves (intact pieces or macerated to a pulp), resins, gravel, and plant trichomes to build their brood cells in the soil, attached to twigs, under surfaces of rocks, or inside pre-existing cavities including man-made constructions (*e.g*., Rozen *et al*., 2010; Gonzalez and Griswold, 2013). The most recent higher-level classificatory proposal for the family (Gonzalez *et al*., 2012) recognizes four subfamilies and nine tribes, three of which are extinct (Ctenoplectrellini, Glyptapini, and Protolithurgini). Morphological (Gonzalez *et al*., 2012) and molecular data (Litman *et al*., 2011) support the monophyly of these tribes, except that of Osmiini, which has long been suspected to be paraphyletic (*e.g*., Engel, 2001; Michener, 2007; Praz *et al*., 2008). In these morphological and molecular analyses, either Megachilini or Megachilini + Dioxyini renders Osmiini paraphyletic. The phylogenetic relationships of Dioxyini are also still not clear. This small monophyletic group of cleptoparasitic bees (∼36 spp.) appeared as sister of *Aspidosmia* Brauns (Aspidosmiini) in the molecular analysis, but in the morphological analysis it was the sister group of Megachilini.

### Diversity and phylogeny of Megachilini

Megachilini contains about half of the species of the family (∼2000 spp.: Michener, 2007; Ascher and Pickering, 2018). The most widely used classificatory proposal for bees worldwide (Michener, 2007) recognizes a free-living genus *Megachile* Latreille, and two cleptoparasitic genera, *Coelioxys* Latreille and *Radoszkowskiana* Popov. Gonzalez *et al*. (2012) recently transferred from the Osmiini another free-living genus, *Noteriades* Cockerell (Fig. 1B). All genera of Megachilini seem monophyletic, except for *Megachile*.

*Coelioxys* is cosmopolitan in distribution and includes about 470 species grouped in 15 subgenera in the classification of Michener (2007), but several neotropical taxa synonymized by him are still recognized by some authors (*e.g*., Moure *et al*., 2007). *Coelioxys* are commonly collected bees and frequently found parasitizing other megachilids and some apids. Multiple authors have studied their behavior and immatures (references in Michener, 2007). *Radoszkowskiana* includes only four species restricted to the Palearctic region, which are morphologically and behaviorally similar to *Coelioxys* (Rozen and Kamel, 2007). Rocha Filho and Packer (2016) explored the phylogenetic relationships among the subgenera of *Coelioxys*.

*Noteriades* includes 16 species that occur across tropical and subtropical regions of sub-Saharan Africa, India, and Southeast Asia. The biology of this group of bees is unknown (Griswold and Gonzalez, 2011). Species are small, heriadiform or hoplitiform in body shape, and non-parasitic considering the presence of a metasomal scopa. Griswold (1985) first suggested the close relationship of this genus with Megachilini, which molecular (Praz *et al.*, 2008; Litman *et al*., 2011) and morphological analyses (Gonzalez *et al*., 2012) further supported it.

Remaining species of Megachilini (∼1500 spp.) are all currently placed in *Megachile*. This is the most ecologically and morphologically diverse group of the family. It includes LC bees and species that primarily use mud or resins as nesting materials. It occurs in a wide diversity of habitats on all continents, ranging from lowland tropical rain forests, deserts, to high elevation environments. In appearance, species of *Megachile* range from nearly bare, elongate, parallel-sided bees to robust, hairy bees resembling some smaller bumble bee species; their body length ranges from about 4 mm in *M.* (*Eutricharaea*) *minutissima* Radoszkowski, to nearly 40 mm in *M.* (*Callomegachile*) *pluto* (Smith), the longest bee in the world (Figs. 1D, G). As we briefly describe below, the taxonomy of *Megachile* is problematic and its phylogenetic relationships largely unexplored.

### What is the genus Megachile?

The concept of *Megachile* has changed multiple times since its conception. Latreille (1802) proposed *Megachile* for the European species *Apis centuncularis* Linnaeus, and it initially included not only species of this genus as currently defined, but also species that now belong to different tribes of Megachilidae. Later, Lepeletier de Saint Fargeau (1841) proposed the genus *Chalicodoma* for another European species, *Apis muraria* Olivier. During the second half of the 1800’s, as well as during the first decades of the 1900’s, several authors (*e.g*., Smith, 1865; Thomson, 1872; Provancher, 1882; Meunier, 1888; Friese, 1899; Robertson, 1901, 1903; Cockerell, 1907, 1922; Mitchell, 1924) proposed a number of generic or subgeneric names for closely allied taxa to *Megachile* from different regions of the world. Until the late 1800’s, most authors recognized both *Megachile* and *Chalicodoma* as morphologically and biologically distinct groups, the first consisting of LC bees and the second of species that use mud or resins to build their nests (*e.g*., Gerstaecker, 1869; Radoszkowsky, 1874; Taschenberg, 1883). However, Dalla Torre (1896) appears to be the first to treat *Chalicodoma* as a subgenus of *Megachile*, a position followed by Friese (1898, 1899, 1909, 1911a,b). The latter author (Friese, 1911a) also recognized two previously described taxa, *Thaumatosoma* Smith and *Stellenigris* Meunier, as genera closely related to *Megachile*.

Mitchell (1934) also considered *Megachile* in a broad sense following earlier authors. He regarded *Thaumatosoma* and other generic names proposed until then as subgenera of *Megachile*, including some that Friese (1911a,b) did not mention. In subsequent years, Mitchell (1935a,b, 1936, 1937a,b,c,d, 1943) proposed several new taxa from the Western Hemisphere and revised their species in a series of monographs that stand until today as major or only resources of identification for these bees.

Based on the generic concepts previously used and the discovery of some morphological features that correlated with the nesting behavior, Michener (1962, 1965) divided *Megachile* into three genera (*Chalicodoma*, *Creightonella* Cockerell, and *Megachile*). *Chalicodoma* included Eastern Hemisphere species with a strongly convex and rather parallel-sided metasoma and female mandibles without interdental laminae (Figs. 1D, G); those morphological features are associated with narrow burrows and the use of mud or resin as nesting materials. In contrast, *Megachile* included a cosmopolitan group of bees with a flattened metasoma and female mandibles with interdental laminae, features that allow them to cut and use leaf or petal pieces for constructing cells in wider burrows. *Creightonella* combined features of both genera, a female mandible with interdental laminae to cut leaves, and a strongly convex, parallel-sided metasoma. *Creightonella* included a relatively small number of species (50 spp.) restricted to the Eastern Hemisphere. Pasteels (1965) also independently developed the same classificatory scheme of Michener (1962, 1965) when considering the African fauna. Both authors, C.D. Michener and J.J. Pasteels, not only described several subgenera within *Megachile* and *Chalicodoma*, but also rendered as subgenera a few other generic names proposed at the time.

In 1980, when T.B. Mitchell revised the LC bees from the Western Hemisphere, he adopted the multigeneric proposal of Michener (1962, 1965) in recognizing three genera. However, he further divided *Megachile* into six genera, each with multiple subgenera. Although he was not concerned with the Eastern Hemisphere fauna, he made an effort to summarize and place this fauna within his classificatory scheme, which was not widely adopted (Appendix S1).

Despite having divided *Megachile* into three genera in the 1960’s, Michener (2000, 2007) no longer recognized them when treating the world fauna because of the exceptions and intergradations he later observed in the main morphological features, as well as for almost all other features he had previously used to characterize these groups. In particular, *M*. (*Megella*) Pasteels and *M*. (*Mitchellapis*) Michener represented major problems within his system. Although Pasteels (1965) and Michener (1965) initially placed both taxa in *Megachile*, they exhibit features of both *Megachile* and *Chalicodoma*. For example, typical *Megachile* characteristics are the interdental laminae in the female mandible and the apex of the female sixth sternum with a fringe of short, dense plumose setae. Features typical of *Chalicodoma* include the elongate, parallel-sided body, apex of the female tibiae with a distinct, sharp spine, and the presence of setae on the lateral margins of the male eighth sternum. Michener (2000, 2007) also synonymized certain subgeneric names that authors created for unusual species and organized the more than 50 subgenera into three informal groups, which corresponded to each genus that he previously recognized in the 1960’s. That is, Groups 1, 2, and 3, are equivalent to the genera *Megachile*, *Chalicodoma*, and *Creightonella*, respectively, in Michener’s (1962, 1965) earlier classification (Appendix S1). Because of the presence of marginal setae on the eighth sternum of the male, Michener (2000, 2007) placed these two “problem” taxa (*Mitchellapis* and *Megella*) in Group 2 (*Chalicodoma*), not in Group 1 (*Megachile*) as he (Michener, 1965) and Pasteels (1965) initially assigned them. Another subgenus that also bridged the gap between *Megachile* and *Chalicodoma* was *M.* (*Chelostomoda*). Michener (1962) described this group as a subgenus of *Chalicodoma* even though it also possesses interdental laminae as in *Megachile*.

Today, there is no consensus in the classification of *Megachile*. Some authors still follow Michener’s earlier classification (Michener, 1962, 1965) in recognizing the genera *Chalicodoma*, *Creightonella*, and *Megachile*, including several subgenera that were proposed for species with aberrant or unusual morphologies and that Michener (2000, 2007) synonymized (*e.g*., Silveira *et al*., 2002; Durante and Abrahamovich, 2006; Moure *et al*., 2007; Ornosa *et al*., 2007). Other authors (Trunz *et al*., 2016) recognize a few other taxa at the generic level, as they were initially proposed [*M*. (*Gronoceras*) Cockerell and *M*. (*Heriadopsis*) Cockerell] or were suggested by Michener (2007) as an alternative classification [*M.* (*Matangapis*) Baker and Engel]. The species-level systematics of *Megachile s.l*. (*sensu* Michener 2000, 2007) is also problematic and thus species identifications are challenging in most groups. Taxonomic revisions for the majority of the subgenera are not available, keys to species are lacking, and many species have not been properly associated with any of the known subgenera (Michener, 2000, 2007). Even in North America, many species are still known from a single sex or from a small number of specimens (Sheffield and Westby, 2007; Gonzalez *et al*., 2013).

The phylogenetic relationships among the genera of Megachilini, as well as the subgenera of *Megachile s.l*., are largely unexplored. Michener (2000, 2007) suggested that *Coelioxys* might render *Megachile s.l*. paraphyletic because it shares some morphological traits, particularly with *M.* (*Chelostomoides*) Robertson. Likewise, the recent inclusion of *Noteriades* in Megachilini might also render *Megachile s.l*. paraphyletic considering that this genus shares the presence of arolia (a rare feature typical of Osmiini) with *M.* (*Matangapis*) and *M*. (*Heriadopsis*). An unpublished dissertation (Gonzalez, 2008) explored the relationships within Megachilini using morphological data but did not include *Noteriades*. Similarly, the positions of *M*. (*Matangapis*) and *M*. (*Heriadopsis*) were unclear in a recent molecular analysis (Trunz *et al*., 2016), as both taxa nested in a clade consisting of *Coelioxys* and *Radoszkowskiana*. Doubtless, species-level revisionary studies and phylogenetic analyses are required to develop a more stable taxonomy and phylogeny-based classification of *Megachile s.l*.

### Fossil record

Engel (1999, 2001), Engel and Perkovsky (2006), and Michez *et al*. (2012) summarized the fossil record for Megachilidae. The extinct tribes Protolithurgini, Ctenoplectrellini, and Glyptapini contain several species in five genera (*Protolithurgus* Engel, *Ctenoplectrella* Cockerell, *Glaesosmia* Engel, *Friccomelissa* Wedmann, Wappler, and Engel, and *Glyptapis* Cockerell), most, but not all, from Eocene Baltic amber (33.9–56 Ma). The first tribe is sister to all Lithurginae while the remaining two are sisters to all Megachilinae, except Aspidosmiini (Gonzalez *et al*., 2012). For Megachilini, most records are trace fossils of dicotyledonous leafs with excisions along the margins, similar to those caused by LC bees of the genus *Megachile s.l*. (Wedmann *et al*., 2009; Engel and Perkovsky, 2006; Sarzetti *et al*., 2008). Body compressions are few and have not been associated to any subgenus. *Megachile glaesaria* Engel, from the Miocene Dominican amber (23–30 Ma), is the best-preserved fossil of Megachilini. Engel (1999) noted the close resemblance of this species to some species of the extant North American *M.* (*Chelostomoides*). However, he placed it in its own subgenus, *M*. (*Chalicodomopsis*), because of the presence of a small inner tooth in the pretarsal claws and some wing features, which appear to be intermediate between Megachilini and Anthidiini. He also suggested that *M. glaesaria* might be a basal member of the Group 2 of subgenera or sister to all Megachilini. To date, the phylogenetic position of *M. glaesaria* is unknown.

## Material and methods

To assess the origin of LC bees, we conducted two sets of total-evidence tip-dating analyses aimed at obtaining more accurate divergence time estimates. First, we conducted a phylogenetic analysis of all tribes in the family Megachilidae. Then, we used the divergence time estimates generated from that analysis to inform priors for the phylogenetic analysis of the genera of Megachilini. We used the morphological data matrix of Gonzalez *et al*. (2012) for the tribal-level analysis of Megachilidae and built a morphological data matrix for the generic-level phylogeny of Megachilini. Below, we provide information on the taxonomic coverage and character statements used in the newly built morphological data matrix, as well as on the phylogenetic analyses conducted on both morphological and molecular datasets. Unless otherwise indicated, we followed Michener’s (2000, 2007) subgeneric classification of *Megachile s.l*. to facilitate comparisons (Appendix S1).

### Taxon sampling

The tribal-level analysis of Megachilidae includes all tribes, representatives of all fossil taxa, and 80% of the extant generic-level diversity of the family (Appendix S2). For the generic analysis of Megachilini, we used eight taxa as outgroups based on the phylogeny of Gonzalez *et al*. (2012) and 114 species of Megachilini as follows: one species of *Noteriades*, one species of *Radoszkowskiana*, three species of *Coelioxys*, and 109 species of *Megachile s.l*. The latter genus is represented by species of 57 subgenera that included those recognized by Michener (2007), the fossil species *M. glaesaria* (Engel, 1999), and three recently described taxa by Baker and Engel (2006), Engel and Baker (2006), Engel and Gonzalez (2011), and Gonzalez and Engel (2012) (Appendix S2). For each subgenus of *Megachile s.l*., we included the type species and, to maximize variation, at least one morphologically divergent species from it, or species separated subgenerically but synonymized by Michener (2000, 2007), Gonzalez *et al*. (2010), Gonzalez and Engel (2012), and Gonzalez (2013). About half of the subgenera are represented by one species because they either are monotypic (10 subgenera) or seemed morphologically uniform (20 subgenera). The only three subgenera of *Megachile s.l*. that we were not able to examine are *M*. (*Austrosarus*) Raw, *M*. (*Neochalicodoma*) Pasteels, and *M*. (*Stellenigris*) Meunier. However, Gonzalez and Engel (2012) and Gonzalez (2013) considered the first as synonym of *M*. (*Chrysosarus*) while the second as synonym of *M*. (*Pseudomegachile*), subgenera represented by several species in our analyses. The identity and correct taxonomic placement of *M*. (*Stellenigris*) is a mistery. Michener (2000, 2007) suggested that it might belong to large species of the Group 2 of *Megachile s.l*., but the type specimen of *Stellenigris vandeveldii* Meunier, 1888, is probably lost or perhaps destroyed, along with other insects described by F. Meunier (Engel, 2007).

Most specimens studied are in the Snow Entomological Collection, University of Kansas Natural History Museum, although we borrowed specimens of a few rare species from the following institutions (names of the people who kindly arranged these loans are in parentheses): Academy of Natural Sciences of Drexel University, Philadelphia, PA (D. Otte, J. Weintraub); American Museum of Natural History, New York (J.G. Rozen, Jr.); Bee Biology and Systematics Laboratory, USDA-ARS, Utah State University, Logan, UT (T. Griswold, H. Ikerd); the Natural History Museum, London, UK (D. Notton); Department of Terrestrial Invertebrates, Western Australian Museum, Welshpool (T. Houston); Illinois Natural History Survey, Urbana, Illinois, USA (P. Tinerella); Museum of Comparative Zoology, Harvard University, Cambridge, MA (P. Perkins, R.L. Hawkins); Musée Royal de L’Afrique Centrale, Tervuren (A. Pauly, E. De Coninck); Museum für Naturkunde der Humboldt-Universität, Berlin, Germany (F. Koch, V. Ritcher); North Carolina State University Insect Museum, Raleigh, NC (Rob Blinn); Oxford University Museum of Natural History, Oxford, UK; and United States National Museum of Natural History, Washington, D.C. (D. Furth, B. Harris).

### Morphological data

Morphological terminology generally follows that of Michener (2000, 2007) and Engel (2001), except for ‘torulus’ and ‘interdental lamina’, which we use herein instead of ‘antennal socket’ and ‘cutting edge’. The first term is in broader application across Hymenoptera while the second describes more accurately the laminae between the teeth of the female mandible that characterizes the majority of LC bee species. ‘Cutting edges’ have widely been used in the taxonomic literature of *Megachile s.l*. (*e.g*., Michener, 1962, 2007) but these terms are functionally and structurally ambiguous. They imply that these are the only structures used in cutting leaves and do not inform on their shape nor on their location in the mandible. The absence of interdental laminae in some species of *Megachile s.l*. [*e.g*., *M*. (*Chrysosarus*) Mitchell] that also cut leaves clearly indicates (*e.g*., Zillikens and Steiner, 2004; Torretta *et al*., 2014) that these are not the only mandibular structures involved in leaf cutting. For example, the upper and lower margins of each tooth are sometimes thin and sharp, and they might function as razors even when the interdental laminae are present. Thus, as initially proposed by Pasteels (1965), the term interdental laminae seems more appropriate than cutting edges to describe the laminae between the teeth. Terminology for the mandible, proboscis, and female’s sting apparatus and associated sterna follows Michener and Fraser (1978), Winston (1979), and Packer (2003, 2004), respectively.

#### Data compilation

For the generic-level phylogeny of Megachilini, we conceptualized and scored the majority of character statements from searching on all parts of the body of both male and female sexes, including the labiomaxillary complex, mandible, and genitalia with its associated terga and sterna. We also took and modified some character statements from the cladistic analyses of Roig-Alsina and Michener (1993) and Gonzalez *et al*. (2012). During the conception and formulation of character statements, the following comparative studies and taxonomic revisions were useful as they mentioned or discussed morphological features of taxonomic importance: Michener (1962, 1965, 2000, 2007), Michener and Fraser (1978), Winston (1979), Mitchell (1980), and Roig-Alsina and Michener (1993).

We examined and measured morphological features using Olympus SZ60 and SZX12 stereomicroscopes with an ocular micrometer. We cleared the labiomaxillary complex and genitalia with 10% KOH at room temperature for about 24h. Then, we washed them with 70% ethanol before storing them in glycerin. To document character states, we prepared line illustrations as well as photomicrographs, which we took with a Canon 7D digital camera attached to an Infinity K-2 long-distance microscope lens, and assembled with Zerene Stacker^TM^ software package. We processed final figures with Adobe Photoshop® CC.

We built a data matrix in WinClada (Nixon, 1999) and scored 272 characters (Appendix S3). However, we were not able to code all characters for all species because some taxa are known only from the type specimen and we could not dissect them, and in other cases, they are only known from one sex. Unless we suspected sexual dimorphism, we took characters from the available sex. We only used continuous characters, such as proportions or measurements, when we found distinct gaps in the measured variable among the examined specimens. To avoid duplication, we coded only in the female some characters that are present in both sexes (*e.g*., labiomaxillary complex).

To facilitate further comparisons, we formulated character statements following Sereno (2007), in which the most general locator is positioned first (*e.g*., antennal scape), followed by a variable (*e.g*., length), a variable qualifier (*e.g*., length relative to torulocellar distance), and mutually exclusive character states, the latter following a colon. In some cases, we added a secondary or tertiary locator to clarify the position of the primary locator.

The following are the descriptions of the character statements used in the generic-level analysis of Megachilini. We indicated the original author of a character statement and used the following abbreviations F, OD, PW, Mt, S, and T for flagellomere, median ocellus diameter, one puncture width, mandibular tooth, and metasomal sterna and terga, respectively.

##### Female Head

1. Subantennal area (i*.e*., clypeoantennal distance), length relative to vertical diameter of torulus (Gonzalez *et al*., 2012: char. 4): 0 = short, equal to or shorter than; 1 = long, ≥ 1.2×.
2. Anterior tentorial pit, location (Roig-Alsina and Michener, 1993: char. 2): 0 = at the intersection of subantennal and epistomal sulci; 1 = on epistomal sulcus, below intersection with subantennal sulcus.
3. Anterior tentorial pit, shape: 0 = rounded, about as long as broad; 1 = elongated, about twice as long as broad.
4. Interantennal area (*i.e*., intertorular distance), length relative to torulorbital distance (Gonzalez *et al*., 2012: char. 9): 0 = equal to or shorter than; 1 = greater than.
5. Antenna, scape, length (excluding basal bulb) relative to torulocellar distance: 0 = equal to or shorter than; 1 = long, ≥ 1.2×.
6. Antenna, pedicel, length relative to length of F1 (modified from Gonzalez *et al*., 2012: char. 13): 0 = short, at most as long as; 1 = long, ≥ 1.5×. In character state 1, the pedicel is often about as long as or longer than length of F1 and F2 combined.
7. Antenna, F1, length relative to F2: 0 = 1.5–2.0× longer than; 1 = about as long as; 2 = shorter than.
8. Vertex, integument, with fine, shining longitudinal line from ocelli to its posterior margin: 0 = absent; 1 = present.
9. Paraocular carina (Roig-Alsina and Michener, 1993: char. 4): 0 = absent; 1 = present.
10. Preoccipital carina (Gonzalez *et al*., 2012: char. 18): 0 = absent; 1 = present.
11. Preoccipital carina, dorsal edge of head behind vertex (modified from Gonzalez *et al*., 2012: char. 19): 0 = present; 1 = absent.
12. Ocelloccipital area, length relative to OD (Gonzalez *et al*., 2012: char. 20): 0 = short, 1.0– 3.0×; 1 = long, ≥ 3.1×.
13. Hypostomal area, short transverse carina: 0 = absent; 1 = present. This short carina encloses a small, shiny, depressed area, behind the mandible and is present in the female of *M.* (*Melanosarus*).
14. Hypostomal carina, porterior portion, tooth or strong protuberance: 0 = absent; 1 = present, distinct. In most species, the hypostomal carina gently curves from the base of the mandible (ventral portion) to behind the head (posterior portion), but in some species a distinct tooth or strong protuberance develops where the ventral portion flexes upwards behind the head.
15. Hypostomal carina, ventral portion, orientation relative to margin of mandibular socket (Gonzalez *et al*., 2012: char. 21): 0 = directed to medial margin; 1 = curving towards posterior margin (Griswold and Gonzalez, 2011: fig. 13).
16. Supraclypeal area, lower portion, shape (modified from Gonzalez *et al*., 2012: char. 8): 0 = flat, elevated or modified, not strongly convex in profile; 1 = strongly convex in profile.
17. Clypeus, width relative to mid length: 0 = short, ≥ 3.0×; 1 = long, ≤ 2.8×.
18. Clypeus, basal portion, shape: 0 = flat or convex, not greatly elevated or ornate; 1 = greatly elevated and ornate.
19. Clypeus, disc, shape: 0 = flat or convex, not elevated; 1 = elevated with flat median section.
20. Clypeus, distal margin, degree of projection over labroclypeal articulation (Gonzalez *et al*., 2012: char. 1): 0 = not projected, articulation clearly visible (Fig. 1D; Engel and Gonzalez, 2011: fig. 8); 1 = slightly projected, articulation not visible (Gonzalez and Engel, 2012: fig. 4); 2 = strongly projected, articulation not visible (Eardley, 2012: fig. 43a). In species having character state 2, the strongly projected distal margin makes the clypeus hexagonal in shape, as in *M.* (*Chalicodoma*). The clypeus of *M.* (*Schrottkyapis*) *assumptionis* Schrottky has a bifid median process strongly produced over the labrum (Silveira *et al*., 2002: fig. 11.25); however, the apicolateral margins of the clypeus slightly cover the base of labrum; thus, we coded this species as having character state 1.
21. Clypeus, complete longitudinal median carina: 0 = absent; 1 = present (Pasteels, 1965: fig. 1059).
22. Clypeus, pubescence, density: 0 = sparse throughout, integument visible among setae; 1 = dense throughout, integument not visible among setae; 2 = dense on sides of clypeus, sparse to absent on disc (Eardley, 2013: fig. 66a).
23. Clypeus, disc, abundant, erect, short and partially hooked or wavy setae: 0 = absent; 1 = present (Durante and Abrahamovich, 2006: figs. 1–3; Gonzalez and Griswold, 2013: fig. 5E). These modified setae are associated with the passive collection of pollen from nototribic flowers.
24. Labrum, shape: 0 = rectangular, base as wide as apex, lateral margins parallel to each other (Mitchell, 1980: fig. 48); 1 = subtriangular, base ≥ 1.5× apical width, lateral margins converging apically (Mitchell, 1980: fig. 30).
25. Labrum, disc, pubescence: 0 = absent; 1 = present.
26. Labrum, disc, type and length of setae: 0 = consisting only of long (≥ 1.0× OD), erect setae; 1 = consisting of two types of setae, minute, yellowish, appressed setae, and long (≥ 1.0× OD), erect setae; 2 = consisting only of minute, yellowish, appressed setae.
27. Labrum, midapical or subapical protuberance: 0 = absent; 1 = present.
28. Mandible, length relative to length of compound eye in lateral view: 0 = short, ≤ 0.7×; 1 = long, ≥ 0.9× (Fig. 1D; Engel and Gonzalez, 2011: fig. 8).
29. Mandible, outer surface, median root of outer ridge: 0 = absent; 1 = present, extending towards abductor swelling (Gonzalez and Engel, 2012: fig. 5).
30. Mandible, outer surface, upper root of outer ridge: 0 = absent; 1 = present, extending towards acetabulum and joining acetabular carina (Fig. 2F).
31. Mandible, outer surface, secondary transverse ridge: 0 = absent; 1 = present, distinct. King (1994: fig. 8) recognized and illustrated this ridge, which is dorsal and parallel to the acetabular groove. In some species, such as *M.* (*Litomegachile*) *brevis* Say, the acetabular interspace is elevated, flattened or evenly convex, with a distinct edge delimiting the superior margin of the acetabular groove. However, we coded these species as having character state 0 because this is an edge, not a ridge.
32. Mandible, transverse ridge, basal portion joining acetabular carina: 0 = absent; 1 = present (King, 1994: fig. 8).
33. Mandible, apex, width relative to base in lateral view: 0 = narrow, equal to or narrower than (Engel and Gonzalez, 2011: fig. 8); 1 = broad, ≥ 1.5× (Figs. 2C–J).
34. Mandible, distal margin, axis: 0 = straight or nearly so, not strongly oblique (Figs. 2C–J); 1 = strongly oblique as in *M.* (*Chalicodoma*) and *M*. (*Chalicodomoides*) (Michener, 2007: fig. 84-12d).
35. Mandible, outer surface, apex, type of integument: 0 = smooth and shiny, or nearly so, between punctures (Figs. 2C–J; Gonzalez and Engel, 2012: fig. 33); 1 = microreticulate to finely punctate (Fig. 3A; Engel and Gonzalez, 2011: fig. 24).
36. Mandible, outer surface, apex of acetabular mandibular groove, distinct tuft or brush of long golden setae: 0 = absent; 1= present (Fig. 2F). In some species, such as *M.* (*Paracella*) *semivenustella* Cockerell, another brush is also present at the apex of the outer groove. In species with a well-developed outer premarginal fimbria, such as *M.* (*Hackeriapis*) *ferox* Smith, the apices of the acetabular and outer grooves often appeared as having brushes; however, the setae on these areas are about the same length and density as those on the outer premarginal fimbria. Thus, we coded these species as having character state 0.
37. Mandible, outer premarginal impressed fimbria (Gonzalez *et al*., 2012: char. 39): 0 = reduced or absent (Fig. 2E); 1 = present, distinct (Fig. 3E; Michener and Fraser, 1978: fig. 29).
38. Mandible, outer surface, acetabular interspace, shape: 0 = not conspicuously flattened or depressed, gently curving towards base of mandible (Fig. 3A); 1 = clearly flattened or depressed, such as outer surface of mandible has a distinguishable basal, lateral surface, and a distal, anterior surface (Fig. 1E). Character state 1 is typical of most Group 1 of subgenera of *Megachile s.l*‥
39. Mandible, tooth count: 0 = two; 1 = three; 2 = four to six; 3 = lower distal margin with one or two large teeth, upper portion edentate or nearly so, or with very small teeth (Michener, 2007: fig. 84-12d, e). In some species, the upper distal margin is incised, resulting in a 5 or 6-toothed mandible (*e.g*., Figs. 2D, H), with the upper teeth closer than other teeth. We coded these species as having character state 2.
40. Mandible, Mt1, width relative to basal width of Mt2: 0 = ≤ 1.4× (Figs. 2C–H); 1 = ≥ 1.5× (Fig. 2I).
41. Mandible, third dental interspace, length relative to combined length of first and second interspaces: 0 = short, ≤ 1.5× or absent; 1 = long, about 2.0× (Michener, 2007: fig. 84-11f).
42. Mandible, upper distal margin, shape: 0 = rounded or pointed with apex anteriorly directed; 1 = pointed, subtriangular, and with apex dorsally directed.
43. Mandible, upper tooth, shape: 0 = acute or right angular (Fig. 2I); 1 = rounded or truncate, not incised (Fig. 2G); 2 = rounded or truncate, incised (Fig. 2H).
44. Mandible, upper margin near distal margin, tooth or projection: 0 = absent; 1 = present.
45. Mandible, upper margin near mandibular base, tooth or projection: 0 = absent; 1 = present (Michener, 1965: fig. 664).
46. Mandible, inner surface preapically: 0 = without a distinct fimbrial ridge or carina; 1 = with a distinct fimbrial ridge running somewhat parallel to the mandibular margin (Fig. 3B); the surface between this ridge and the mandibular margin is sloping; 2 = with a distinct fimbrial carina running parallel to the mandibular margin, usually posterior to the bases of teeth and not apically extended into a lamina; the surface formed between this carina and the mandibular margin somewhat perpendicular (Figs. 3C, D, F); 3 = with a distinct lamina projecting beyond bases of upper teeth (Figs. 2E, 3H).
47. Mandible, second interspace, interdental lamina: 0 = absent; 1 = present (Fig. 2H).
48. Mandible, second interspace, type of interdental lamina: 0 = incomplete, not filling interspace (Fig. 3G); 1 = complete, filling interspace.
49. Mandible, second interspace, origin of interdental lamina: 0 = not arising from inferior border of third tooth and thus interpreted as an apical extension of the fimbrial carina (ctenogenic laminae, see results); 1 = arising from the inferior border of third tooth (odontogenic laminae, Figs. 3C–D). In *M*. *assumptionis* and *M*. (*Stelodides*) *euzona* Pérez, a very small laminar projection (not visible in frontal view) arises from the inferior border of Mt3, and thus suggesting an incomplete interdental lamina; however, we coded these species as having character state 0. In *M.* (*Tylomegachile*) *orba* Schrottky and *M.* (*Tylomegachile*) *simplicipes* Friese, the interdental laminae of the second and third interspaces are presumably fused; however, a frontal view of the mandibular margin reveals that these laminae are in different planes. This suggests that the interdental lamina of the second interspace arises from the third tooth and thus we coded these species as having character state 1.
50. Mandible, second interspace, interdental lamina fused with third tooth, thus resulting in a broad, thin tooth with a more or less truncate margin: 0 = absent; 1 = present. Character state 1 is a putative synapomorphy of *M.* (*Amegachile*) (Michener, 2007: fig. 84-11e).
51. Mandible, third interspace, interdental lamina: 0 = absent; 1 = present (Fig. 3G).
52. Mandible, third interspace, type of interdental lamina: 0 = incomplete, not filling interspace; 2 = complete, filling interspace.
53. Mandible, third interspace, origin of interdental lamina: 0 = not arising from inferior border of fourth tooth and thus interpreted as an apical extension of the fimbrial carina; 1= arising from inferior border of fourth tooth. In *M*. *semivenustella*, in addition to a complete interdental lamina, there seems to be a small, incomplete interdental lamina arising from MT4; thus, we coded this species as having both character states.
54. Mandible, inner surface, inner fimbria, length relative to apical mandibular margin (Gonzalez *et al*., 2012: char. 36): 0 = short, restricted to upper margin (Michener and Fraser, 1978: fig. 25); 1 = long, extending across entire margin (Fig. 3F).
55. Mandible, inner surface, secondary fimbria: 0 = absent; 1 = present (Fig. 3F).
56. Mandible, adductor interspace, setae (Gonzalez *et al*., 2012: char. 37): 0 = absent; 1 = present (Figs. 3B, C).
57. Mandible, adductor interspace, length of setae relative to OD: 0 = short, ≤ 0.2×; 1 = long, ≥ 0.4×.
58. Mandible, adductor interspace, longitudinal, impressed line below adductor apical ridge marked with a series of setae: 0 = absent (Fig. 3F); 1 = present.
59. Mandible, strong adductor apical ridge (Gonzalez *et al*., 2012; char. 34): 0 = absent; 1 = present (Fig. 3C).
60. Labium, glossa (in repose), length: 0 = short, not reaching metasoma; 1 = long, reaching metasoma.
61. Labium, prementum, subligular process, shape (modified from Gonzalez *et al*., 2012: char. 48): 0 = elongated, long and narrow, styliform (Winston, 1979: fig. 12f); 1 = broad, apex truncated or nearly so (Winston, 1979: fig. 38); 2 = broad, with pointed apex (Winston, 1979: fig. 28).
62. Labium, first palpomere, length relative to length of second palpomere: 0 = short, ≤ 0.5×; 1 = long, ≥ 0.8×.
63. Labium, first palpomere, length relative to width: 0 = ≤ 3.5×; 1 = ≥ 4.0×.
64. Labium, first palpomere, distinct brush of setae on midbasal concavity (Gonzalez *et al*., 2012: char. 51): 0 = absent; 1 = present (Winston, 1979: fig. 11a).
65. Labium, third palpomere, axis relative to second palpomere: 0 = on same plane; 1 = at an angle.
66. Maxilla, stipes, dististipital process (Roig-Alsina and Michener, 1993: char. 31): 0 = absent or reduced (Winston, 1979: fig. 7a); 1 = present, elongated, almost joining stipital sclerite.
67. Labium, glossa, shape: 0 = not broadened or ligulate; 1 = broadened or ligulate (Michener, 1965: fig. 716).
68. Maxilla, palpomere count, including basal segment (Modified from Gonzalez *et al*., 2012: char. 60): 0 = two or three; 1 = four or five.
69. Maxilla, palpi, setae length relative to palpomere diameter: 0 = short, ≤ 2.0×; 1 = long, ≥ 2.1×.
70. Maxilla, second palpomere, length relative to width: 0 = short, ≤ 1.6×; 1 = long, ≥ 2.0×.
71. Maxilla, third palpomere, length relative to width: 0 = short, ≤ 2.6×; 1 = long, ≥ 3.0×.
72. Maxilla, lacinia, apical setae, length and thickness setae relative to setae on medial margin: 0 = similar in length and thickness; 1 = distinctly longer and thicker.
73. Hypostoma, paramandibular process (Gonzalez *et al*., 2012: char. 22): 0 = short or absent; 1 = present, long (Gonzalez *et al*., 2012: fig. 6).
74. Hypostoma, paramandibular carina, shape and length relative to distance between paramandibular process and hypostomal carina (modified from Gonzalez *et al*., 2012: char. 23): 0 = short, half or less; 1 = long, ending at hypostomal carina; 2 = long, not reaching hypostomal carina, usually curving upwards or downwards; 3 = long, reaching posterior component of the hypostomal carina and forming a strong lobe.

**Fig 3.**
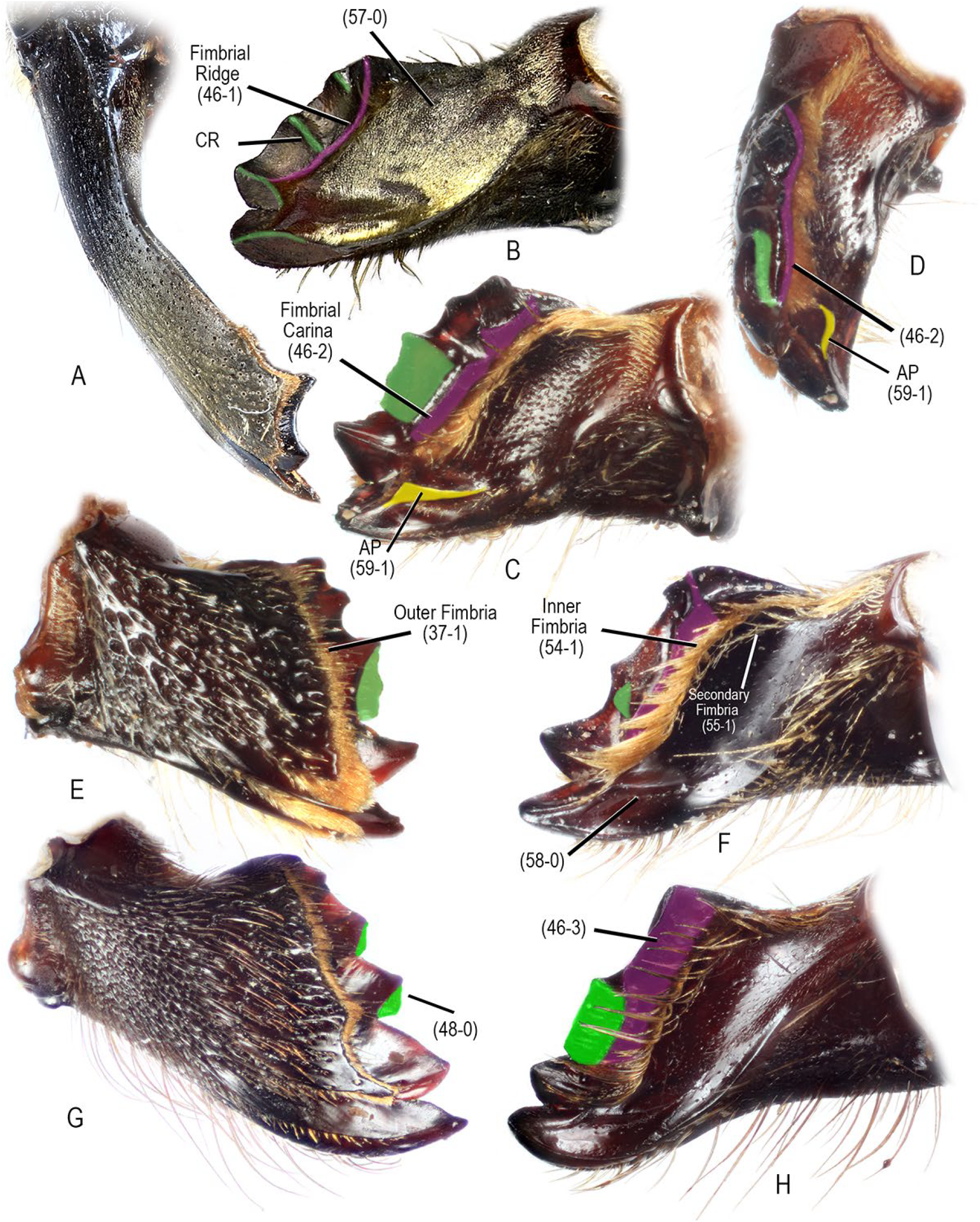
Female mandible of *Megachile s.l*. in outer (A, E, G), frontal (D), and inner views (B, C, F, H). **A.** *M.* (*Callomegachile*) *pluto* (Smith). **B.** *M.* (*Callomegachile*) sp. **C–E**. *M.* (*Chelostomoda*) *spissula* Cockerell. **F**. *M*. (*Rhyssomegachile*) *simillima* Smith. **G.** *M*. (*Creightonella*) *frontalis* (Fabricius). **H.** *M*. (*Pseudocentron*) *pruina* Smith. Interdental laminae highlighted in green (odontogenic) and pink (ctenogenic). Abbreviations: CR = corono-radicular ridge; AP = adductor apical ridge.

##### Mesosoma

75. Pronotal lobe, shape (Gonzalez *et al*., 2012: char. 61): 0 = rounded, without carina or strong lamella; 1 = with strong carina or border; 2 = with conspicuously broad, thin lamella.
76. Omaular carina (Gonzalez *et al*., 2012: char. 65): 0 = absent; 1 = present.
77. Mesepisternum, punctation: 0 = finely or coarsely punctuate, not forming strong rows with distinct shining ridges among them; 1 = coarsely punctuate, forming strong rows with distinct shining ridges among them.
78. Mesoscutum, anterior margin in profile, shape and sculpturing (Gonzalez *et al*., 2012: char. 69): 0 = rounded, without distinctly different surface sculpture; 1 = truncate, perpendicular, or nearly so, shinier and less punctate than dorsal portion.
79. Mesosoma, dorsum, yellow or reddish maculations: 0 = absent; 1 = present.
80. Mesoscutum, disc, length and density of setae: 0 = consisting only of long setae (≥ 3.0– 4.0× OD), integument barely visible; 1 = consisting only of very short setae (≤ 0.5× OD), integument sparsely covered to almost bare; 2 = consisting only of short setae (1.5–2.0× OD), integument visible or partially obscured among setae; 3 = consisting of two types of setae, minute, yellowish, appressed setae, and erect longer setae (2.0× OD); 4 = consisting of semierect or appressed yellowish tomentum uniformly covering the integument.
81. Mesoscutum, notalus line, fascia: 0 = absent; 1 = present.
82. Mesoscutum, parapsidal line, length relative to length of tegula in dorsal view (Gonzalez *et al*., 2012: char. 72): 0 = long, ≥ 0.4×; 1 = short, ≤ 0.3× or absent.
83. Mesoscutum, disc, punctation, density and size: 0 = finely and closely (≤ 1.0–2.0× PW) punctate, punctures (≤ 0.2× OD) not in row; 1 = coarsely and densely punctate, punctures (≥ 0.5× OD) arranged in rows, thus giving a striate or wrinkled appearance (Engel and Gonzalez, 2011: fig. 37); 2 = coarsely and densely punctate, punctures (≥ 0.5× OD) not arranged in rows.
84. Mesoscutal-mesoscutellar suture, white fascia: 0 = absent; 1 = present.
85. Preaxilla (below posterolateral angle of mesoscutum), incline and pubescence (Gonzalez *et al*., 2012: char. 73): 0 = sloping, with setae as long as those on adjacent sclerites (Gonzalez *et al.*, 2012: fig. 10); 1 = vertical, usually nearly asetose (Gonzalez *et al.*, 2012: fig. 11).
86. Axilla, posterior margin, shape (modified from Gonzalez *et al*., 2012: char. 74): 0 = rounded, not projected in acute angle or spine; 1 = weakly projected, not reaching posterior transverse tangent of mesoscutellum; 2 = strongly projected into acute angle or spine, surpassing posterior transverse tangent of mesoscutellum (Michener, 2007: fig. 84-6a).
87. Axilla, lateral surface, shape: 0 = not depressed; 1 = depressed, partially or entirely hidden by dorsal surface.
88. Axilla, lateral surface, sculpturing and pubescence: 0 = similarly punctate and setose as on its dorsal surface; 1 = smooth and shiny, asetose; 2 = micropunctate to strongly imbricate on at least its ventral half, dull, asetose or with sparse setae.
89. Axillar fossa, depth: 0 = shallow, surface behind it subhorizontal, without a high mesoscutellar crest between it and metanotum (Fig. 4A); 1 = deep, its posterior surface usually ascending to strong mesoscutellar crest between fossa and metanotum (Fig. 4B).
90. Mesoscutellum, shape in profile: 0 = flat or convex, forming relatively uninterrupted surface with metanotum, thus without a distinct posterior surface; 1 = elevated from metanotum, with a distinct posterior surface.
91. Metanotal pit: 0 = absent; 1 = present, distinct (Fig. 4B).
92. Metanotum, sublateral length relative to midlength: 0 = about as long as; 1 = narrower than.
93. Metanotum, degree of visibility given by mesoscutellum in dorsal view (modified from Roig-Alsina and Michener, 1993: char. 74): 0 = entirely or partially hidden; 1 = fully exposed (Fig. 4C).
94. Metanotum, median tubercle or spine (Michener, 1996: char. 7): 0 = absent; 1 = present (Michener, 2007: fig. 83-1).
95. Propodeal triangle (= metapostnotum), pubescence (Roig-Alsina and Michener, 1993: char. 79): 0 = present; 1 = absent.
96. Propodeum, shape in profile (Roig-Alsina and Michener, 1993: char. 73): 0 = largely vertical; 1 = entirely slanting or with slanting dorsal portion rounding onto vertical portion.
97. Propodeal pit, shape (Gonzalez *et al*., 2012: char. 87): 0 = rounded or elongate, but not linear; 1 = linear.
98. Legs, color: 0 = dark brown to black, concolor with remaining areas of mesosoma; 1 = reddish or orange, contrasting with dark brown to black mesosoma.
99. Metatibia, outer surface, strong tubercles or spicules that do not end in setae or bristles (Gonzalez *et al*., 2012: char. 102): 0 = absent; 1 = present (Michener, 2007: fig. 80-3b).
100. Mesotibia, outer surface, apically with acute angle and distinct notch anteriorly (Gonzalez *et al.*, 2012: char. 92): 0 = absent; 1 = present (Gonzalez *et al.*, 2012: fig. 14).
101. Mesotibia, outer surface, long, acute medial spine on apical margin: 0 = absent; 1 = present (Fig. 4D).
102. Mesotibia, outer surface, distinct longitudinal carina on apical one-fourth: 0 = absent; 1 = present (Fig. 4E). This carina joins the distal margin of the tibia, sometimes in a sharp angle, and thus appearing as a spine [*e.g*., *M.* (*Amegachile*) *bituberculata* Ritsema]; however, there is always a concave, bare area posterior to this carina, which is absent in taxa that possess a true spine. In some species, this carina and the distal margin of tibia form a distinct spatulate or spoon-like process, easily visible in posterior view. This carina is apically notched in *M*. (*Melanosarus*).
103. Mesotibia, outer surface, area behind longitudinal carina, setae: 0 = present; 1 = absent.
104. Mesotibia, outer surface, posterodistal margin projected into a distinct spine: 0 = absent; 1 = present (Gonzalez and Engel, 2012: fig. 7). In *M.* (*Lophanthedon*) *dimidiata* (Smith) the projection is small and sometimes absent. We coded this species as having character state 0.
105. Mesotibia, outer surface, tuft of stiff setae on posterodistal margin: 0 = absent; 1 = present.
106. Metatibia, basitibial plate (Roig-Alsina and Michener, 1993: char. 84): 0 = absent; 1 = present.
107. Metatibia, scopa consisting of uniformly dispersed long setae on outer surface: 0 = absent; 1 = present.
108. Metatibia, spurs, shape: 0 = pointed, straight or gently curving apically; 1 = pointed, straight with apex strongly curved inward; 2 = not pointed, parallel-sided and with apex blunt.
109. Metabasitarsus, length relative to length of tibia: 0 = short, ≤ 0.5×; 1 = long, ≥ 0.8×.
110. Metabasitarsus, lengh relative to width (modified from Roig-Alsina and Michener, 1993: char. 90): 0 = ≥ 3.0×; 1 = ≤ 2.8×.
111. Pretarsal claws, shape (Roig-Alsina and Michener, 1993: char. 99): 0 = simple (Fig. 4G– I); 1 = bifurcate (Fig. 4F).
112. Pretarsal claws, one or two basal projections: 0 = absent; 1 = present (Figs. 4H, I).
113. Pretarsal claws, seta count: 0 = one; 1 = two.
114. Pretarsal claws, thickness and length of setae relative to each other: 0 = similar thickness, one of them at least half length of the other (Fig. 4H, I); 1 = one conspicuously shorter and stouter than the other (Fig. 4G).
115. Propretarsus, arolium (Roig-Alsina and Michener, 1993: char. 98): 0 = reduced or absent; 1 = present.
116. Forewing, first submarginal cell, length relative to second as measured on posterior margin (modified from Gonzalez *et al*., 2012: char. 116): 0 = equal to or shorter than; 1 = longer than.
117. Forewing, basal vein (M), location relative to cu-a (modified from Gonzalez *et al*., 2012: char. 118): 0 = posterior to; 1 = confluent with, or basal to.
118. Forewing, second recurrent vein (2m-cu), location relative to second submarginal crossvein (2rs-m) (Michener, 1996: char. 12): 0 = basal to (Michener, 2007: fig. 81-1b); 1= confluent with, or distal to (Michener, 2007: fig. 82-1).
119. Forewing, pterostigma, length relative to width (Gonzalez *et al*., 2012: char. 112): 0 = long, ≥ 2.1×; 1 = short, ≤ 2.0×.
120. Forewing, coloration: 0 = entirely hyaline, yellowish, or dusky; 1 = apical half dusky, contrasting with hyaline or yellowish basal half; 2 = yellowish wing base with dusky costal margin.
121. Hind wing, second abscissa of vein M+Cu, length relative to length of cu-v (Gonzalez *et al*., 2012: char. 122): 0 = short, ≤ 3.0×; 1 = long, ≥ 3.1×.
122. Hind wing, jugal lobe, length relative to length of vannal lobe (each lobe measured from wing base to apex) (modified from Roig-Alsina and Michener, 1993: char. 105): 0 = ≤ 0.5×; 1 = ≥ 0.6×.

##### Metasoma

123. Metasoma, shape: 0 = strongly convex dorsally, more or less parallel-sided as in *M.* (*Chalicodoma*) and *M*. (*Chalicodomoides*) (Michener, 2007: fig. 84-9); 1 = not parallel-sided, cordate, triangular, and rather flattened as in *M.* (*Megachile*) (Michener, 2007: fig. 84-8); 2 = as in *Coelioxys* (Michener, 2007: fig. 84-2).
124. T1, junction of anterior and dorsal surfaces, shape (Gonzalez *et al*., 2012: char. 125): 0 = rounded; 1 = angled; 2 = carinate.
125. T1, disc in profile and posterior margin in dorsal view, shape (Gonzalez *et al*., 2012: char. 124): 0 = flattened, posterior margin rounded, anterior and dorsal surfaces indistinguishable; 1 = convex, posterior margin straight or nearly so, with distinct anterior and dorsal surfaces.
126. T1, pubescence, length, density, and color relative to those on T2 and T3: 0 = about the same length, density, and/or color, not contrasting notoriously with these terga; 1 = not of the same color, distinctly longer (2.0–3.0×) and denser.
127. T1, dorsal surface, length relative to length of T2 (measured at midline): 0 = ≥ 0.7×; 1 = ≤ 0.6×.
128. T2, laterally with distinct oval velvety patch: 0 = absent; 1 = present (Pasteels and Pasteels 1971: fig. 1). This velvety patch is sometimes present also on T3.
129. T3, deep postgradular groove (Gonzalez *et al*., 2012: char. 126): 0 = absent; 1 = present.
130. T3, mid portion, deep postgradular groove: 0 = absent, clearly visible only laterally; 1 = present.
131. T3, fasciate marginal zones: 0 = absent; 1 = present.
132. T3, well marked premarginal line: 0 = absent; 1 = present.
133. T6, pygidial plate (Roig-Alsina and Michener, 1993: char. 116): 0 = present; 1 = absent.
134. T6, pubescence, color relative to that of T1–T4: 0 = concolorous (black, pale or yellowish); 1 = not concolorous (orange, yellowish, or pale).
135. T6, short (≤ OD), appressed setae: 0 = absent; 1 = present.
136. T6, disc in profile, shape: 0 = straight or slightly concave; 1 = strongly convex, without preapical notch; 2 = strongly convex, with preapical notch.
137. T6, erect setae on disc: 0 = present; 1 = absent.
138. T6, clubbed setae on disc: 0 = absent; 1= present.
139. T6, wide apical hyaline flange (Gonzalez *et al*., 2012: char. 131): 0 = absent; 1 = present (Gonzalez *et al*., 2012: fig. 15).
140. Sternal scopa (Roig-Alsina and Michener, 1993: char. 110): 0 = present; 1 = absent.
141. S1, midapical tooth or spine (Gonzalez *et al*., 2012: char. 137): 0 = absent; 1 = present.
142. S3, apical white fasciae under scopa: 0 = absent; 1 = present.
143. S3, mid portion, apical white fasciae: 0 = absent; 1 = present.
144. S6, length (measured along midline) relative to width (Gonzalez *et al*., 2012: char. 138): 0 = short, as long as or shorter than; 1 = elongated, ≥ 2.0×.
145. S6, shape: 0 = subtriangular or broad basally, not parallel-sided; 1 = somewhat parallel-sided, not subtriangular or broad basally.
146. S6, apodeme, disc between marginal ridge and transapodemal ridge (Gonzalez *et al*., 2012: char. 139): 0 = present (Gonzalez *et al*., 2012: fig. 18; Packer, 2004: fig. 6a, d); 1 = reduced or absent (Gonzalez *et al*., 2012: figs 19, 20; Packer, 2004: fig. 7f).
147. S6, anterior margin between apodemes, depth and shape: 0 = shallow, without U or V-shaped concavity; 1 = deep, with U or V-shaped concavity.
148. S6, anterior margin, deep and narrow medial furrow: 0 = absent; 1 = present (Gonzalez *et al*., 2012: fig. 19).
149. S6, superior lateral margin just below apodemes, distinct swollen border: 0 = absent; 1 = present.
150. S6, lateral surface near lateral ridge, with a strong recurved border or carina: 0 = absent; 1 = present.
151. S6, pregradular area parallel to lateral margin, with a deep invagination: 0 = absent; 1 = present.
152. S6, pregradular area, degree of sclerotization (modified from Gonzalez *et al*., 2012: char. 142): 0 = well sclerotized; 1 = entirely membranous or weakly sclerotized, often easily broken during dissection (Gonzalez *et al*., 2012: fig. 19); 2 = membranous or weakly sclerotized only medially.
153. S6, apex, shape (modified from Gonzalez *et al*., 2012: char. 144): 0 = truncate to broadly rounded; 1 = V-shaped, pointed.
154. S6, distal margin, shape: 0 = simple, not bilobed; 1 = bilobed.
155. S6, setose area, length of area relative to sternal length, as measured it from base of apodemes to apex of sternum: 0 = covering at most apical fourth; 1 = covering about one-third; 2 = covering at least half.
156. S6, setose area, density of setae: 0 = uniformly covered or nearly so; 1 = bare or nearly so.
157. S6, strong preapical border or carina: 0 = absent; 1 = present.
158. S6, fringe of branched setae on or near apical margin: 0 = absent; 1 = present.
159. S6, smooth, bare rim behind apical fringe of branched setae: 0 = absent; 1 = present.
160. S6, bare rim, thickness and shape: 0 = thin, translucent, posteriorly directed; 1 = thick, rolled or abruptly bent dorsally.
161. Sting apparatus, 7^th^ hemitergite, orientation: 0 = vertical (sting apparatus laterally-compressed); 1 = horizontal (sting apparatus dorso-ventrally compressed).
162. Sting apparatus, apex of gonostylus, setal density and length relative to maximum gonostylar width as seen in lateral view (Gonzalez *et al*., 2012: char. 147): 0 = nearly hairless to sparsely covered by short setae (≤ 1.0×); 1 = densely covered by long plumose setae (≥ 1.2×).
163. 7^th^ hemitergite, lamina spiracularis, sculpturing: 0 = smooth and shiny, not sculptured; 1 = weakly to markedly sculptured (Packer, 2003: fig. 2e).
164. 7^th^ hemitergite, lamina spiracularis with a strong protrusion near base of lateral process (Gonzalez *et al*., 2012: char. 150): 0 = absent or reduced; 1 = present (Packer, 2003: fig. 5b).

**Fig 4.**
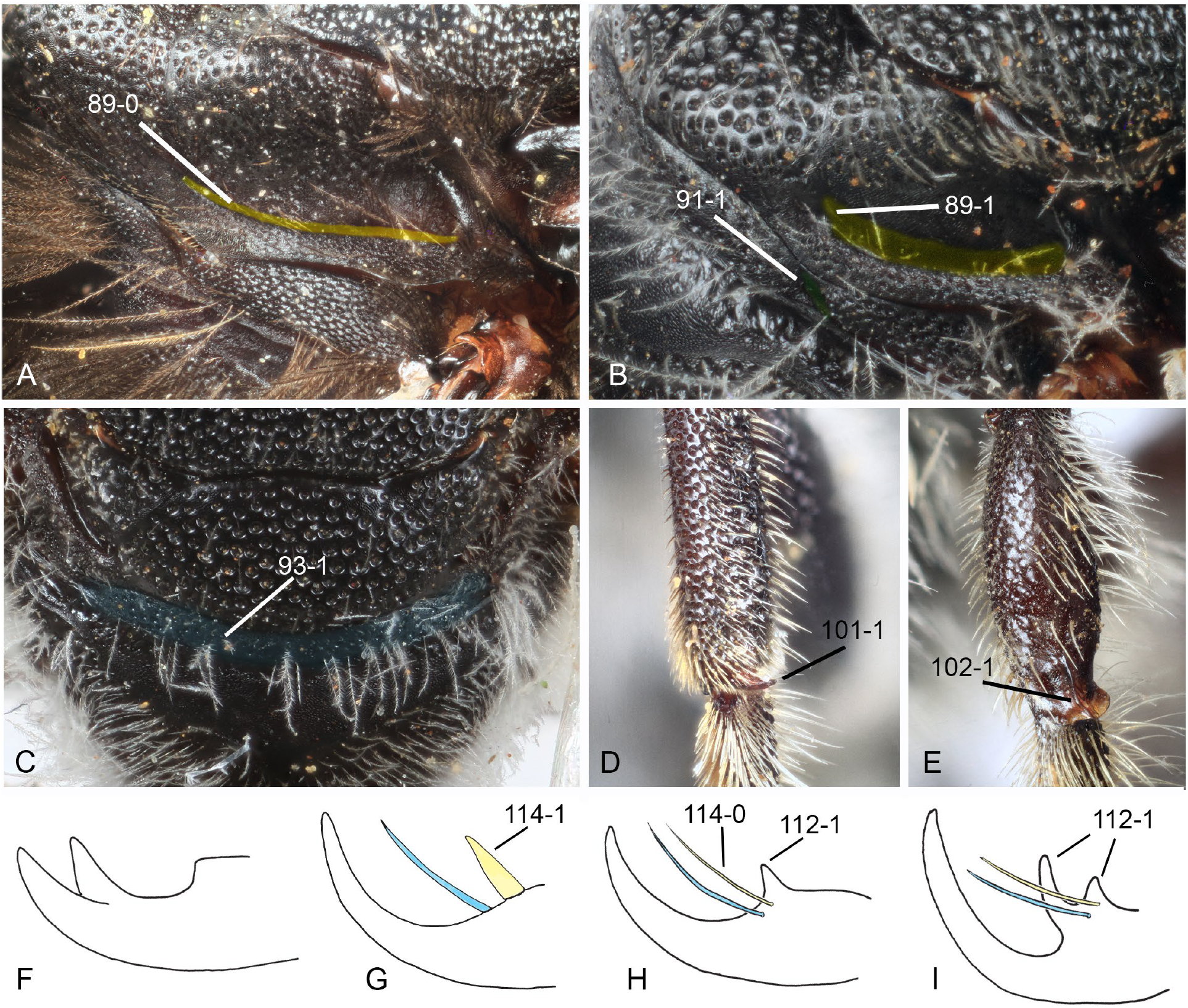
Some male morphological features used in the phylogenetic analysis. **A, B.** Lateral view of axilla. **C.** Dorsal view of mesoscutellum and metanotum. **D, E**. Outer view of apex of mesotibia. **F–I.** Pretarsal claws. *Megachile* (*Melanosarus*) *xylocopoides* Smith (**A**); *M*. (*Stenomegachile*) *dolichosoma* Benoist (**B, C**); *M.* (*Chelostomoides*) *rugifrons* (Smith) (**D**); *M*. (*Megachiloides*) *pascoensis* Mitchell (**E**); *Dioxys productus* (Cresson) (**F**); *M.* (*Acentron*) *albitarsis* Cresson (**G**); *M*. (*Hackeriapis*) *ferox* Smith (**H**); *M*. (*Schizomegachile*) *monstrosa* Smith (**I**).

**Fig. 5.**
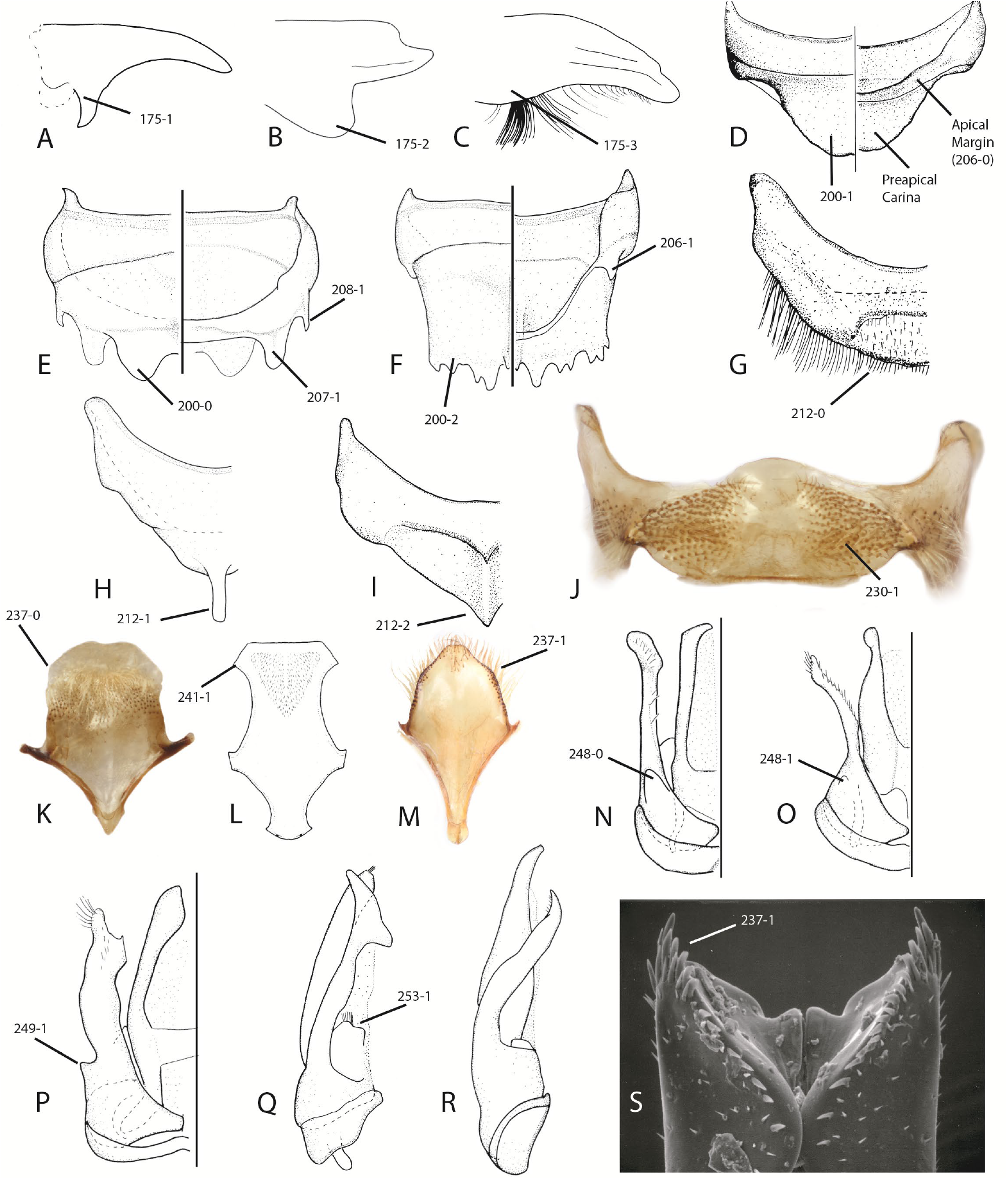
Some male morphological features used in the phylogenetic analysis. **A–C.** Ventral projection of mandible. **D–F.** Dorsal (left half) and ventral (right half) views of sixth tergum. **G– I.** Dorsal view of T7. **J.** Ventral view of sixth sternum. **K–M.** Ventral view of eighth sternum. **N–P.** Dorsal view of genital capsule. **Q, R.** Profile view of genital capsule. **S.** Apex of penis valves. *Megachile* (*Acentron*) *albitarsis* Cresson (**A, L**); *M.* (*Callomegachile*) *biseta* Vachal (**B**); *M*. (*Maximegachile*) *maxillosa* Guérin-Méneville (**C**); *M*. (*Argyropile*) *longuisetosa* Gonzalez and Griswold (**D, G**); *M*. (*Grosapis*) *cockerelli* (**E, H, R**); *M*. (*Creightonella*) *cognata* Smith (**F, I**); *M*. (*Zonomegachile*) *moderata* Smith (**J, K**); *M*. (*Largella*) *donbakeri* Gonzalez and Engel (**M**); *M*. (*Austromegachile*) *montezuma* Cresson (**N**); *M.* (*M.*) *centuncularis* (Linnaeus) (**O**); *M.* (*Moureapis*) *maculata* Smith (**P**); *M.* (*Chalicodoma*) *parietina* (Geoffroy) (**Q**); *M.* (*Chalicodoma*) *sicula* (Rossi) (**S**).

##### Male Head

165. Clypeus, pubescence, density: 0 = sparse throughout, integument visible among setae; 1 = dense throughout, integument not visible among setae (Gonzalez *et al*., 2018: fig. 5C); 2 = basal half with sparse setae (integument visible) or mostly bare, distal half densely covered by setae (integument not visible) (Gonzalez *et al*., 2018: fig. 5D).
166. Clypeus, coloration: 0 = dark brown to black; 1 = yellow.
167. Antenna, F1, length relative to length of F2: 0 = 1.5–2.0×; 1 = about as long as; 2 = shorter than.
168. Antenna, F5–F10, shape: 0 = cylindrical, flattened, or crenulate; 1 = deeply concave on one side.
169. Antenna, F11, shape: 0 = cylindrical; 1 = compressed or flattened (Engel and Baker, 2006: fig. 5).
170. Hypostomal area, with a concavity or protuberance: 0 = absent; 1 = present (Gonzalez *et al*., 2018: fig. 5E).
171. Gena, with an oblique, low, smooth, and shiny carina bordered with a dense row of white branched setae: 0 = absent; 1 = present.
172. Mandible, tooth count: 0 = two; 1 = three; 2 = four; 3 = distal margin of mandible with basal two-thirds edentate or nearly so, at most, one or two very small teeth as in *M.* (*Chalicodoma*).
173. Mandible, upper distal margin, shape and size relative to length and width as remaining teeth: 0 = rounded or pointed, similar length and width; 1 = triangular, conspicuously broader and longer than.
174. Mandible, inferior border, with tooth, process, or projection (Gonzalez *et al*., 2012: char. 156): 0 = absent; 1 = present (Gonzalez *et al*., 2018: fig. 5F).
175. Mandible, inferior process, shape and orientation: 0 = broad, subtriangular, posteriorly-directed, on basal third of inferior border (Gonzalez *et al*., 2018: fig. 5E); 1 = slender, posteriorly-directed (Fig. 5A; Praz, 2017: fig. 8); 2 = broad, small or large, anteriorly-directed, on basal two-thirds of inferior border (Fig. 5B); 3 = broad, with a very dense brush of stiff branched setae (Fig. 5C; Gonzalez and Engel, 2012: fig. 42); 4 = with a small angle midapically (Durante and Cabrera, 2009: fig. 6).
176. Mandible, inner surface, degree of concavity: 0 = weak; 1 = strong.

**Fig 6.**
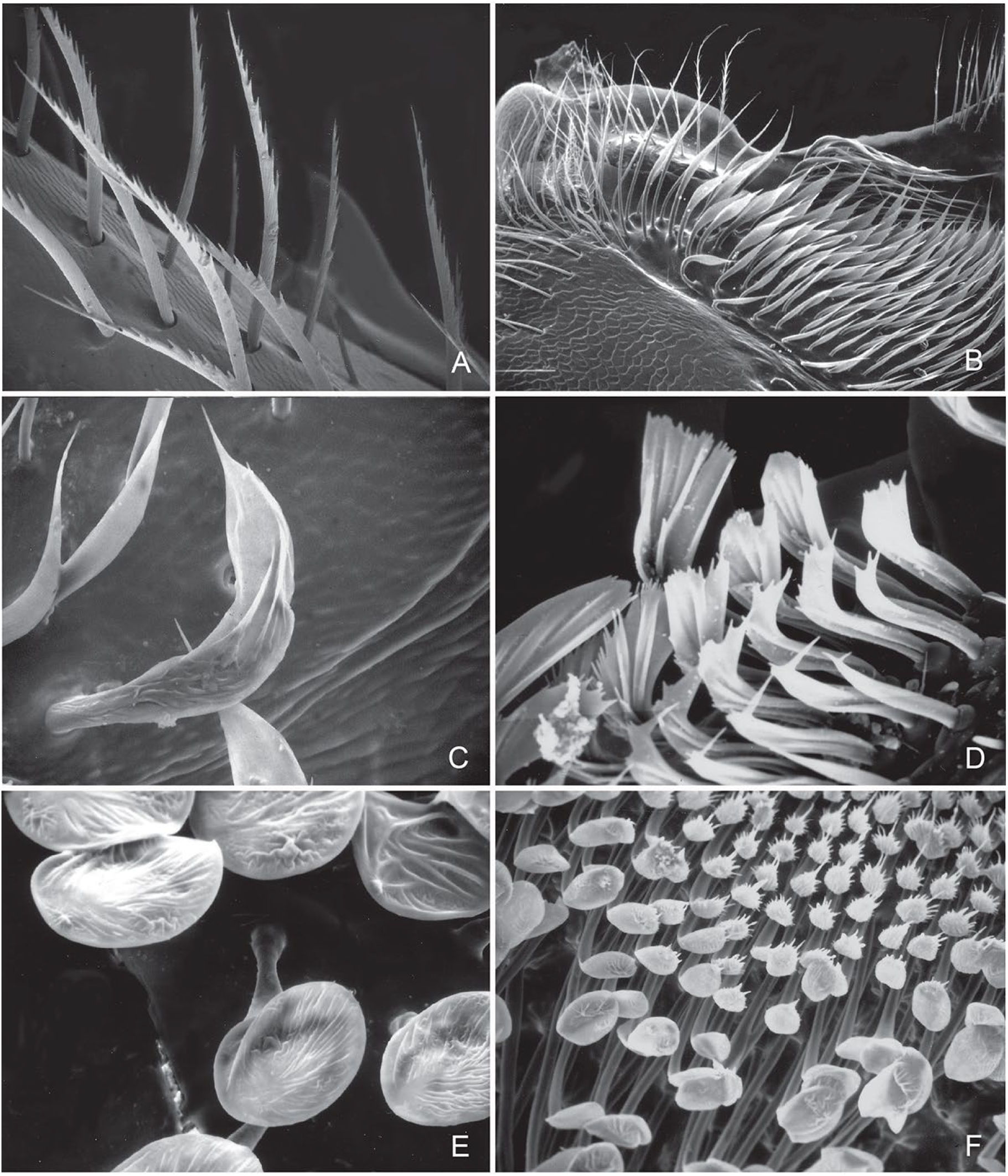
Examples of the types of setae found on the male S4–S6 of *Megachile s.l*. **A.** Branched, unmodified, *Megachile* (*Acentron*) *albitarsis* Cresson, S4. **B.** Acuminate, S4, *M.* (*Megachile*) *centuncularis* (Linnaeus). **C.** Acuminate, S6, *M.* (*Chalicodoma*) *sicula* (Rossi). **D.** Fan-shaped, S6, *M.* (*Chelostomoides*) *exilis* Cresson. **E.** Capitate-spatulate, S5, *M.* (*Chelostomoides*) *rugifrons* (Smith). **F.** Capitate-spatulate, S5, *M.* (*Xanthosarus*) *fortis* Cresson.

##### Mesosoma

177. Procoxal spine (Gonzalez *et al*., 2012: char. 157): 0 = absent; 1 = present (Gonzalez and Engel, 2012: fig. 43).
178. Procoxal spine, shape and length relative to OD: 0 = short (≤ 1.5×), pointed or somewhat parallel-sided; 1 = long (≥ 2.0×), not parallel-sided; 2 = long (≥ 2.0), parallel-sided or nearly so.
179. Procoxal spine, ventral surface, pubescence: 0 = very sparse to nearly asetose, integument clearly visible; 1 = densely covered with branched setae, integument barely visible among setae.
180. Procoxa, disc, pubescence: 0 = uniformly covered with branched setae, integument barely visible among setae; 1 = asetose or nearly so, integument clearly visible.
181. Procoxa, tuft of stiff ferruginous setae: 0 = absent; 1 = present.
182. Protrochanter, inferior margin apically produced: 0 = absent; 1 = present.
183. Profemur, shape and color: 0 = not strongly compressed, same color of femora of remaining legs; 1 = antero-posteriorly strongly compressed, bright yellow or pale, contrasting with color of femora of remaining legs.
184. Protibia, shape and length relative to width: 0 = not enlarged or swollen, ≥ 3.0×; 1 = distinctively swollen, enlarged, ≤ 2.8×.
185. Protarsi, shape and color (Gonzalez *et al*., 2012: char. 158): 0 = not enlarged or excavated, without conspicuous dark spots on inner surface; 1 = slightly or distinctly enlarged or excavated, often with conspicuous dark spots on inner surface.
186. Protarsi, degree of excavation and color: 0 = slightly excavated, with dark spots on inner surface, usually of the same color of tarsi of remaining legs (Michener, 2007: fig. 84-19b); 1 = strongly modified, distinctively enlarged or excavated, inner surface with dark spots, bright yellow or pale, contrasting with tarsi of remaining legs (Michener, 2007: fig. 84-19a).
187. Protarsi, basal tarsomere, shape: 0 = not in the shape of a concave, long, distally directed lobe; 1 = forming a distinctly concave, long, distally directed lobe.
188. Mesocoxa, inner surface, small tooth or protuberance: 0 = absent, 1 = present.
189. Mesotibia, inner surface, tooth or protuberance: 0 = absent, 1 = present.
190. Mesotibial spur: 0 = present; 1 = absent.
191. Mesotibial spur, length relative to apical width of metatibia: 0 = at least as long as; 1 = much shorter than.
192. Mesotibial spur, articulation with metatibia: 0 = free, not fused to tibia; 1 = fused to tibia.
193. Mesobasitarsus, length relative to width: 0 = long, ≥ 2.5×; 1 = short, ≤ 2.0×.
194. Metafemur, posterior surface, patch of microtrichia (metafemoral keirotrichia): 0 = absent; 1 = present (Gonzalez *et al*., 2018: fig. 5I). We coded *M. laeta* as having character state 1 even though this structure is very small.
195. Metatibia, inner spur: 0 = present; 1 = absent.
196. Metabasitarsus, length relative to width: 0 = long, ≥ 2.3×; 1 = short, ≤ 2.0×.
197. Propretarsus, arolium: 0 = present (Baker and Engel, 2006: fig. 5); 1 = reduced or absent.

##### Metasoma

198. T3, marginal zone, color relative to tergal disc: 0 = concolorous; 1 = not concolorous, semi-translucent to translucent.
199. T6, transverse preapical carina (Gonzalez *et al*., 2012: char. 162): 0 = absent; 1 = present.
200. T6, transverse preapical carina, shape: 0 = strong, medially emarginate, not toothed or denticulate (Fig. 5E); 1 = strong, entire or nearly so (Fig. 5D); 2 = strong, toothed or denticulate, with or without a median emargination (Fig. 5F); 3 = weak, little projected in profile, entire or nearly so (Baker and Engel, 2006: fig. 2).
201. T6, preapical carina divided in two or more dorsal processes, and a pair of ventral processes: 0 = absent; 1 = present (Michener, 2007: fig. 84-7).
202. T6, above preapical carina, with strong longitudinal median ridge or protuberance: 0 = absent; 1 = present.
203. T6, above preapical carina, with distinct median concavity: 0 = absent; 1 = present.
204. T6, region of preapical carina, shape: 0 = not swollen or bulbous; 1 = swollen or bulbous, except medially.
205. T6, dorsal surface, density and length of setae relative to OD: 0 = densely covered (integument not visible) by long (2.0–3.0×) setae; 1 = bare or sparsely covered (integument visible) by long (2.0–3.0×) or short (≤ 1.0×) setae; 2 = densely covered by short (≤ 1.0×), appressed, branched setae.
206. T6, apical margin, with lateral spine or tooth: 0 = absent (Fig. 5D); 1 = present (Fig. 5F).
207. T6, apical margin, with submedian spine or tooth: 0 = absent; 1 = present (Fig. 5E).
208. T6, apical margin, size of lateral spine or tooth: 0 = large; 1 = small (Fig. 5E).
209. T6, apical margin, submedian spine or tooth, size relative to size of lateral spine or tooth: 0 = similar; 1 = conspicuously longer and broader than.
210. T7, degree of visibility in dorsal view and orientation (Gonzalez *et al*., 2012: char. 164): 0 = exposed, posteriorly directed; 1 = hidden, and/or anteriorly or ventrally directed.
211. T7, strongly carinate gradulus: 0 = absent; 1 = present (Gonzalez *et al*., 2018: fig. 6B).
212. T7, transverse carina, shape: 0 = rounded, truncate, or emarginate (Fig. 5G); 1 = long, acute, spiniform (Fig. 5H); 2 = angular (Fig. 5I).
213. T7, with distinct, strong longitudinal median ridge: 0 = absent; 1 = present.
214. T7, apical margin, shape: 0 = straight or nearly so, not emarginate or strongly projecting; 1 = with a small median tooth; 2 = deeply and broadly emarginate, forming two prominent teeth (Engel and Baker, 2006: fig. 6); 3 = little projected medially, with small, submedian tooth; 4 = little projected medially, without submedian tooth.
215. T7, pygidial plate (Roig-Alsina and Michener, 1993: char. 118): 0 = present; 1 = absent.
216. Sterna, number of fully exposed sclerites (modified from Gonzalez *et al*., 2012: char. 168): 0 = three; 1 = four; 2 = five or six; 3 = two.
217. S1, midapical spine: 0 = absent; 1 = present.
218. S5, width relative to length (Gonzalez *et al*., 2012: char. 175): 0 = ≤ 2.0×; 1 = ≥ 2.1×.
219. S5, gradulus, degree of sclerotization and definition: 0 = weak, barely distinguishable; 1 = strong, indicated by a well-defined transverse line or border.
220. S5, postgradular area laterally, with setose, sclerotized surface: 0 = absent; 1 = present.
221. S5, apical margin, shape: 0 = straight or nearly so; 1 = deeply or shallowly concave.
222. S5, with short, well-sclerotized midapical process: 0 = absent; 1 = present (Mitchell, 1980: fig. 42).
223. S5, postgradular area, size of setose area relative to width of sternum (Gonzalez *et al*., 2012: char. 177): 0 = large, ≥ 0.6×; 1 = small, ≤ 0.5×.
224. S5, postgradular area, type of setae (modified from Gonzalez *et al*., 2012: char. 176): 0 = simple, branched or plumose (Fig. 6A); 1 = lanceolate, ovate-acuminate (Figs. 6B, C); 2 = capitate or spatulate (Figs. 6E, F); 3 = fan-shaped (Fig. 6D). Sometimes we found more than one type of setae, and thus we coded the most abundant type.
225. S5, postgradular area, with broad, asetose, and weakly sclerotized area above pubescence: 0 = absent; 1 = present (Gonzalez *et al*., 2018: fig. 26D).
226. S5, apicolateral margin, type and length of setae relative to those on postgradular area: 0= asetose or with short setae of similar length; 1 = with simple or branched longer (2.0– 3.0×) setae.
227. S5, midapical margin, with dense tuft of stiff, thickened, simple setae: 0 = absent; 1 = present.
228. S6, width relative to length (Gonzalez *et al*., 2012: char. 183): 0 = ≤ 2.0×; 1 = ≥ 2.1×. Because the midapical margin of S6 is highly variable, we measured the length of S6 on its lateral margin, from the base of the apodeme to the apical margin of the sternum.
229. S6, degree of sclerotization (Gonzalez *et al*., 2012: char. 182): 0 = well-sclerotized; 1 = weakly sclerotized to membranous.
230. S6, postgradular area, pubescence (Gonzalez *et al*., 2012: char. 184): 0 = absent or very sparse (integument clearly visible among setae), without forming distinct patches; 1 = dense, forming distinct patches (Fig. 5J). In *Trichothurgus wagenknechti* (Moure), a mediolongitudinal bare area divides the discal pubescence of S3–S6. Thus, the resulting patches of setae on S6 might not be homologous to those found in other megachiline bees. However, we coded this species as having character-state 1.
231. S6, bare area between setal patches, width relative to one patch width: 0 = wide, ≥ 1.0 ×; 1 = small, ≤ 0.5×.
232. S6, postgradular area, type of setae (Modified from Gonzalez *et al*., 2012: char. 185): 0 = unmodified, simple or branched (Fig. 6A); 1 = modified, lanceolate, ovate-acuminate (Figs. 6B, C), capitate, spatulate (Figs. 6E, F), or fan-shaped (Fig. 6D).
233. S7, degree of sclerotization (modified from Gonzalez *et al*., 2012: char. 186): 0 = entirely well-sclerotized, usually setose; 1 = weakly sclerotized, membranous, frequently asetose.
234. S8, length relative to width: 0 = ≤ 2.5×; 1 = ≥ 2.6×.
235. S8, spiculum, shape: 0 = pointed or broadly rounded (Michener, 2007: figs. 77-1b, 80-4d); 1 = subrectangular; 2 = as an elongated, narrow process (Michener, 2007: fig. 82-2i); 3 = as a short process with an expanded apex (Gonzalez and Griswold, 2013: fig. 508).
236. S8, lateral apodemes: 0 = absent or weakly sclerotized (Michener, 2007: fig. 80-4d); 1 = distinct (Michener, 2007: fig. 82-2b).
237. S8, lateral margins, setae: 0 = absent (Figs. 5K, L); 1 = present, forming a distinct fringe (Fig. 5M).
238. S8, apex, length relative to sternal length, as measured from lateral apodemes to distal margin: 0 = short, about 1/4 (Michener, 2007: fig. 80-4d); 1 = long, about half.
239. S8, apex, width relative to width of spiculum: 0 = wider; 1 = about as wide as or narrower than.
240. S8, apex, shape: 0 = broadly or narrowly rounded; 1 = subrectangular (Fig. 5L).
241. S8, apex, shape: 0 = not expanded; 1 = expanded (Fig. 5L).
242. S8, distal margin, shape: 0 = entire, straight, broadly rounded or pointed (Michener, 2007: fig. 84-4b); 1 = entire, with a small midapical projection (Michener, 2007: fig. 77-1b); 2 = bilobed (Michener, 2007: fig. 82-2b).
243. Genital capsule, length relative to width: 0 = short, about as long as; 1 = elongated, longer than. We measured maximum total length from base of gonobase to apex of penis valves or gonostylus and maximum width at the base of gonobase.
244. Genital foramen, orientation: 0 = anteriorly directed or nearly so (Michener, 2007: fig. 80-4c); 1 = ventrally directed (Michener, 2007: fig. 77-1a).
245. Gonobase (modified from Roig-Alsina and Michener, 1993: char. 122): 0 = present, distinguishable; 1 = reduced or absent.
246. Articulation between gonostylus and gonocoxite (modified Roig-Alsina & and Michener, 1993: char. 125): 0 = distinct, at least ventrally; 1 = fused, thus forming an unsegmented appendage.
247. Gonocoxite, dorsal lobe: 0 = absent; 1 = present (Gonzalez *et al*., 2018: fig. 6E).
248. Gonocoxite, dorsal lobe, shape: 0 = large, strong, digitiform (Fig. 5N; Engel and Baker, 2006: fig. 11); 1 = small, acute (Fig. 5O).
249. Gonocoxite, small sublateral lobe: 0 = absent; 1 = present (Fig. 5P).
250. Volsella (modified from Roig-Alsina and Michener, 1993: char. 126): 0 = absent; 1 = present.
251. Articulation between volsella and gonocoxite: 0 = fused; 1 = articulated, distinguishable as a separated sclerite (Michener, 2007: fig. 77-1a).
252. Volsella, apex, shape: 0 = rounded or pointed; 1 = distinctly notched or bilobed, thus suggesting a medial digitus and a lateral cuspis (Gonzalez and Engel, 2012: fig. 28).
253. Volsella with setae on distal margin: 0 = absent; 1 = present (Fig. 5Q).
254. Gonostylus, length relative to gonocoxite: 0 = equal or shorter than; 1 = ≥ 2.0×.
255. Gonostylus, length relative to penis valves in ventral view (modified from Gonzalez *et al*., 2012: char. 196): 0 = subequal to; 1 = longer than; 2 = shorter than.
256. Gonostylus, shape in lateral view: 0 = curved or arched; 1 = straight or nearly so.
257. Gonostylus, width in lateral view: 0 = not conspicuously narrow, widest at midlength or at apex (Fig. 5Q); 1 = very narrow, about the same width across its entire length (Fig. 5R).
258. Gonostylus, shape in cross section: 0 = not flattened; 1 = flattened.
259. Gonostylus, apex, orientation in dorsal view: 0 = laterally directed; 1 = medially directed; 2 = posteriorly directed.
260. Gonostylus, apex, shape: 0 = not expanded; 1 = clearly expanded.
261. Gonostylus, apical lobes: 0 = absent; 1 = present.
262. Gonostylus, apical lobes, types: 0 = one lateral and one medial (Gonzalez *et al*., 2018: fig. 6D); 1 = one dorsal and one ventral. The gonostylus of *M.* (*Xanthosarus*) *lagopoda* (Linnaeus) has three apical lobes; one on each medial, ventral, and dorsal surfaces. We coded this species as having character states 1 and 2.
263. Gonostylus, medial apical lobe, size: 0 = small, barely indicated; 1 = large and conspicuous (Gonzalez *et al*., 2018: fig. 6D).
264. Gonostylus, apex with large, deep concavity between dorsal and medial lobes: 0 = absent; 1 = present (Gonzalez *et al*., 2018: fig. 6D).
265. Gonostylus, medial surface, pubescence: 0 = absent; 1 = present.
266. Gonostylus, medial surface, length of setae relative to maximum apical gonostylar width: 0 = short, ≤ 2.0×; 1 = long, ≥ 2.1× (Gonzalez and Engel, 2012: fig. 28).
267. Penis valve, apodemes, length relative to their visibility outside genital capsule: 0 = short, not visible; 1 = long, visible as they project through genital foramen (Michener, 2007: fig. 82-2d).
268. Penis valve, shape in dorsal view: 0 = distinctly curved or arched; 1 = straight or nearly so.
269. Penis valve, basal shape: 0 = not enlarged or protuberant; 1 = distinctly expanded.
270. Penis valve, lateral margin, shape: 0 = not enlarged or protuberant; 1 = distinctly enlarged or protuberant.
271. Penis valve, apical shape in ventral view: 0 = straight or nearly so; 1 = distinctly curved or arched inward; 2 = distinctly curved or arch outward (as in *Aztecanthidium*).
272. Penis valve, apex with row of thick, spine-like setae: 0 = absent; 1 = present (Fig. 5S).

#### Data characterization

Both morphological datasets used in the phylogenetic analyses consisted of characters scored from all tagmata of the adult body in both sexes (Fig. 7). However, more characters were scored from the female than from the male (Chi Square test, Megachilidae: *X*^*2*^_[1]_ = 52.02, *P* ≤ 0.001, *n* = 200; Megachilini: *X*^*2*^ _[1]_= 5.14, *P* = 0.023, *n* = 252), even after excluding some characters (∼10%) that are present in both sexes but were scored only in the female to avoid duplication. The number of characters was similar among tagmata in the Megachilidae dataset (*X*^*2*^_[2]_ = 0.13, *P* = 0.937, *n* = 200), but more characters are from the metasoma than from other tagmata in the Megachilini dataset (*X*^*2*^_[2]_ = 13.07, *P* = 0.001, *n* = 272). In both datasets, the number of characters within each sex was significantly different among tagmata (Megachilidae: ♀: *X* ^*2*^_[2]_= 13.60, *P* = 0.001, *n* = 151; ♂: *X*^*2*^ _[2]_ = 51.47, *P* ≤ 0.001, *n* = 49; Megachilini, ♀: *X*^*2*^_[2]_ = 10.59, *P* = 0.005, *n* = 164; ♂: *X*^*2*^ _[2]_= 64.5, *P* < 0.001, *n* = 108). In the female, most characters are from the head and mesosoma whereas for the male most characters are from the metasoma.

**Fig 7.**
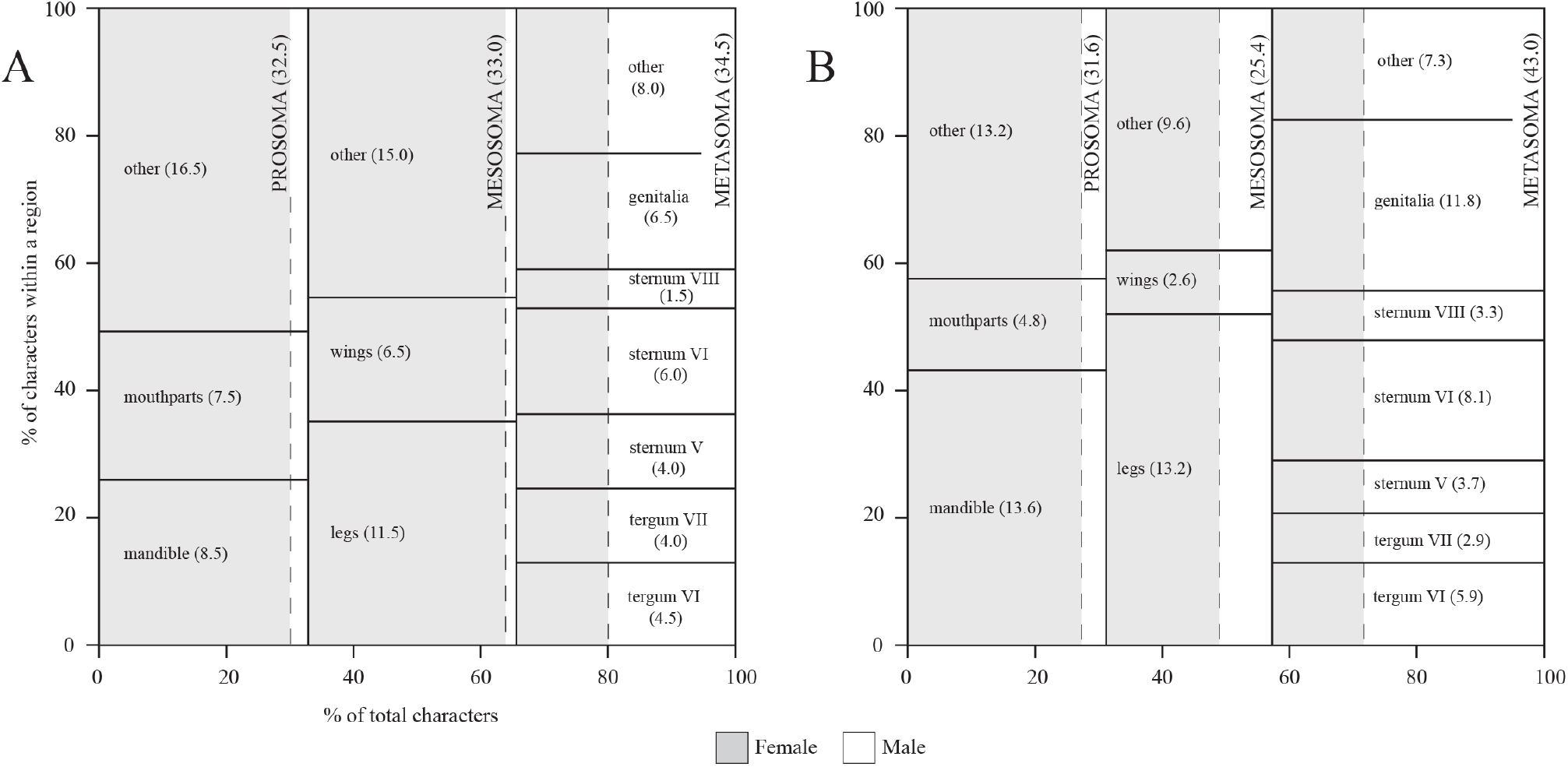
Character distribution maps of the morphological datasets used in the tribal-level phylogeny of Megachilidae (A, *n* = 200 characters) and generic-level phylogeny of Megachilini (B, *n* = 272). The *x*-axis represents the percentage of total characters in each tagma or body region (*e.g*., prosoma) while the *y*-axis represents the percentage of characters of selected structures (*e.g*., mandible) within a tagma. Percentage in parentheses represents contribution to the total number of characters. See Whitlock and Wilson (2013) for further explanation on character distribution maps.

### Molecular data

We used molecular sequences available through GenBank from the following five gene regions generated by Litman *et al*. (2011) and Trunz *et al*. (2016): the protein coding genes elongation factor 1-α (EF), LW-rhodopsin (Opsin), conserved ATPase domanin (CAD), sodium potassium adenosine triphosphatase (NAK), and the ribosomal gene 28S (Appendix S4). We aligned gene fragments using MAFFT ver. 7.305 (Katoh and Standley, 2013), with the secondary structure of 28S accounted for using the Q-INS-I method (Katoh and Toh, 2008). The alignments were then cleaned, frame checked, and concatenated in Mesquite ver. 3.40 (Maddison and Maddison, 2018). Because the gene fragments EF1a, opsin, and CAD have introns (Trunz *et al.*, 2016), we conducted analyses using two approaches. In one analysis, we retained introns, as originally aligned by MAFFT, whereas in the other we removed them and their surrounding variable regions using Gblocks (Castresana, 2000; Talavera and Castresana, 2007) under the less stringent selection options of ‘allow gap positions’ and ‘allow less strict flanking positions’. We also removed the highly variable regions of 28S using Gblocks under these parameters. To find the partition scheme of the molecular data for phylogenetic analysis, we used PartitionFinder ver. 2.1.1 (Lanfear *et al.*, 2016) on both approaches of the concatenated molecular datasets under the ‘greedy’ search algorithm (Lanfear *et al.*, 2012), with unlinked branch-lengths, and Akaike information criterion corrected (AICc) model selection.

### Combined data

DNA sequences are available for many of the species used in the morphological analyses. However, they are not available for many others, particularly those known from the holotype or from a small number of specimens, which in most cases represent the type species of a genus-group name. In those cases, we used available molecular data for closely related species (*i.e*., same subgenus or species group) to those scored in the morphological analysis. We chose to use these chimeric taxa for pragmatic reasons, in an attempt to increase the taxonomic representation of our analyses. Although we did not assess the differences in the number of character states between the pair of species combined, we are confident that the anatomical overlap is high because closely related taxa tend to share a high number of morphological features. In addition, because our goal was to explore the relationships among tribes and genera, we pursued and scored morphological characters that might reflect those levels of relationships (*i.e*., morphological features common to a group of genera or subgenera), not characters aimed to reveal relationships among the species within a subgenus. We referred to those chimeric taxa by their genus name, and sometimes subgenus, followed by a combination of the first three letters of both specific epithets in square brackets (Appendix S2). For example, the name for the operational taxonomic unit (OTU) resulting from *Nomada utahensis* and *N. maculata* is referred as *Nomada* [uta×mac] in the combined dataset.

For the tribal-level analysis, six taxa with morphological information [*Anthidioma chalicodomoides* Pasteels, *Gnathanthidium prionognathum* (Mavromooustakis), *Indanthidium crenulaticauda* Michener and Griswold, *Osmia scutellaris* (Morawitz), *Xenoheriades micheneri* Griswold, *Xenostelis polychroma* Baker] did not have closely related species with DNA sequences available and thus we excluded them from the analyses. Six out of the 73 remaining taxa of the original morphological dataset of Gonzalez *et al*. (2012) are Baltic amber fossils and 44 are species with available DNA sequences. Thus, the remaining 24 terminal taxa are chimeric taxa (Appendix S2). The resulting dataset consisted of 73 OTUs, 200 morphological characters, and 5667 aligned nucleotide positions. For the generic-level analysis, the combined dataset consisted of 67 OTUs, 268 morphological characters, and 6981 aligned nucleotide positions. One of the OTUs is a fossil taxon, 34 have available DNA sequences, and the remaining 33 are chimeric taxa. This combined dataset have 45% less the number of taxa used in the morphological analysis because species of many subgenera do not have available molecular data. However, most of these are from a large, well-supported clade that includes the LC bees (see results). Thus, reducing the taxonomic representation of this clade does not significantly affect the overall taxonomic coverage of the different lineages of the tribe. After reducing the number of taxa from the original morphological dataset, four characters became inapplicable and thus we excluded them. To explore other hypotheses of generic-level relationships, we also analyzed a combined datataset that had all taxa used in the morphological analysis, even though many of them lacked molecular data. We referred to these datasets as the reduced (67 OTUs) and full datasets (122 OTUs).

### Phylogenetic analyses

We analyzed the morphological dataset of the generic-level study using maximum parsimony under two weighting schemes. Because Gonzalez *et al*. (2012) already analyzed the morphological dataset of the tribal-level study, we did not reapeat these analyses. For the analyses of the molecular datasets and combined molecular and morphological datasets, we used maximum likehood (ML) and Bayesian inference (BI). The following are the parameters and treatments used:

#### Parsimony

We treated all characters as unordered and nonadditive, and used equal weights (EW) and implied weigths (IW) in Tree Analysis Using New Techology (TNT; Goloboff *et al*., 2003a). The IW analysis downweights characters according to their degree of homoplasy (*i.e*., characters with higher homoplasy have lower weigths) during the heuristic search for most parsimonious hypotheses (Goloboff, 1993). In IW analyses, instead of using random *k*-values to vary the strength of the weigthing function, we explored a range of constant *k-*values calculated for average character fits (*F*) of 50, 54, 58, 62, 66, 70, 74, 78, 82, 86, and 90%. We obtained these *k*-values using the following formula described in Mirande (2009), and Reemer and Ståhls (2012, 2013): *k* = (*F*×*S*)/1-*F*). *S* is a measure of the average homoplasy per character, calculated as the number of observed steps minus the minimum number of steps divided by the number of characters. The number of observed steps is based on the shortest tree found under EW, which for our dataset is 2364. The minimum number of steps is the cumulative number of minimum character state changes for all 272 characters, which amounts to 323. Thus, the value of *S* is: (2364-323)/272 = 7.50 while the *k*-value for the character fit of 50% is: (0.5×7.50)/1-0.5) = 7.50. Resulting *k*-values are in Appendix S5. We chose to conduct the IW analysis because studies have proven its effectiveness in recovering topologies congruent with those of total evidence phylogenies (*e.g*., Reemer and Ståhls, 2012), sometimes outperforming other methods (*e.g*., Goloboff *et al*., 2017). This weighting approach is also frequently used in the analysis of morphological datasets along with EW (*e.g*., Kim and Ahn 2016; Marín *et al.*, 2017; Rocha Filho and Packer, 2017).

We searched for trees under both weighting schemes (EW and IW) by implementing sectorial searches with tree drifting (TD) and tree fusing (TF), and ratchet runs with TD and TF. We used the following search: keep a maximum of 10,000 random trees, 500 random addition sequences, and 1000 ratchet iterations, including 100 cycles of TD and 100 rounds of TF per iteration. In EW analysis, we estimated branch robustness using standard bootstrap (sample with replacement) and absolute Bremer support in TNT, and plotted the values on the strict consensus topology obtained from the final TNT parsimony run. We used 10,000 bootstrap replicates under a heuristic tree search that consisted of 10,000 replicates of Wagner trees with random addition sequences, followed by Tree Bisection Reconnection (TBR) branch swapping (saving 10 trees per replicate). Resulting values per node represent frequency differences GC for Group present/Contradicted (Goloboff *et al*., 2003b). We calculated Bremer support by withholding 10,000 suboptimal trees up to 10 steps longer than the most parsimonious trees under a traditional search (10,000 replicates of Wagner trees, followed by TBR, saving 10 trees per replicate).

We assessed the performance of each IW analysis by comparing the number of most parsimonious trees, tree length, retention index, and node support. For the latter, we used Jacknife with symmetric resampling expressed as GC frequency-difference values, which Reemer and Ståhls (2012) found useful when determining the reliability of trees under different *k*-values. We searched trees under each *k*-value and used 1000 replicates under a heuristic tree search that consisted of 10 replicates of Wagner trees with random addition sequences, followed by Tree Bisection Reconnection (TBR) branch swapping (saving 10 trees per replicate). We calculated average and median GC frequency-difference from the value displayed at each node, which we plotted on the resulting tree or strict consensus tree (if the analysis yieled more than one most parsimonious tree). Groups that are more often contradicted than supported displayed values in brackets, which we considered as having a support of zero and excluded them from the calculations. In addition, we calculated in TNT the SPR distance (Goloboff, 2008) between the resulting topology from each IW analysis and the topology obtained from the analysis of the combined full dataset.

We visualized cladograms in WinClada, collapsing unsupported nodes and using DELTRAN (slow) for character optimization; when the choice is equally parsimonious, the latter favours repeated origns of characters over reversals. We used the abbreviations MPT, L, CI, and RI for most parsimonious tree, tree length, and consistency and retention indices, respectively. In the text, we referred to characters states in the form 21-1, where 21 is the character and 1 the character state.

#### Maximum likelihood

We conducted analyses using the message passing interface (MPI) version of IQ-Tree 1.5.5 (Nguyen *et al.*, 2015). We used the command ModelFinder (Kalyaanamoorthy *et al.*, 2017) to select the substitution model during the analyses to avoid *a priori* models. To examine the effects of introns on the phylogeny, we first ran ML analyses on the concatenated molecular datasets with and without introns, as described above. While introns had negligible effects on the topology of both the tribal- and generic-level phylogenetic trees, their inclusion resulted in higher support values in the generic-level analysis (Appendix S6). Thus, we used the molecular data without introns in the combined analysis of the tribal-level phylogeny and the dataset with introns in the generic-level study. Then, we conducted ML analyses on these combined datasets, giving the morphological data a separate partition. We estimated branch support using 1,000 replicates of ultrafast bootstrapping (Minh *et al.*, 2013).

#### Bayesian inference

We conducted these analyses using the MPI version of MrBayes 3.2.6 (Ronquist *et al.*, 2012b; Zhang *et al.*, 2016) on the total-evidence datasets described above, with the morphological data given its own partition. We did not select an *a priori* substitution model; instead, we used the reversible-jump Markov chain Monte Carlo (MCMC) method with gamma-distributed rate variation across sites to test the probability of different models *a posteriori* during the analysis (Huelsenbeck *et al.*, 2004; Ronquist *et al.*, 2012b). We conducted two different types of BI analyses, a time-free and a time-calibrated analysis. For the time-free analysis, we did not add further specifications following the input for the reversible-jump MCMC. We set the MCMC generation to run 10 million generations using four chains (three heated, one cold) with the swap number set to one, and a temperature of 0.1 for the heated chains. We monitored MCMC convergence of both time-free and time-calibrated analyses with Trace v.1.6 (Rambaut *et al.*, 2014). We considered a value of ESS ≥ 200 a good indicator of convergence.

We used a tip-dating approach (Pyron, 2011; Ronquist *et al.*, 2012a) for the time-calibrated analyses. First, we estimated the base molecular clock rates as outlined by Ronquist *et al.* (2012a). For the tribal-level analysis, the base clock rate and root age were informed using the age of the oldest crown bee fossil (Engel, 2000, 2001) for the minimum age, and 120 Ma for the mean age based on previous estimates for the age of Megachilidae by Cardinal and Danforth (2013). Then, we used the divergence time estimates generated by the tip-dated tribal-level analysis to inform these priors in the generic-level analysis. We used the fossilized birth-death macroevolutionary model (Heath *et al.*, 2014) following the methods of Zhang *et al.* (2016). The sampling strategy was set to diversity, with sampling probability set to 0.016 in the tribal-level analysis (66 megachilids sampled from the 4,105 known species) and to 0.029 for the generic-level analysis (59 megachilines of the 2,000 known species of the tribe Megachilini). We assigned an uncorrelated relaxed clock model IGR with the prior on rate variation across lineages set to exponential 10. We gave the fossils a uniform calibration prior based on the dated ages of their amber deposits (Engel, 2001).

To aid convergence in the time-free analyses, we applied several constraints on well-supported groups. In the tribal-level analysis, we constrained both melittine taxa, as well as Apidae and Megachilidae, as sister groups. In the generic-level analysis, we applied constraints to unite taxa representing the following groups: Lithurgini, Osmiini, Megachilini, and Osmiini + Megachilini. We also conducted additional analyses constraining the fossil species *M.* (*Chalicodomopsis*) *glaesaria* with different taxa of *Megachile s.l*. based on the results of the EW parsimony analysis of the morphological dataset (Appendix S6).

The MCMC generation settings for the time-calibrated analyses were initially set identically to the time-free analyses, and we completed the preferred tribal-level tip-dated tree under these settings. However, we experienced considerable difficulty getting both the tribal-and generic-level tip-dated analyses to converge. Convergence of the generic-level analysis was accomplished after providing the time-free total-evidence Bayesian tree as a starting tree, lowering the heated chain temperature to 0.010, and increasing the MCMC generations to 50 million. Thus, we applied a similar approach to the tribal-level analysis, increasing the number of chains and swaps, and increasing MCMC generation to 200 million, but it still did not attain convergence. Furthermore, allowing the tip-dated tribal-level analysis to run longer resulted in unrealistically old divergence time estimates, placing the root of the tree in the Permian with a median age of 256 Ma (Appendix S6). Thus, the resulting preferred tip-dated tribal-level tree did not reach convergence, but the topology is identical to that obtained across the different attempts to reach convergence (Appendix S6). It has similar support values, but it differs in the age estimates, which are considerably more realistic and in line with previous estimates using node-calibration approaches (Cardinal and Danforth, 2013; Litman *et al.*, 2013).

### Phylogenetic signal of morphological characters

The current trend in evolutionary biology is the analysis of large datasets composed of both molecular and morphological data. Thus, to facilitate future comparative cladistics analyses, we assessed the phylogenetic signal of the scored characters and determined the level of homoplasy among character sets. Given the current taxonomic problems within Megachilini, we conducted these analyses on the morphological dataset used in the generic-level study only. We compared the median value of RI per character set and conducted partitioned phylogenetic analyses. We grouped characters by sex (male and female characters), tagmata (pro-, meso-, and metasoma), and by the following set of characters of taxonomic importance in the diagnosis and recognition of supraspecific groups: female mandible, female terminalia (T6, S6, and sting apparatus), male legs, and male terminalia (T6, T7, S5–S8, and genitalia). We conducted phylogenetic analyses only to the subset of male and female characters using the settings for EW analyses under parsimony as indicated above. For each analysis, we recorded the number of MPT and tree statistics (L, CI, and RI).

### Origin and patterns of variation of the interdental lamina

To determine the possible mandibular structure(s) from which interdental laminae originated, we conducted a comparative study of the female mandible across all Megachilini taxa used in the phylogenetic analyses. We made inferences based on topological correspondence, a robust criteria for recognizing primary homologies (*e.g*., Rieppel and Kearney, 2002; Agnarsson and Coddington, 2008). We used Mesquite (Maddison and Maddison, 2018) to trace the evolutionary history of the interdental lamina. We reconstructed ancentral character states using a Parsymony model with unordered character states and visualized them on the tree topology obtained from the analysis of the combined full dataset using BI.

To test for character association between the LC behavior and some female cephalic and mandibular features, we used the phylogenetic pairwise comparison implemented in Mesquite. We used the option that searches for pairs of taxa contrasting in the state of two characters, with a maximum number of pairings set to 1000000 pairs. To maximize the number of pair comparisons, we used the resulting tree topology from the analysis analysis of the combined full dataset using BI. We tested the following five characters that we considered dependent characters: ocelloccipital distance (character #12), mandible length (#28), mandibular apical width (#33), shape of acetabular interspace in the outer surface of mandible (#38), and pubescence on the adductor interspace in the inner surface of mandible (#56). We chose these characters because they appeared to be under the same selective pressure, which is the type of nesting material used (Mitchell, 1980; Williams and Goodell, 2000). The size and shape of the mandible also influences the size and shape of the head, as the latter contains the mandibular musculature. Thus, we used these characters as proxy of the head size and mandibular size and shape. We also included the pubescence on the adductor interspace of the mandible because setae on this area appear to be lost in LC bees. We tested each of these five characters for association with the presence of interdental laminae, a feature unquestionably indicative of LC behavior. For analysis, we scored each species as having either character state 0 when these laminae are absent, or as having character state 1, when they are present on at least one dental interspace. We did not compare these characters with the presence or absence of LC behavior because the nesting biology of most species in our analysis is unknown. Additionally, some species that lack interdental laminae still cut leaves while others do not (*e.g*., *M. montivaga* Cresson). Thus, assuming an absence of LC behavior in species that lack interdental laminae is not applicable.

## Results

### Total-evidence phylogeny of Megachilidae

In the preferred Bayesian analysis (Fig. 8), Pararhophitinae resulted as the sister group of Lithurginae, both taxa sister to Megachilinae. Within the latter subfamily, Dioxyini were the sister group of Glyptapini while Aspidosmiini rendered Ctenoplectrellini paraphyletic. Megachilini also rendered Osmiini paraphyletic, as they clustered with the osmiine genera *Afroheriades* Peters and *Pseudoheriades* Peters. The remaining osmiines are together in another clade (Fig. 8). The origin of crown Megachilidae was estimated at a median age of 111.3 Ma (95% highest posterior density 80.94–127.56 Ma) with crown Megachilini at 42.0 Ma (24.75– 49.55 Ma). A similar topology resulted from the ML analysis of the combined dataset with extant taxa only (Appendix S6C). However, Pararhophitinae were the sister group of Lithurginae + Megachilinae, Aspidosmiini and Dioxyini clustered in the same clade, and the osmiine genus *Ochreriades* Mavromoustakis resulted as the sister group of a clade that contained Megachilini and the remaining Osmiini.

**Fig 8.**
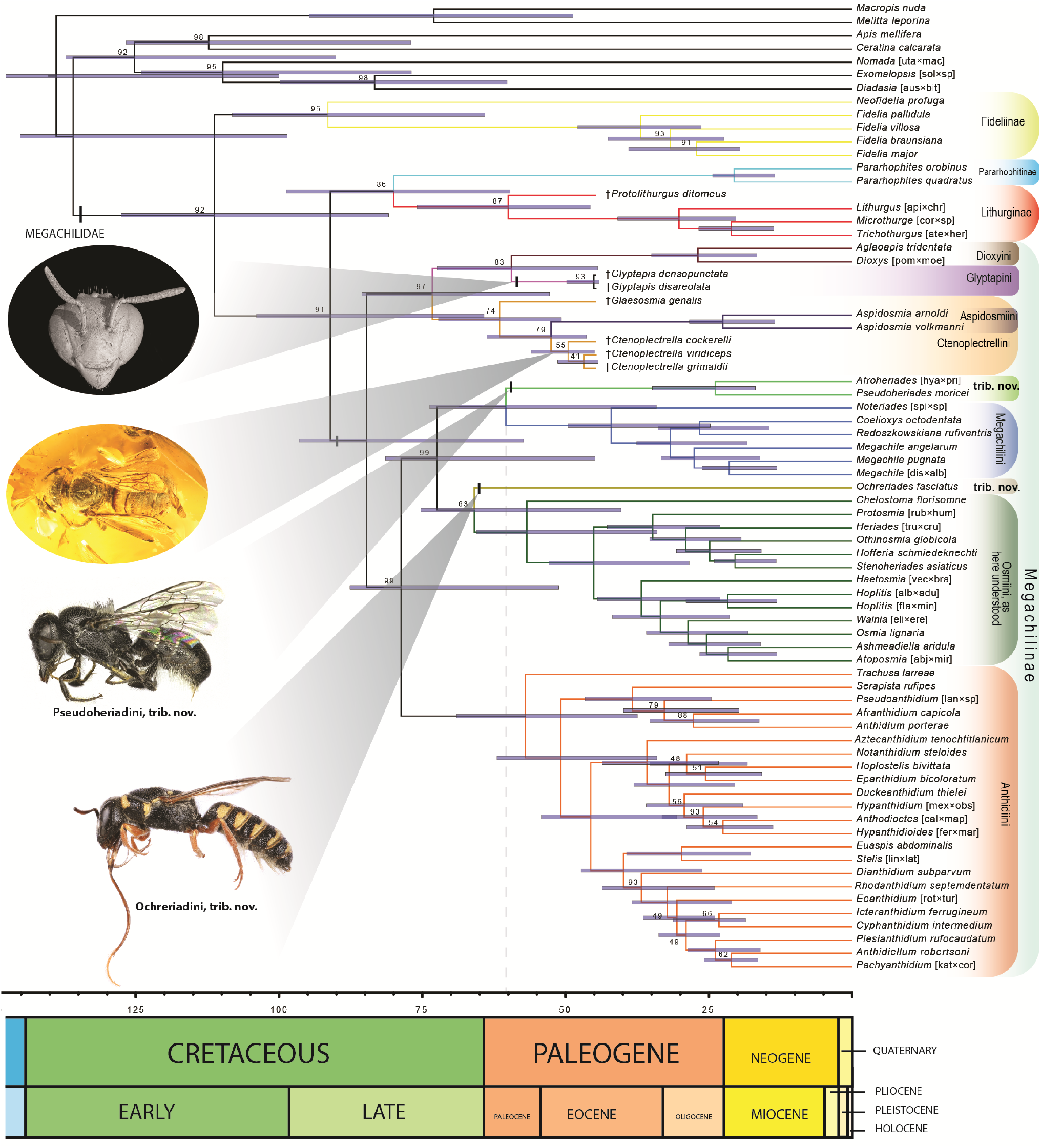
Preferred total evidence dated phylogeny of Megachilidae. Majority-rule consensus tree from Bayesian analysis using fossils as terminals under the FBD tree prior. Blue bar at each node represents the 95% highest posterior density age rage. Posterior probability below 100 indicated above each node.

### Morphological-based phylogeny of Megachilini

The analysis of the morphological data matrix under EW yielded 30 MPTs (L = 2364, CI = 13, RI = 57); nine nodes collapsed in the consensus tree (Fig. 9) and most branches were weakly supported by homoplastic characters. The clade of cleptoparasitic bees (Clade A) consisting of *Coelioxys* and *Radoszkowskiana* resulted as the sister group of all other Megachilini. *Noteriades* and *M.* (*Rhodomegachile*) (Group 2) are successive sister taxa, with the latter being sister to the remaining *Megachile s.l*. Most subgenera of Group 2 clustered in multiple clades along the tree, except for *M*. (*Mitchellapis*) and *M*. (*Megella*), which were in a large derived clade (Clade C) containing all subgenera of Group 1 and *M*. (*Creightonella*) (Group 3). The fossil taxon, *M*. (*Chalicodomopsis*), resulted in a clade that included *M*. (*Matangapis*), *M*. (*Chelostomoda*), *M*. (*Hackeriapis*), and other hoplitiform or heriadiform taxa of Group 2 (Clade B).

**Fig 9.**
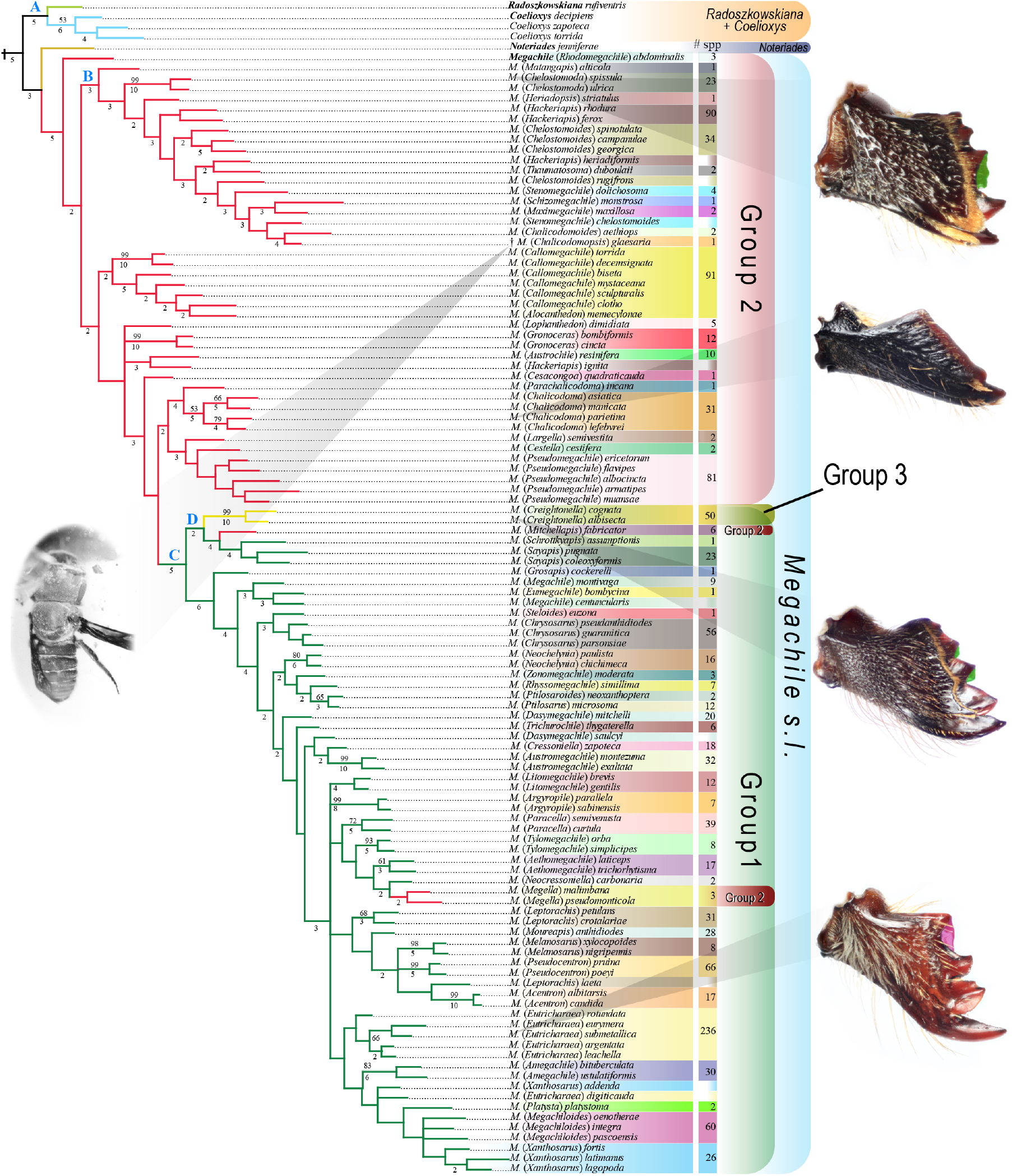
Strict consensus tree of 30 most parsimonious trees obtained under equal weighting. Numbers above nodes are standard bootstrap values, numbers below nodes are absolute Bremer values. Branches without numbers indicate bootstrap values below 50% and Bremer values of 1. A capital letter above a node indicates a clade discussed in the text. Species within boxes of the same color correspond to the same subgenus of *Megachile s.l*. following Michener’s (2007) classification. The colored column after the species names indicates approximate number of species per subgenus. Half-colored boxes without a number correspond to species that did not cluster with the other species of the same subgenus included in the analysis. Species richness taken from Michener (2007), Moure *et al*. (2007), and Ascher and Pickering (2018). Mandibles with interdental laminae highlighted in green (odontogenic) and pink (ctenogenic).

Implied weighting analyses under the 11 *k-*values each resulted in a single MPT, except for two analyses (character fits 66 and 70%) that yielded two MPTs. All resulting trees are longer than the MPTs obtained under EW, but have similar CI and RI values. The topologies obtained with character fits 54 and 62 to 70% have the highest SPR values (Appendix S5A). However, the median GC frequency-difference value was similar among analyses (*H* = 0.773, *p* > 0.99) and it was not associated with SPR values (Spearman’s correlation, *r*_*s*_ = 0.146, *p* = 0.667). Likewise, the number of supported nodes was similar among analyses [Chi-Square Goodness-of-fit test, *X*^*2*^ (10, *n* = 715) = 1.139, *p* = 1.00].

Implied weighting analyses largely recovered the backbone topology of Megachilini, but the position of several taxa significantly changed among *k-*values (Appendix S5B–E). For example, *Noteriades* resulted as the sister group of Megachilini in all analyses, except in those with character fits 86 and 90%, in which it appeared as sister of *Megachile s.l*. as in EW analysis. *Megachile* (*Chelostomoda*), a member of Group 2 of subgenera (see Clade B in EW consensus, Fig. 9), clustered in Clade D with *M*. (*Mitchellapis*) and *M*. (*Sayapis*) in analyses with character fits ranging from 50 to 70% (Appendix S5). In remaining analyses, *M.* (*Chelostomoda*) clustered within Clade B. *Megachile* (*Rhodomegachile*) and *M*. (*Chalicodomopsis*) were sister groups, either as part of Clade B (analyses with character fits 50 and 58%), or as the sister group of all other Megachilini (analyses with character fits 62 to 86%) excluding *Noteriades* and the Clade A. In the analysis with the highest character fit, both taxa resulted in positions similar to those in the EW consensus topology. Likewise, *M*. (*Gronoceras*) (Group 2) resulted either as the sister group of all other Megachilini, excluding *Noteriades* and Clade A (character fits 50 and 54%), or in the same clade with *M*. (*Lophanthedon*) (Character fits 58, 70–90%). *Megachile* (*Callomegachile*) *decemsignata* and *M.* (*Callomegachile*) *torrida*, members of Group 2, clustered with *M*. (*Lophanthedon*) in Clade D (Character fits 50–58, 66, 70%) or with other members of *M*. (*Callomegachile*) in the remaining analyses. *Megachile* (*Matangapis*) was the sister group of *M*. (*Heriadopsis*) and clustered within Clade B (Character fits 50–58%). However, in analyses with character fits 62–70, 86% (Appendix S5C–E), *M*. (*Heriadopsis*) remained within the same clade but *M.* (*Matangapis*) resulted singly in a branch after the clade consisting of *M.* (*Rhodomegachile*) and *M*. (*Chalicodomopsis*). In remaining analyses, *M.* (*Matangapis*) was the sister group of all members of Clade B, as in the EW consensus. Clade C was consistenly recovered, but the number of clades changed among analyses.

### Total-evidence phylogeny of Megachilini

In the analysis of the reduced dataset, the fossil *M. glaesaria* resulted as the sister group of all other Megachilini, Clade A was the sister of some members of Group 2 of subgenera, Clade B was segregated in several clades, and *M*. (*Chelostomoda*) was sister to Clade C (Fig. 10). The origin of Clade C was estimated at a median age of 15.61 Ma (95% highest posterior density 12.80–19.16) and that of *M*. (*Chelostomoda*) + Clade C at a median age of 14.92 Ma (11.83– 19.42). Constraining the position of *M. glaesaria* (Appendix S6J, K) to the clade that includes *M*. (*Thaumatosoma*), as in the EW topology, yielded older estimates for the origin of Clade C (23.63 Ma, 18.96–28.93) and for that of *M*. (*Chelostomoda*) + Clade C (24.82 Ma, 20.25–30.24). Likewise, constraining the position of *M. glaesaria* to the clade that includes *M*. (*Matangapis*) yielded older estimates for the origin of Clade C (22.42 Ma, 19.11–26.98) and for that of *M*. (*Chelostomoda*) + Clade C (23.58 Ma, 20.27–28.24). We obtained the same topology when we analyzed the full dataset (Fig. 11). However, estimated median ages were older for Clade C (17.6 Ma, 15.22–21.80 Ma) as well as for that of *M*. (*Chelostomoda*) + Clade C (18.6 Ma, 16.12–23.35 Ma). A similar topology resulted from the ML analysis of the reduced dataset (Appendix S6H), except that *Noteriades* was the sister group of all Megachilini, and *M*. (*Matangapis*) and *M*. (*Heriadopsis*) clustered with members of Clade C (Fig. 1S).

**Fig 10.**
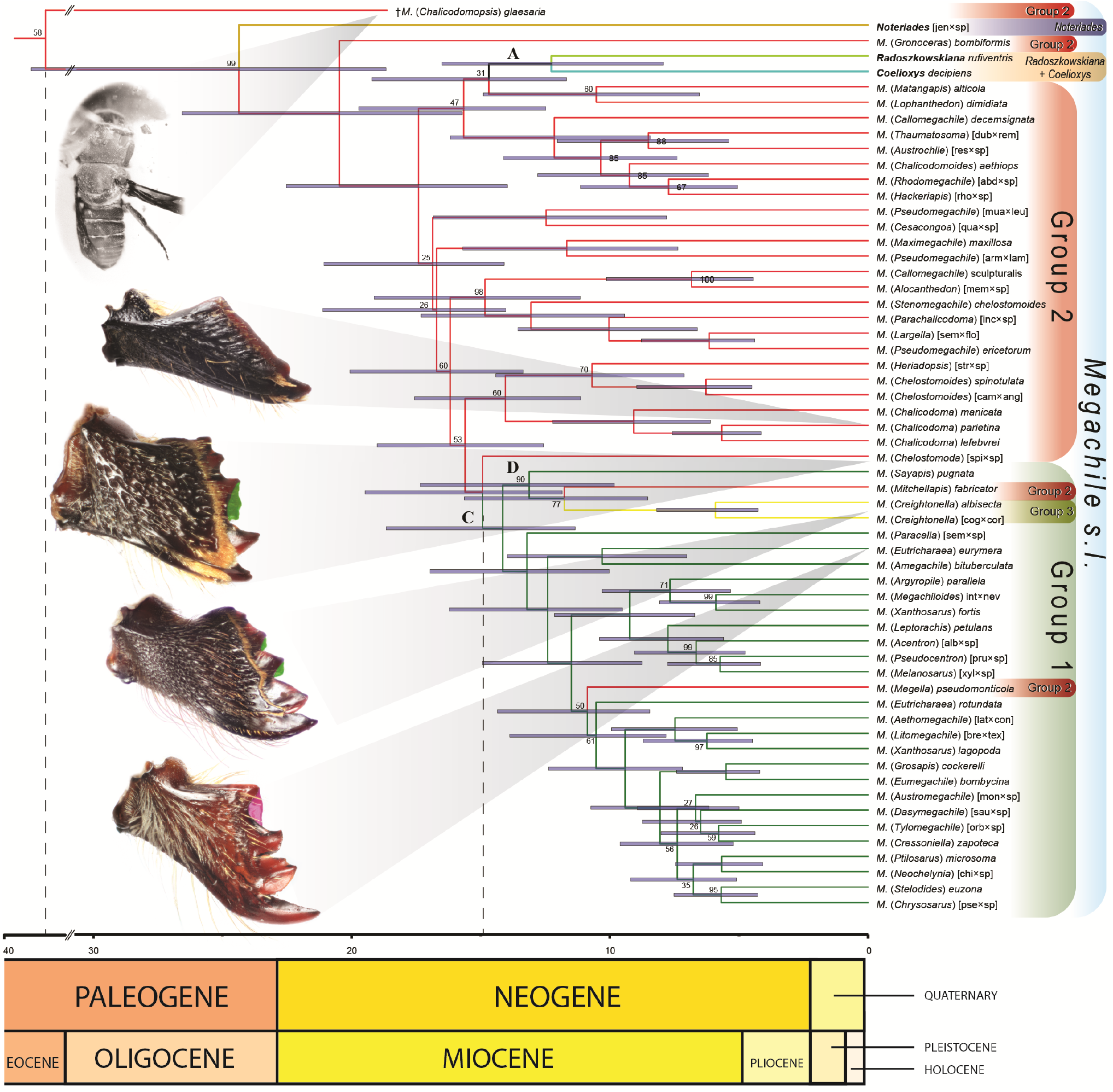
Preferred total evidence dated phylogeny of Megachilini from the analysis of the reduced morphological data matrix (67 OTUs). Majority-rule consensus tree from Bayesian analysis using fossils as terminals under the FBD tree prior. Blue bar at each node represents the 95% highest posterior density age rage. Posterior probability below 100 indicated above each node. A capital letter above a node indicates a clade discussed in the text. Mandibles with interdental laminae highlighted in green (odontogenic) and pink (ctenogenic).

**Fig 11.**
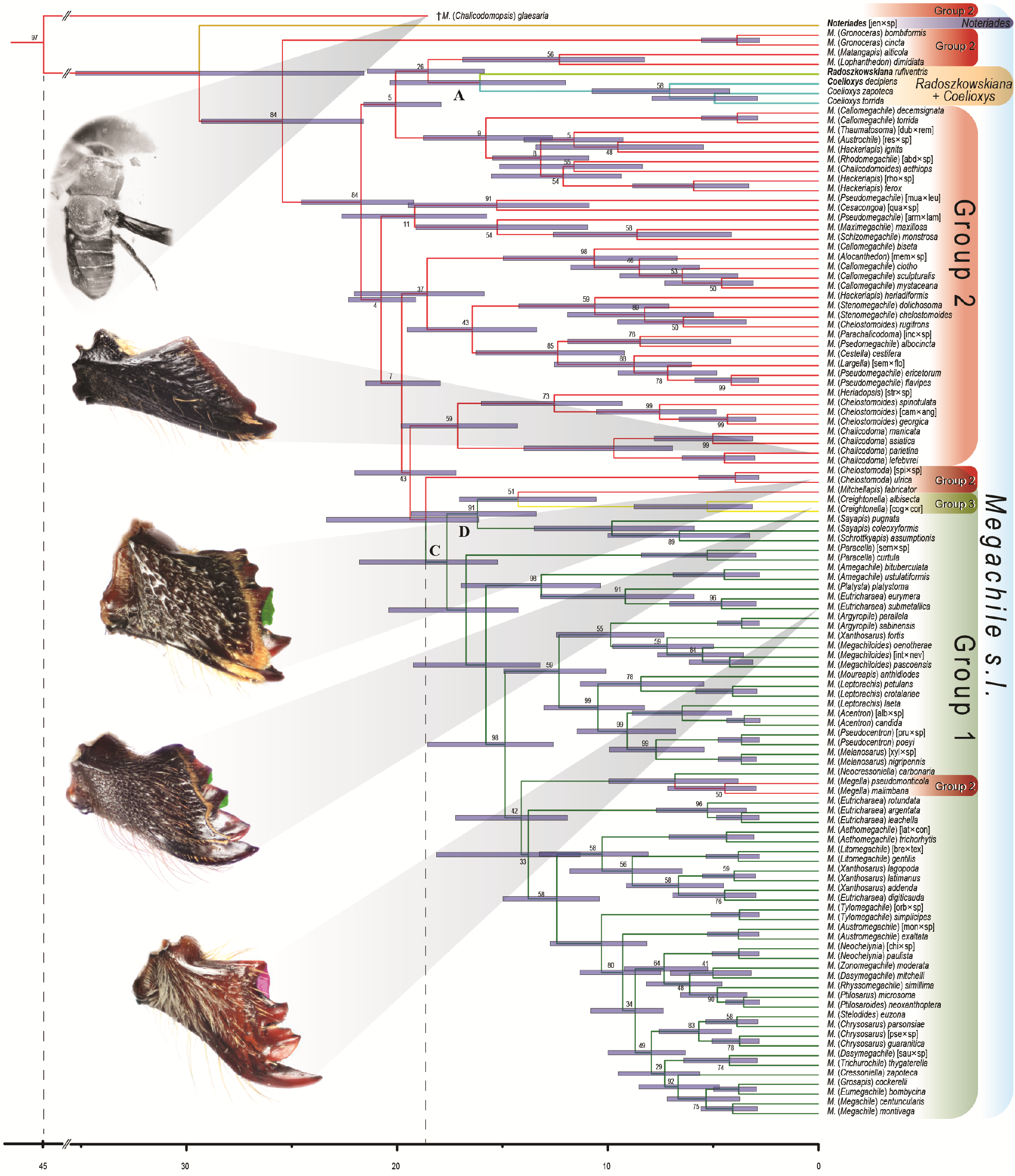
Total evidence dated phylogeny of Megachilini from the analysis of the full morphological data matrix (122 taxa). Majority-rule consensus tree from Bayesian analysis using fossils as terminals under the FBD tree prior. Blue bar at each node represents the 95% highest posterior density age rage. Posterior probability below 100 indicated above each node. A capital letter above a node indicates a clade discussed in the text. Mandibles with interdental laminae highlighted in green (odontogenic) and pink (ctenogenic).

### Phylogenetic signal of morphological characters

The female and male character sets were similar in the median RI value (Mann–Whitney test, *U* = 13334, *p* = 0.192), as well as in the percentage of unambiguous synapomorphic characters (Table 1). However, unlike the analysis of the male characters that resulted in a highly unresolved tree, female characters recovered Megachilini and several major lineages (not shown). For both sexes, we did not find statistical significant differences between the median RI values and percentage of unambiguous synapomorphic characters between character sets (Female, RI value: *U* = 1452, *p* = 0.183; % unambiguous synapomorphic characters: Chi-square goodness of fit test, *X*^*2*^[1] = 0.67, *P* = 0.414. Male: *U* = 1630, *p* = 0.209; *X*^*2*^[1] = 0.014, *P* = 0.907).

**Table 1.**
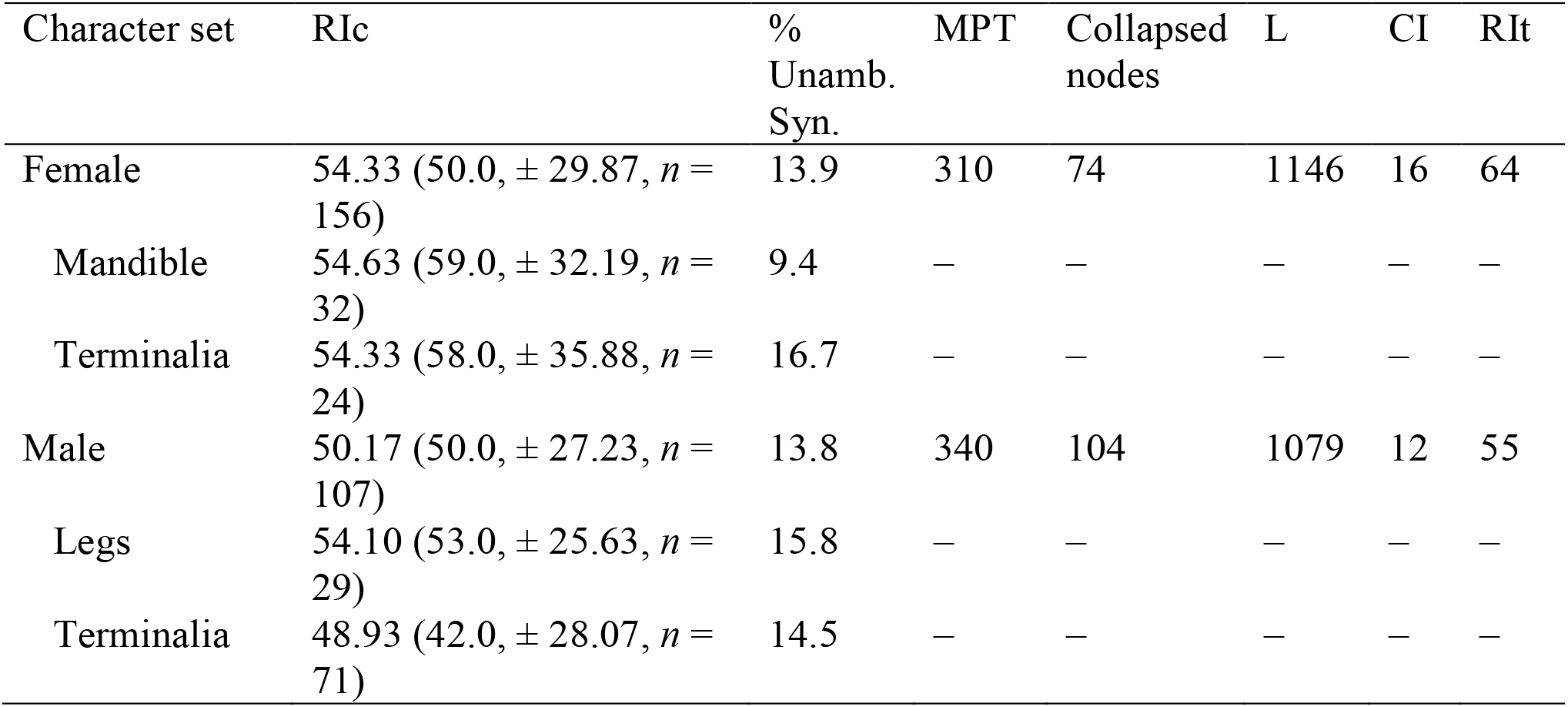
Retention index per morphological character set and quantitative descriptors of trees obtained from partitioned analysis using female and male characters. RIc = average retention index of character set followed, in parentheses, by median, standard deviation, and number of characters; % Unamb. Syn. = Percentage of unambiguous synapomorphic characters in the analysis of the full data matrix; MPT = number of most parsimonious trees; Collapsed nodes = number of nodes that collapsed in the consensus strict tree; L = tree length; CI = consistency index; RIt = retention index of MPTs. Non-applicable are indicated by an en dash symbol (–).

### Origin and evolution of the interdental lamina

We found two types of interdental laminae that seem to develop from different structures in the mandible. The first type is clearly a ventral extension of the *corono-radicular ridge* (CR), a strong ridge that runs basally from the apex or cusp of the inner surface of each tooth (Fig. 3B). This CR ridge is usually strongest on Mt1, sometimes continuing dorsally into the adductor interspace or curving ventrally, thus running pararell to or towards the adductor ridge (Figs. 3C, D). Gonzalez *et al*. (2012) recognized the portion of this ridge running pararell to or towards the adductor ridge as the adductor apical ridge. Interdental laminae originating from teeth, hence called *odontogenic laminae*, often partially fill the dental interspaces (*i.e*., incomplete). Even in species of *M.* (*Schrottkyapis*) and *M*. (*Stelodides*), which have secondarily lost interdental laminae, there still is a hidden small projection from the CR ridge of Mt3 suggestive of a lamina.

The second type of interdental lamina is usually complete and is likely an extension from a transverse ridge at the base of the teeth, which runs parallel to the inner fimbria on the inner surface of the mandible. When this transverse ridge is laminate and apically extended so that it can be seen from the outside of the mandible in between the teeth, it becomes an interdental lamina. In megachiline bees that never developed interdental laminae, this ridge runs from the upper carina to the base of Mt2, merging with the CR of that tooth, and forming a rather concave surface, which sometimes is divided by the CR of the other teeth (Fig. 3B). In *M*. (*Chelostomoda*), *M*. (*Creightonella*), and *M*. (*Sayapis*), such a transverse ridge is laminate or nearly so, but it does not extends enough apically to form an interdental lamina (Fig. 3C–E). In species that have secondarily lost interdental laminae, this transverse ridge is conspicuous and distinctly elevated compared to that of most species of Group 2 that never developed interdental laminae. Michener and Fraser (1978) recognized that the transverse ridge was associated with the inner fimbria and thus named it *fimbrial ridge* or *carina*. Consequently, we referred to the lamina that develops from this fimbrial ridge as either the *fimbrial lamina* or *ctenogenic lamina* (from the Greek κτε, *kteis*, meaning “comb”).

Odontogenic laminae evolved first and have secondarily been lost or modified multiple times. In constrast, ctenogenic laminae developed in more derived clades of LC bees and have been lost or modified comparatively less times than odontogenic laminae (Fig. 12). Odontogenic laminae are often restricted to Mt3 and thus visible in the second interdental space only (Figs. 2E–H), except in *M*. (*Creightonella*) where they are also present on Mt4–5 (Fig. 3G). Mandibles of species that only have odontogenic laminae tend to have a thick distal margin with the acetabular interspace of the outer surface gently curving towards the base of the mandible, thus resembling the mandible of species of Group 2 that never developed interdental laminae (Fig. 2C). In contrast, mandibles that have both odontogenic and ctenogenic laminae exhibit a distinct basal, lateral surface and a distal, anterior surface because the acetabular interspace is clearly flattened or depressed (Figs. 2E–J). The mandible is also much broader apically, with the distal margin flattened, and with both types of laminae often at different levels from the mandibular margin and from each other. However, in some species whose mandible is flattened at apex, both laminae are thin and nearly at the same level with the interspace margin, sometimes fused and indistinguishable. The CR ridges are usually absent in apically flattened mandibles, except for that of Mt1. In some species with thicker mandibular apex, the ctenogenic interdental laminae are narrow (not reaching apex of teeth), well behind or deeper to the interspace margin, and often hidden by it (Fig. 2D). A mandible of this kind would appear to lack laminae when seen in it from the outside.

**Fig 12.**
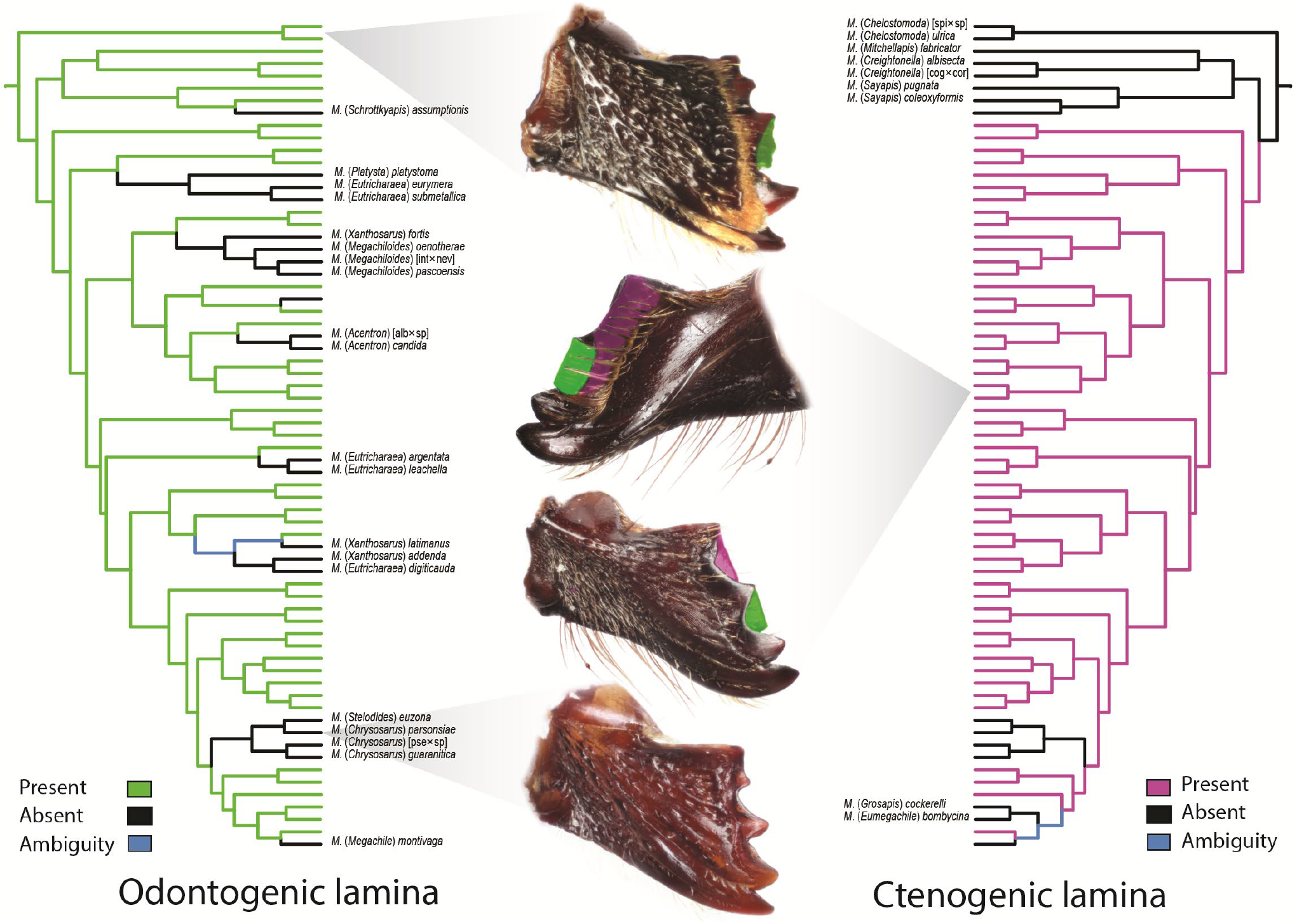
Parsymony reconstruction of the two types of interdental laminae of the leaf-cutter bee mandible. We used the tree topology obtained from the total-evidence analysis of the full data set (122 taxa) to visualize character states on the clade of leaf-cutter bees. All photographs are outer views of the mandibles, except for the second from top to bottom, which is an inner view of the mandible below. Odontogenic lamina highlighted in green and ctenogenic lamina in pink.

According to the phylogenetic pairwise comparisons, the presence of interdental laminae is not associated with characters #12, 28, 33, 38, or 56 (ocelloccipital distance, mandible length, mandibular apical width, shape of acetabular interspace, and pubescence on the adductor interspace, respectively), which we used as proxy of head size and mandible size and shape (Table 2).

**Table 2.** Phylogenetic pairwise comparisons between the presence of interdental laminae in the female mandible (independent character) and some female cephalic and mandibular characters (dependent characters). See materials and methods for description of each character. # Pairs = number of pairs contrasting in the state of two characters; Relationship = number of pairs with a positive (+) or negative (-) relationship. In a positive relationship, one of the paired species has a character state 1 for both characters and the other species character state 0 for both characters. In a negative relationship, one of the paired species has a character state 1 for one character and a 0 for the other character while the other species has the opposite. *P*-value: significance value for the number of pairs contrasting in the state of two characters; Range of *P*-values: range of significance values for all optimal set of pairs of pairwise comparisons.

| Compared character # | # Pairs | Relationship | *P*-value | Range of *P*-values |
| --- | --- | --- | --- | --- |
| 12-ocelloccipital distance | 4 | 1<sup>+</sup>, 3<sup>-</sup> | 0.313 | 0.313–0.688 |
| 28-mandible length | 3 | 2<sup>+</sup>, 1<sup>-</sup> | 0.5 | 0.5–0.5 |
| 33-mandibular apical width | 3 | 2<sup>+</sup>, 1<sup>-</sup> | 0.5 | 0.5–0.5 |
| 38-acetabular interspace | 4 | 3<sup>+</sup>, 1<sup>-</sup> | 0.313 | 0.313–0.313 |
| 56-pubescence on adductor interspace | 4 | 3<sup>+</sup>, 1<sup>-</sup> | 0.313 | 0.063–0.313 |

## Discussion

### Origins of LC behavior and interdental laminae

We used for the first time a total-evidence tip-dated approach to estimate the origin of LC behavior in bees. We aimed to assess the temporal discrepancy between the recent age (20–25 Ma) estimated from molecular analyses using a node-dating approach and the much older age (60 Ma) suggested by fossil traces (Michez *et al*., 2012; Litman *et al*., 2011; Trunz *et al*., 2016). A tip-dating approach allows the incorporation of all available fossils into phylogenetic analyses, which not only expands the taxonomic coverage and ancentral character states, but also provides more accurate fossil information to the analysis than *ad hoc* node age constraints (*e.g*., Pyron, 2011; Ronquist *et al*., 2012a; Slater *et al*., 2013; Arcila *et al*., 2015). However, tip-dating analyses tend to estimate much older divergence times when compared to the node-dating approach, sometimes unrealistically (*e.g*., O’Reilly *et al*., 2015; O’Reilly and Donoghue, 2016). In our case, the tip-dated analyses suggested a similar recent divergence time (15–25 Ma) to those obtained from molecular analyses using a node-dating approach (Figs. 10, 11). The position of the fossil *M. glaesaria* as the sister group to all other Megachilini, which differed from that obtained in the morphological analysis (Fig. 9), influenced the divergence time estimates. Constraining the position of this fossil near *M*. (*Thaumatosoma*), as in the morphological analysis, yielded older estimates for the origin of LC bees, yet these values were never greater than 30 Ma. Although our results support the idea that Eocene trace fossils might not be the result of LC bees, we used a low number of fossils in our analyses, particularly in the generic-level phylogeny. Including more fossil taxa across different clades might increase the precision of tip-dating estimates (Pyron, 2011; Ronquist *et al*., 2012a; Dos Reis and Yang, 2013).

Although the presence of interdental laminae is associated with leaf cutting, these structures seem not to be required to express this behavior. For example, LC bees that secondarily lost these laminae retained this behavior (*e.g*., Laroca *et al*., 1992; Zillikens and Steiner 2004; Torretta *et al*., 2014). Leaf-cutter ants, which never developed interdental laminae (Figs. 13A– C), are also able to cut leaves, petals, and grasses efficiently. Shifts in behavior might act as drivers of evolutionary diversification and phenotypic change (*e.g*., Duckworth, 2009; Lapiedra *et al*., 2013). Thus, it is likely that interdental laminae might have evolved after the LC behavior was already in place. This idea is supported by the use of chewed leaf pulp, large petal pieces, and irregular leaf pieces in some Osmiini and in some *M*. (*Callomegachile*) that lack interdental laminae (Michener 2007; Rozen *et al*., 2010). Lower costs in handling and processing large leaf pieces when compared with masticated plant material, and a greater access to more readibly vegetative plant resources (flowers are not often continuously available), might have facilitated a transition to leaf cutting.

**Fig 13.**
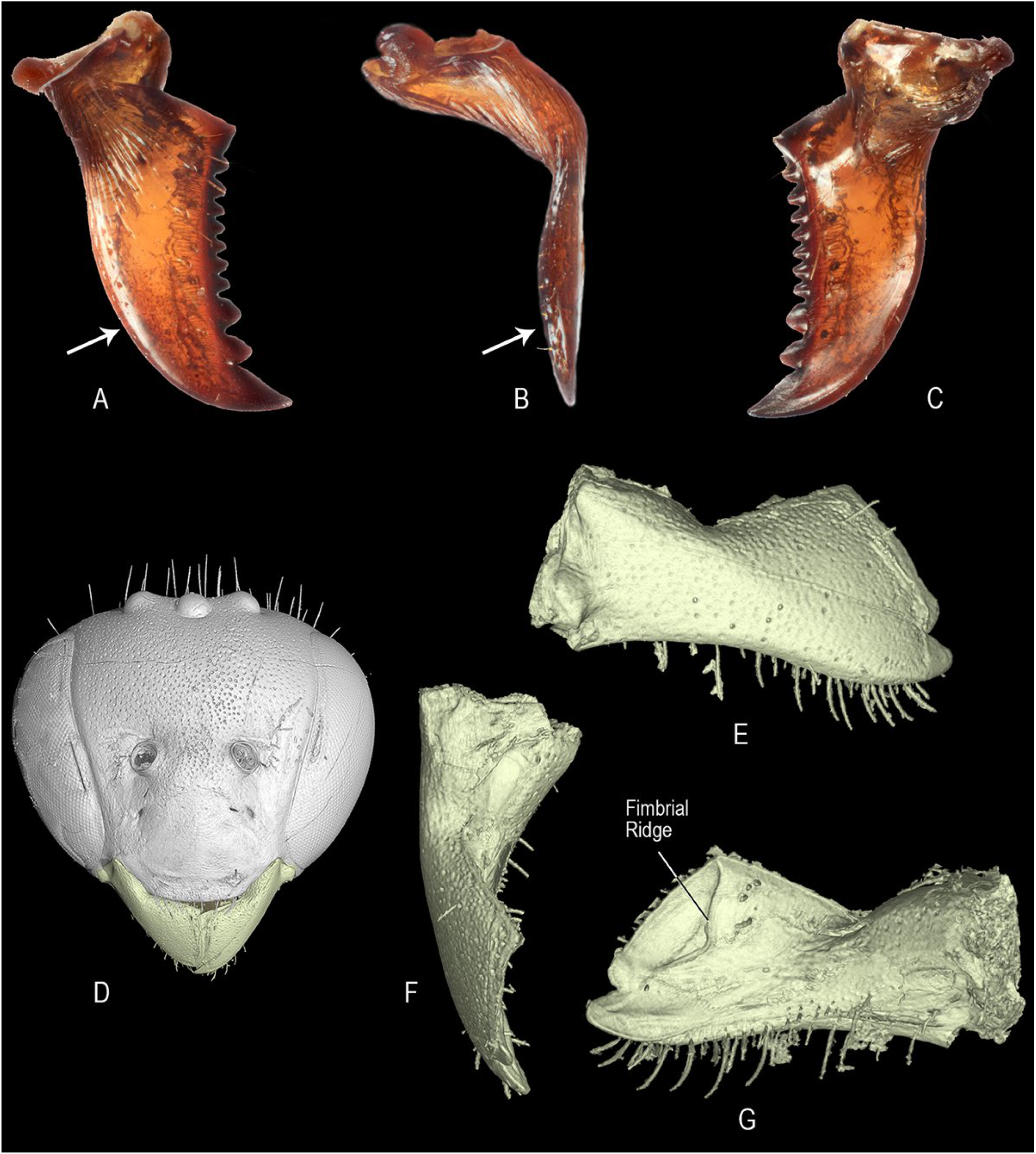
Female mandible of leaf cutter ants and extinct Baltic amber megachilids. **A–C**. Right mandible of leaf cutter ant (Formicidae: Attini, *Atta* sp.) in frontal, lateral, and inner views, respectively. Arrow points to the lower margin. **D–G.** Facial view of the head and right mandible of *Glyptapis* sp. (Glyptapini) in outer, superior, and inner views, respectively.

The development of interdental laminae might have either allowed a more efficient way to cut and process leaves, or allowed access to more leaf types and plants. Bee species with odontogenic laminae only or without it, cut irregularly margined leaf pieces. In species with both types of laminae, the margins of the leaf pieces are smooth (Michener, 2007). The study of MacIvor (2016) supports the second adaptive hypothesis. For example, *Megachile* (*Sayapis*) *pugnata* Say, a species with odontogenic laminae only, significantly uses less plant species than *M*. (*Megachile*) *centuncularis* and *M*. *rotundata*, species from lineages that developed both types of laminae (MacIvor, 2016). However, leaf choice might not only depend on the ability to cut certain types of leaves, but also on the local availability as well as on their chemichal, mechanical, and antimicrobial properties.

If the derived clade of LC bees evolved recently, and interdental laminae are not required for and likely evolved after the LC behavior, which insects are then responsible for the Eocene fossil leaf excisions? There are few insects capable of leaving similar cuts on leaf margins (for a discussion see Wedmann *et al*., 2009) and one of them are LC ants. However, LC ants are restricted to the Neotropics and they evolved even more recently (8–12 Ma) than LC bees (Schultz and Brady, 2008). Another possibility is that extinct lineages of megachilid bees from the Eocene (*e.g*., Ctenoplectrellini and Glyptapini) also cut leaves, or even extinct lineages within Osmiini or stem-group Megachilini.

An examination of the mandibular structure of fossils from Ctenoplectrellini and Glyptapini, both using light microscopy and CT scans, showed two different types of mandibles. In Glyptapini, the lateral surface of the mandible gently curves towards the apex; its distal margin has a single apical tooth and a long, edentate upper margin (trimmal expansion); and the inner surface of the mandible possess a distinct fimbrial ridge apically, which is long and parallel to the edentate upper margin (Figs. 13D, G). Thus, the mandible of Glyptapini resembles that of some species of Anthidiini that use resins, as well as that of some species of Group 2 of *Megachile s.l*. Unlike Glyptapini, the female mandible of Ctenoplectrellini is tridentate (males are bidentate) and has two distinct outer surfaces that merge rather abruptly with each other, one lateral basally and one anterior distally (Engel, 2001). Thus, the mandible of Ctenoplectrellini is somewhat similar to that of some species of *Osmia* Panzer. Finally, interdental laminae and fimbrial carina are completely absent in both tribes.

Based on the mandibular structure, it is therefore unlikely that these particular extinct taxa might have cut leaves, principally Glyptapini. However, it is likely that Ctenoplectrellini might have used plant resources as nesting materials because *Aspidosmia*, its extant closest relative, uses masticated leaf pulp to build their nests (Brauns, 1926). Thus, the identity of the Eocene LC insects remains elusive, and may or may not have included extinct Megachilinae or stem-group Osmiini or Megachilini.

We demonstrate for the first time that interdental laminae, the most distinctive and taxonomically significant feature of LC bees, developed from two different structures in the female mandible. We also show that odontogenic laminae evolved once and prior to the development of ctenogenic laminae, which developed from the fimbrial ridge and appeared in more derived LC taxa (Fig. 12). These findings have major implications for our understanding of the conection between character evolution and diversification. The most obvious is that interdental laminae represent two characters that evolved in a sequence of evolutionary events, not a single character that evolved once as current dogma surmises (Michener, 2007; Trunz *et al*., 2016).

Our analyses also suggest that the presence of odontogenic laminae is a putative synapomorphy for all LC bees, which exhibits more phenotypic plasticity than ctenogenic laminae. The multiple modifications and secondary losses observed across the phylogeny are more likely in this type of lamina perhaps because of its small size and less involved structural modifications relative to ctenogenic laminae. However, losses in ctenogenic laminae occurred independently in two clades, the *Chrysosarus* and *Megachile s.str*. groups of subgenera, sometimes also with the loss of odontogenic laminae (Fig. 12). While retaining the use of leaves, these groups have incorporated other nesting materials, such as mud or petals (*e.g*., Laroca *et al*., 1992; Banaszak and Romasenko, 1998; Zillikens and Steiner, 2004; Michener, 2007). At least one species, *M*. (*M*.) *montivaga* Cresson, makes nests entirely of petals (*e.g*., Mitchell 1935b; Michener, 2007; Orr *et al*., 2015). The environmental factors that favor the use of mud or petals in LC bees are unknown. As far as we know, species with secondarily lost interdental laminae co-occur in the same habitats with LC bees, and sometimes even occupy a wide range of habitats (*e.g*., *M. montivaga*).

Although the mandible of LC bees appears to be shorter, apically wider, and with rather two distinct outer surfaces than that of dauber bees, pairwise tests for trait correlation revealed no significant associations between each of them and the presence of interdental laminae. Pairwise comparisons also suggest similar independence of trait evolution between a short ocelloccipital distance and the absence of setae on the adductor interspace in the inner surface of the mandible (Table 2). Parsimony reconstructions suggest that these are ancestral character states (not shown). They are also present in some dauber bees in the genus *Megachile s.l*., as well as in some outgroups. Thus, these cephalic and mandibular features, rather than being associated with leaf cutting, are likely the result of a shift in processing and handling foreing materials (leaves, plant hairs, pebbles, resins, etc.) as an alternative to using glandular secretions to waterproof cells in the soil (Michener and Fraser, 1978; Eickwort *et al*., 1981).

### Phylogenetic relationships

The phylogenetic relationships obtained from our total-evidence analyses were generally congruent with previous phylogenetic hypotheses. However, they revealed new relationships, resolved phylogenetic positions of problematic taxa, and challenged current classificatory proposals. The total-evidence phylogeny of Megachilidae suggests that both Dioxyini and Aspidosmiini are extant relatives of the extinct tribes Ctenoplectrellini and Glyptapini. Previous authors placed all these tribes whitin either Anthiidini or Osmiini (*e.g*., Engel, 2001; Michener, 2007), but our analyses clearly show them in a well-supported clade, sister to the remaining Megachilinae. The relationship of Dioxyini with these fossil taxa is a new phylogenetic hypothesis for this distinctive taxon. All four of these tribes have aroliae, cleft pretarsal claws, and the vein 2m-cu basal to 2rs-m, but these features are also present in other Megachilinae. Aspidosmiini rendered Ctenoplectrellini paraphyletic in our analysis, and in that case, it would be appropriate to synonymize the former under the latter tribe. Gonzalez *et al*. (2012) discussed this option but ultimately decided to recognize it in its own tribe because of the limited number of scored characters for these fossils, which could have biased the results. Thus, we suggest recognizing Aspidosmiini until further analyses test these relationships using a larger number of characters for the fossil taxa.

The osmiines *Afroheriades* and *Pseudoheriades* are sister genera that form a well-supported clade. The morphological analysis of Gonzalez *et al*. (2012) placed them among the *Heriades* group of genera of Osmiini whereas the molecular analysis of Praz *et al*. (2008) placed them either as sister of either Anthidiini or Megachilini. Griswold (1985) also suggested a relationship with Megachilini based on the modified setae on the fifth sternum of the male. Our analyses consistently placed these two genera and Megachilini in a well-supported clade (Fig. 8). All other Osmiini clustered in another clade. Thus, in order to recognize a monophyletic Osmiini, we would need to either transfer them to Megachilini or distinguish them in their own tribe. Tranferring them to Megachilini weakens the recognition and diagnosis of this tribe because a morphological synapomorphy unambiguously present in that clade is unknown. In contrast, recognizing *Afroheriades* and *Pseudoheriades* in their own tribe highlights their distinctiveness while maintaining the current taxon concept for Megachilini. The tribe Osmiini will thus contain those taxa clustered in the other clade, which is sister to Megachilini + (*Afroheriades* + *Pseudoheriades*), except for *Ochreriades*. The phylogenetic position of *Ochreriades* has varied among morphological (Gonzalez *et al*., 2012), molecular (Praz *et al*., 2008), and our combined analyses (Fig. 1S, x). For example, it resulted among the *Heriades* group of genera, as sister to all of Megachilinae, or as sister to Megachilini and Osmiini. In our Bayesian analysis, *Ochreriades* clustered with the majority of osmiine (excluding *Afroheriades* + *Pseudoheriades*) in a clade with low support.

*Ochreriades* is morphologically distinctive among megachilids. It has yellow integumental markings as in the Anthidiini, a very long body with an elevated pronotum that surrounds the mesoscutum anteriorly, and long mouthparts that reach the tip of metasoma. Thus, *Ochreriades* is a taxon with distinctive features whose separation from Osmiini is desirable. Given this, herein we proposed two new tribes, Pseudoheriadini for *Afroheriades* + *Pseudoheriades*, and Ochreriadini for *Ochreriades* (see below). Our analyses also support the recognition of three subtribes or genus groups within Osmiini (Chelostomina Kirby, Heriadina Michener, and Osmiina Newman).

Neither our morphological analysis nor the total-evidence phylogeny supports the proposal of Trunz *et al*. (2016) of recognizing *M*. (*Heriadopsis*) and *M*. (*Matangapis*) at the generic level. In our analyses, *M*. (*Heriadopsis*) always clustered near *M*. (*Chelostomoides*), but the position of *M*. (*Matangapis*) changed from being part of the same clade with *M*. (*Heriadopsis*) to sister of *M*. (*Lophanthedon*) in the combined analysis. Thus, separating these taxa at the generic rank creates a large paraphyletic *Megachile s.l*.

Although sharing the presence of arolia with both *M*. (*Heriadopsis*) and *M*. (*Matangapis*), *Noteriades* never clustered with any of these taxa. It resulted either as the sister group of *Megachile s.l*. (EW analysis of morphological data, Fig. 9) or as sister of Megachilini (IW analyses of morphological data and combined analyses, Figs. 10, 11). Thus, our study further supports the placement of this group within Megachilini, as well as its recognition at the generic rank.

Michener (2007) suggested that *Coelioxys* might render *Megachile s.l*. paraphylectic because of its shared morphological features, particularly with *M.* (*Chelostomoides*). He also suggested that cleptoparasitism might have evolved independently in *Radoszkowskiana* and *Coelioxys*. Morphologically, *Radoszkowskiana* differs from *Coelioxys* in the short axilla, bare eyes, and the blunt metasoma of the male, which has a low transverse apical carina on T6 that is not divided into dorsal and ventral processes as in most *Coelioxys* [but is similar to that of males in *M.* (*Chelostomoides*)]. Some species of *Coelioxys* combine features of both genera. For example, *C.* (*Boreocoelioxys*) *funeraria* Smith and *C*. (*Liothyrapis*) *decipiens* Spinola have short axillae and bare eyes; also, the S6 of the female of *C*. (*Torridapis*) *torrida* Smith is broad and rounded, and entirely sclerotized as in *Radoszkowskiana* whereas it is elongated and pointed with a distinct median weakly sclerotized area in most *Coelioxys*. Furthermore, the mode of cleptoparasitism in *Radoszkowskiana* seems to fall within the known repertoire of parasitism of *Coelioxys* (Rozen and Kamel, 2007). Thus, *Radoszkowskiana* seems to be a *Coelioxys* despite the distinctive male characters. Our analyses consistently placed *Radoszkowskiana* as the sister group to *Coelioxys*, a relationship also recovered in the phylogenetic analyses of Rocha Filho and Parker (2016) and Trunz *et al*. (2016). However, the position of this clade of cleptoparasitic bees varied among our analyses.

In the morphological analysis under EW, this clade resulted as the sister group of *Noteriades* + Megachilini (Fig. 9), but it rendered *Megachile s.l*. paraphyletic when we analized the morphological data under IW and in combination with molecular data (Figs. 10, 11). Thus, our total-evidence phylogeny supports Michener’s (2007) hypothesis that *Coelioxys* renders *Megachile s.l*. paraphyletic, but it does not support his view of two independent origins of cleptoparasitism. Behavioral, morphological, and molecular data strongly indicate that cleptoparasitism evolved only once in Megachilini.

Most subgenera of *Megachile s.l*. fell into morphological groups previously associated with differences in nesting behavior. Basal branches included those subgenera of Group 2 that use mud or resins as nesting materials. These subgenera grouped in different clades whose taxonomic composition changed among analyses, except in a few cases [*e.g*., *M*. (*Maximegachile*) and *M*. (*Schizomegachile*) always resulted as sister groups]. Michener (2007: p. 553) discussed some of these relationships, which we mostly recovered in the morphological analysis under EW, but not under IW nor in the combined phylogeny. Thus, our analyses support the suspicion of Michener (2007) that Group 2 [*Chalicodoma sensu* Michener (1962) and Mitchell (1980)] is nonmonophyletic but it does not support the majority of his divisions or phylogenetic lines.

Taxa that exhibit LC behavior clustered in a large, more derived clade containing all subgenera of Group 1, and included *M*. (*Megella*), *M*. (*Mitchellapis*) (Group 2), and *M*. (*Creightonella*) (Group 3). These taxa combine some features that are typical of Group 1 and 2 and thus, they have been difficult to place with confidence in any group based on a few morphological features. The basal position *M*. (*Creightonella*) and *M*. (*Mitchellapis*) within the LC clade indicate that they retained some of the Group 2 features (chalicodiform body, male S8 with marginal setae) whereas the more derived position of *M*. (*Megella*) indicate the recurrence of some features of Group 2. Because it is commonly argued that the cost of a character gain is much higher than its loss, the reacquisition of characters makes some taxa, such as *M*. (*Megella*), difficult to place in a given taxonomic category. However, studies have documented the recurrence of complex structures, such as eyes and wings (*e.g*., West-Eberhard, 2003; Whiting *et al*., 2003). Thus, the gain of less complex structures, such as the marginal setae of S8 and arolia, seems even more plausible in Megachilini.

*Megachile* (*Chelostomoda*) is another group that combines features of Groups 1 and 2 of *Megachile s.l*. In the EW analysis of the morphological data, this subgenus was near *M*. (*Chelostomoides*) in Clade B, but it resulted as the sister group of LC bees when we analyzed this data set under IW and in combination with molecular data. The IW scheme shows that the characteristics of Group 2 (*e.g*., elongate body, terga with strong postgradular grooves, and S8 with marginal setae) exhibited by *M*. (*Chelostomoda*) are homoplastic features. The nesting biology of *M*. (*Chelostomoda*), *M*. (*Creightonella*) and *M*. (*Megella*), which make extensive use of leaf pieces, also supports their placement in Group 1; the biology of *M*. (*Mitchellapis*) is unknown.

Our results also recovered some major phylogenetic lines or groups of subgenera within the LC clade previously discussed by Michener (1965, 2007) and Mitchell (1980). Some of them, such as the *Creightonella* and *Pseudocentron* groups (Fig. 14), are distinct and easily recognizable by one or two morphological features; others lack distinct features. We briefly discussed some of them below.

**Fig 14.**
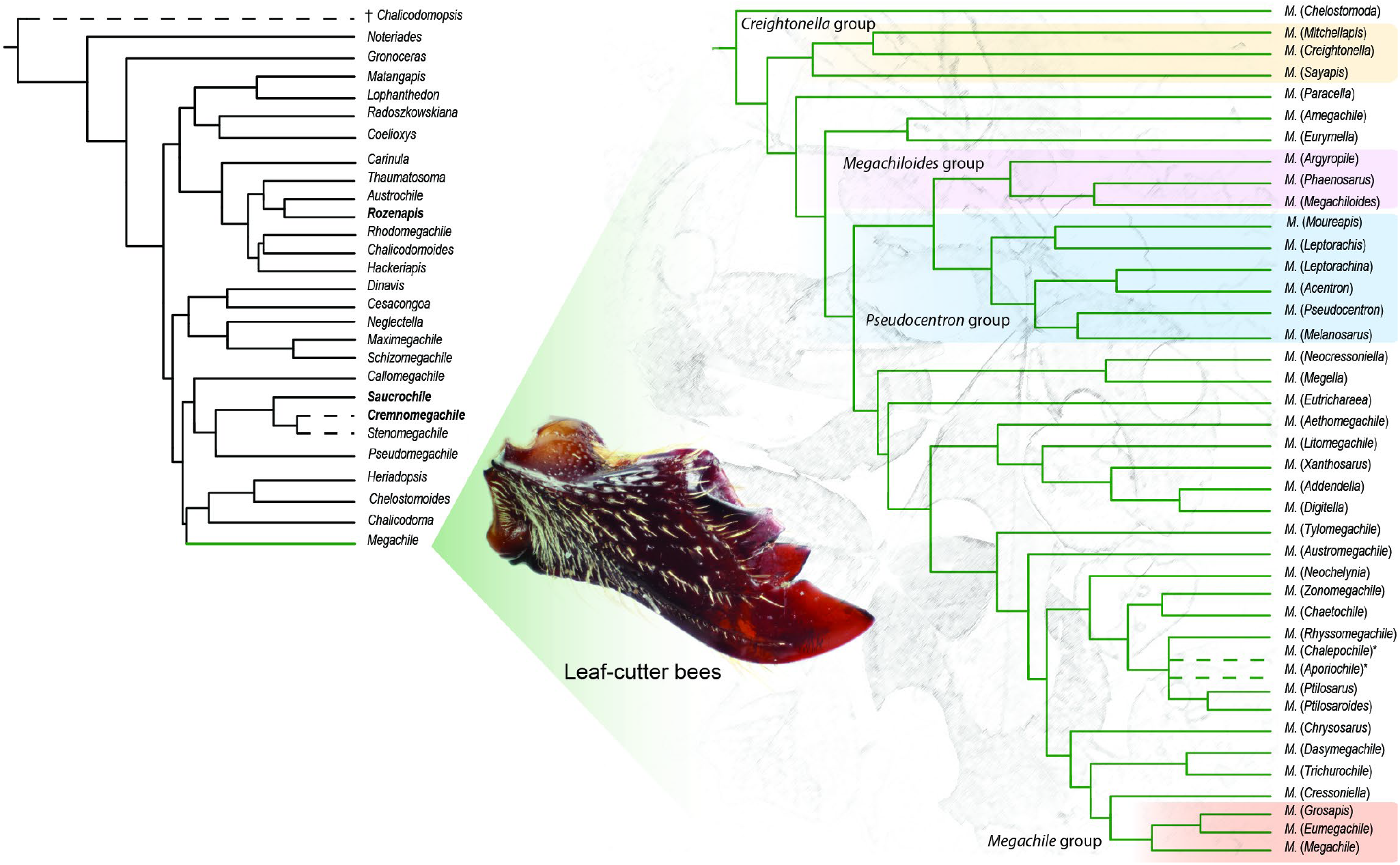
Summary of the proposed classification for Megachilini. Broken lines indicate uncertain position. New taxa described herein are bold faced. Gonzalez *et al*. (2018) recognized and described two new subgenera of *Megachile* after this work was completed (indicated with an asterisk) and were not included in the analyses. Groups of subgenera of *Megachile* highlighted in color are discussed in the text.

*Chrysosarus* group.—Mitchell (1980) included in this lineage the subgenera *M*. (*Chrysosarus*), *M*. (*Dactylomegachile*), *M*. (*Stelodides*), and *M*. (*Zonomegachile*). Based on the description (Raw, 2006) and photographs of the types, *M*. (*Austrosarus*) seems to belong here. Both type of interdental laminae are secondary lost in this group, except in *M*. (*Zonomegachile*). Gonzalez (2013) considered these taxa within a single subgenus, *M*. (*Chrysosarus*), but our analyses indicate that *M*. (*Zonomegachile*) does not belong to this group.

*Creightonella* group.—This lineage includes the subgenera *M*. (*Creightonella*), *M*. (*Mitchellapis*), *M*. (*Sayapis*), and *M*. (*Schrottkyapis*). The members of this group have a chalicodomiform body shape and odontogenic interdental laminae only. A remarkable feature of this lineage is the S6 of the female; at least in the species we examined for this study, it is elongated and with a membranous or weakly sclerotized pregradular area (visible only after dissection). Mitchell (1980) recognized this lineage under the generic name of *Eumegachile*; however, he also included the subgenera *M*. (*Eumegachile*) and *M*. (*Grosapis*) but separated *Creightonella* generically.

*Megachiloides* group.— Mitchell (1980) included *M*. (*Megachiloides*), *M*. (*Argyropile*), and three other names that Michener (2000, 2007) synonymized under *M*. (*Megachiloides*) or *M*. (*Xanthosarus*). Unlike most species of LC bees, members of this group appear to dig their own nest in the ground (Eickwort *et al*., 1981).

*Pseudocentron* group.—All members of this group of subgenera are primarily Neotropical in distribution; *M*. (*Acentron*), *M*. (*Leptorachis*), *M*. (*Melanosarus*), *M*. (*Moureapis*), and *M*. (*Pseudocentron*) are included here. Mitchell (1980) recognized this lineage and placed them in the genus *Pseudocentron*. The most distinctive character is the S6 of the female. It has at least the posterior half bare or nearly so, except for a subapical row of short setae, behind which there is a bare, smooth rim directed posteriorly.

Mitchell (1980) grouped the remaining subgenera of LC bees in two genera, *Megachile* and *Cressoniella*. In our analyses, these subgenera resulted in multiple clades but some taxa clustered as suggested by Mitchell. For example, he placed *M*. (*Ptilosarus*), *M*. (*Ptilosaroides*), *M*. (*Rhyssomegachile*), and *M*. (*Neochelynia*) in *Cressoniella*. We recovered these subgenera in the same clade but apart from the other subgenera included by Mitchell in his genus *Cressoniella*. Unlike the fossil tribes Ctenoplectrellini and Glyptapini, whose phylogenetic positions were consistent among analyses, that of the Dominican fossil *M. glaesaria* varied from being near *M*. (*Chelostomoides*) to be the sister group of all Megachilini. Interestingly, Engel (1999) discussed the possibility of both phylogenetic positions. Such instability might be the result of the low number of characters that we were able to score from this fossil (75 of 272).

### Monophyly of subgenera

Ten of the 57 subgenera of *Megachile s.l*. included in this study are monotypic (Table 3). The 19 subgenera containing more than one species but represented in our analyses by a single species are also likely monophyletic because each is morphologically uniform (*e.g*., *Maximegachile*, *Ptilosarus*). The monophyly of 18 subgenera was either strongly supported (*e.g*., *Pseudocentron*) or weakly supported but consistently suggested among analyses. Our analyses support the non-monophyly of several subgenera, which previous authors had already suspected or suggested (Michener, 2007; Trunz *et al*., 2016). Among the subgenera of Group 2, *M. biseta*, *M. decemsignata*, *M. memecylonae*, and *M. torrida* rendered *M*. (*Callomegachile*) paraphyletic; *M. ignita* and *M. heriadiformis* rendered *M.* (*Hackeriapis*); *M. muansae*, *M. cestella*, and *M. incana* rendered *M*. (*Pseudomegachile*); *M. dolichosoma* rendered *M*. (*Stenomegachile*); and *M*. *rugifrons* rendered *M*. (*Chelostomoides*). Among the clade of LC bees, *M. assumptionis* rendered *M*. (*Sayapis*) paraphyletic; *M. platystoma*, *M. eurymera*, *M. submetallica*, and *M. digiticauda* rendered *M*. (*Eutricharaea*); *M. fortis* and *M. addenda* rendered *M*. (*Xanthosarus*); *M. laeta* rendered *M*. (*Leptorachis*); *M. mitchelli* rendered *M*. (*Dasymegachile*); and *M. euzona* rendered *M*. (*Chrysosarus*).

**Table 3.** Monotypic, monophyletic, and non-monophyletic subgenera of *Megachile s.l*. * = subgenera represented by a single species in this study but they are likely monophyletic given their morphological uniformity; † = fossil taxa.

| Monotypic | Monophyletic | Likely monophyletic* | Nonmonophyletic |
| --- | --- | --- | --- |
| *Cesacongoa* | *Acentron* | *Alocanthedon* | *Callomegachile* |
| †*Chalicodomopsis* | *Aethomegachile* | *Austrochile* | *Chelostomoides* |
| *Eumegachile* | *Amegachile* | *Cestella* | *Chrysosarus* |
| *Grosapis* | *Argyropile* | *Chalicodomoides* | *Dasymegachile* |
| *Heriadopsis* | *Austromegachile* | *Cressoniella* | *Eutricharaea* |
| *Matangapis* | *Chalicodoma* | *Largella* | *Hackeriapis* |
| *Parachalicodoma* | *Chelostomoda* | *Lophanthedon* | *Leptorachis* |
| *Schizomegachile* | *Creightonella* | *Maximegachile* | *Pseudomegachile* |
| *Schrottkyapis* | *Gronoceras* | *Mitchellapis* | *Sayapis* |
| *Stelodides* | *Litomegachile* | *Moureapis* | *Stenomegachile* |
|  | *Megachile* | *Neocressoniella* | *Xanthosarus* |
|  | *Megachiloides* | *Platysta* |  |
|  | *Megella* | *Ptilosaroides* |  |
|  | *Melanosarus* | *Ptilosarus* |  |
|  | *Neochelynia* | *Rhyssomegachile* |  |
|  | *Paracella* | *Rhodomegachile* |  |
|  | *Pseudocentron* | *Thaumatoma* |  |
|  | *Tylomegachile* | *Trichurochile* |  |
|  |  | *Zonomegachile* |  |

In some cases, the recognition of highly derivative species at the subgeneric level render some taxa paraphyletic. For example, as Michener (2007) suspected, the monotypic subgenus *M*. (*Schrottkyapis*) renders *M*. (*Sayapis*) paraphyletic. A single putative synapomorphy supports such a relationship: S6 of the female with a nearly membranous pregradular area and a distinct invagination parallel to the lateral margin of the sternum (visible only after dissection). In other cases, such as in *M*. (*Eutricharaea*), *M*. (*Hackeriapis*), and *M*. (*Callomegachile*), current taxon concepts are broad and Michener (2007) proposed them as practical solutions to show relationships among diverse, poorly known groups. For example, when Michener (2007) synonymized various groups under *M*. (*Eutricharaea*), as he did for many other bees, he acknowledged the arbitrarity of his decision. He also highlighted morphological features that supported their relationships, which turned out to be homoplastic characters in our analyses (*e.g*., apical fasciae under scopa, T6 preapical carina toothed or denticulate and medially emarginated).

### Phylogenetic signal of morphological characters

The morphological character sets used in the phylogeny of Megachilini showed different levels of homoplasy (Table 1), as per RI value and percentage of unambiguous synapomorphic characters, but such differences were not statistically significant. Thus, these character sets were equally informative for the phylogeny. However, the analysis of female characters alone recovered Megachilini and several major lineages, unlike the analysis of male characters that resulted in a large polytomy. This does not mean that male characters were useless, but rather that our dataset was female-biased (Fig. 7).

Several authors have recognized important morphological features of taxonomic value in the male (*e.g*., Mitchell, 1980; Michener, 2007; Engel and Gonzalez, 2011; Gonzalez and Engel, 2012), which appear to show high rates of evolution, as they are sometimes highly variable within and among distinct phylogenetic lineages. For example, secondary sexual features such as the preapical carina of T6, mandibular projection, coxal spine, and modifications of front legs, are associated with particular strategies of mating system (Wittmann and Blochtein, 1995). The morphology of these structures are largely unexplored and, depending on the level of study and finer levels of character conceptualization, they might prove phylogenetically informative as in other group of bees (*e.g*., *Xylocopa* Latreille; Minckley, 1998). Equally unexplored is the female mandible. We conceptualized a number of characters for our level of study, but the mandible has a plethora of anatomical features with potential phylogenetic and taxonomic values at other levels of study. Michener and Fraser (1978) established terminology and homologies for the various structures of the bee mandible, but they only considered the body, not the apex. Among these ignored structures in the mandibular apex are interdental laminae, which as we have shown, are relevant for understanding the biology, taxonomy, and phylogeny of Megachilini.

### Classificatory proposals

Our phylogenetic results support the proposal of Gonzalez *et al*. (2012) in recognizing four subfamilies within Megachilidae, each with several tribes. It also clarifies the phylogenetic position of Aspidosmiini and Dioxyini, and provides evidence to resolve the long recognized non-monophyly of Osmiini. Based on these results, we propose to recognize two new tribes (Pseudoheriadini and Ochreriadini), thus, narrowing and strengthening the taxon concept of Osmiini (Table 4).

**Table 4.** Hierarchichal suprageneric classification of Megachilidae including two new tribes described in the text. Classification follows Engel (2005) and Gonzalez *et al*. (2012). † = fossil taxa.

| Taxon | Taxon (continued) |
| --- | --- |
| Subfamily Fideliinae Cockerell |  |
| Tribe Neofideliini Engel | Tribe Pseudoheriadini, **trib. nov.** |
| Tribe Fideliini Cockerell | Tribe Megachilini Latreille |
| Subfamily Pararhophitinae Popov | Tribe Ochreriadini, **trib. nov.** |
| Subfamily Lithurginae Newman | Tribe Osmiini Newman |
| Tribe †Protolithurgini Engel | Subtribe Chelostomina Kirby |
| Tribe Lithurgini Newman | Subtribe Heriadina Michener |
| Subfamily Megachilinae Latreille | Subtribe Osmiina Newman |
| Tribe †Glyptapini Cockerell | Tribe Anthidiini Ashmead |
| Tribe Dioxyini Cockerell | Subtribe Trachusina Robertson |
| Tribe †Ctenoplectrellini Engel | Subtribe Anthidiina Ashmead |
| Tribe Aspidosmiini Gonzalez *et al.* |  |

For Megachilini, we confirmed Michener’s (2007) suspicious that *Coelioxys* (as well as its sister group *Radoszkowskiana*) renders *Megachile s.l*. paraphyletic. In addition, our phylogenetic results suggest that recognizing *M*. (*Gronoceras*), *M*. (*Heriadopsis*), and *M*. (*Matangapis*) as genera distinct from *Megachile s.l*. further adds to the non-monophyly of this group. Thus, the current classification requires modification. We discuss three possible phylogeny-based solutions, but advocate for one that maximizes information storage and retrieval, memorability, and congruence with modern classification in other bee taxa (Table 5).

**Table 5.**
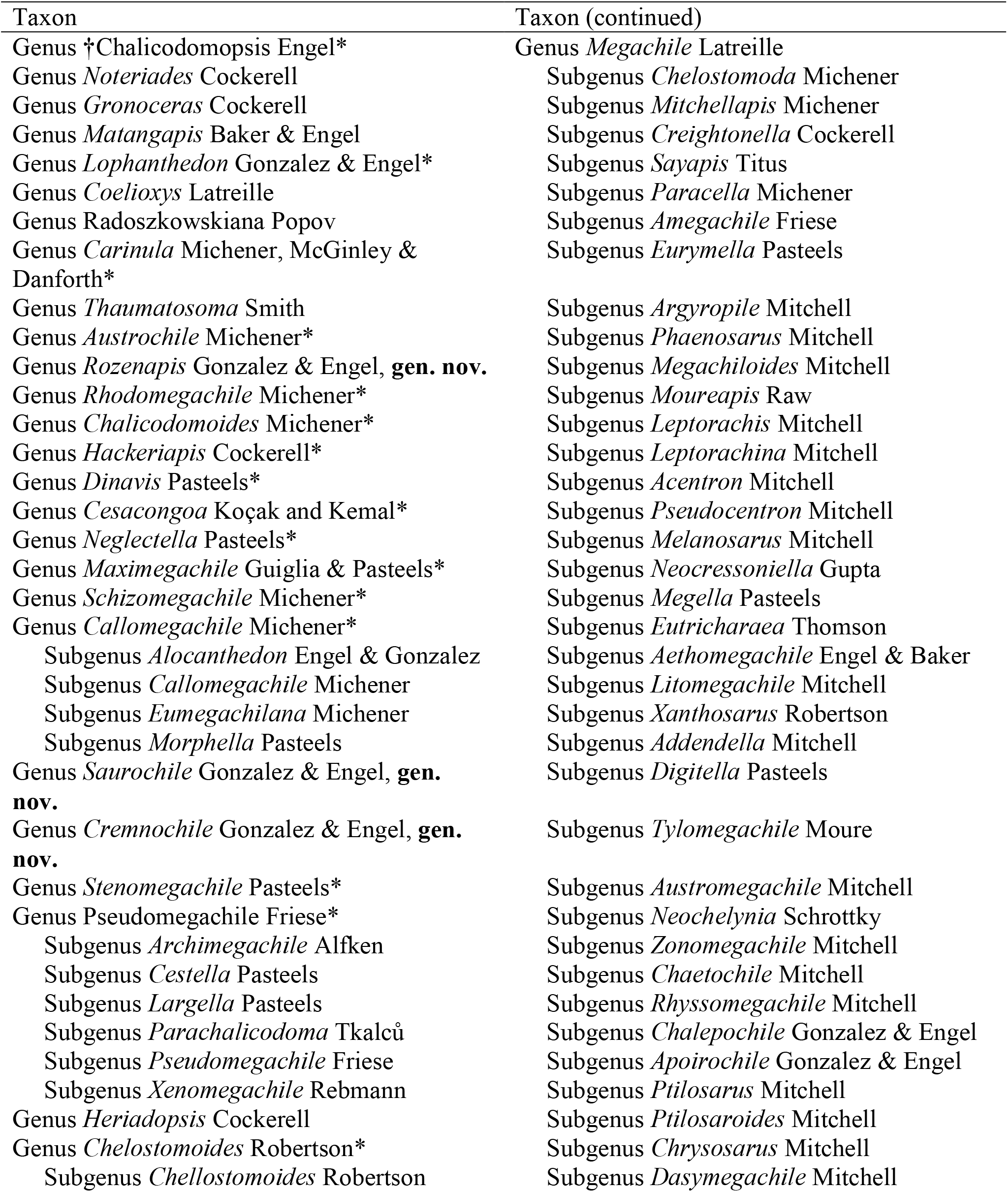
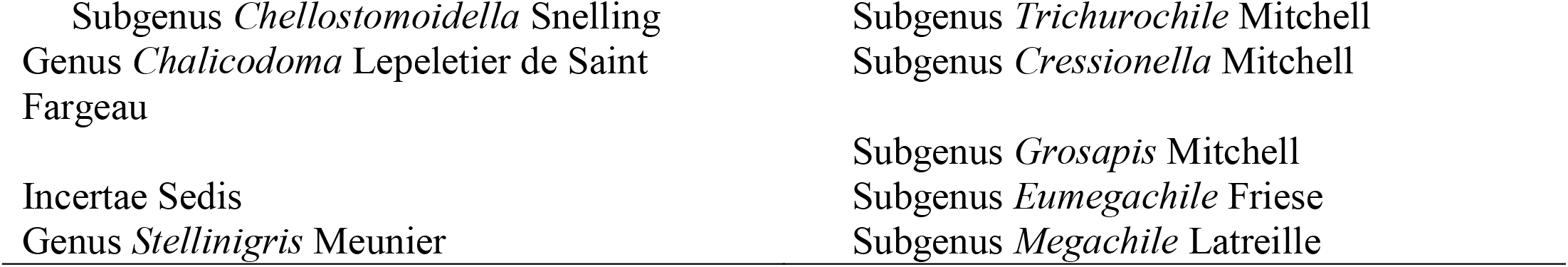
New classification of Megachilini following Proposal # 3 (see text). The list follows the order of taxa according to the phylogeny represented in Fig. 11. It does not includes the subgenera of *Coelioxys*. † = fossil taxa; * = new status

The first classificatory proposal is to recognize only two extant genera in Megachilini, *Noteriades* and *Megachile*. The cleptoparasitic genera *Coelioxys* and *Radoszkowskiana*, would be subgenera of *Megachile*. The fossil subgenus *M*. (*Chalicodomopsis*) could be treated either as a subgenus of *Megachile* given its position in the morphological analysis under EW or as a genus, as suggested in the IW and combined analyses.

The second proposal recognizes some of the subgenera of Group 2 at the generic rank, namely those taxa that clustered in a large clade with *Coelioxys* and *Radoszkowskiana*. All of these taxa are Old World in distribution and the majority of them are hoplitiform or heriadiform in body shape [*e.g*., *M*. (*Thaumatosoma*), *M*. (*Rhodomegachile*), *M*. (*Hackeriapis*)]. Thus, this proposal would recognize about 15 genera alongside *Megachile*, the latter including a mixture of Groups 1, 2, and 3.

The third proposal differs from the second in that *Megachile* would be restricted to the derived, well-supported clade that includes the LC bees only. This proposal would treat the remaining taxa at the generic level, thus recognizing about 20 genera total. Therefore, our proposals are somewhat similar to those discussed by Trunz *et al*. (2016), but differ in the number and identity of the taxa recognized at the generic level due to differences in the clade composition with our total-evidence phylogeny.

All three proposals imply new combinations of names and each proposal has practical advantages and disadvantages. An obvious advantage of retaining a large genus *Megachile s.l*., as in the first proposal, is that even with further knowledge of its phylogeny, the combinations of names created by the second and third proposals would not have to be accepted and perhaps, later, altered again. Phylogenies are always subject to change with the discovery of new taxa or the analysis of new morphological and molecular data. For example, features of immature stages might provide additional informative characters, but available information suggests little morphological variation in the major lineages of *Megachile s.l*. (Rozen *et al*., 2016). However, the first proposal also requires the inclusion of *Coelioxys* and *Radoszkowskiana* in *Megachile*, which would create more than 470 combinations of names and perhaps many new homonyms. *Megachile* would be an enormous genus with nearly 2000 species, more than 70 subgenera, and a wide range of biologies and morphologies. Such a retrograde classification is therefore highly undesirable.

Adopting the second or third proposal would not create as many new combinations of names as in the first proposal. In addition, authors initially proposed some taxa of Group 2 at the generic rank (*e.g*., *Chelostomoides*, *Gronoceras*, *Heriadopsis*, *Thaumatosoma*), which others subsequently treated first as subgenera of *Chalicodoma* and then of *Megachile*. Furthermore, because of the economic importance and worldwide distribution of Group 1, most published work has been done on members of this group rather than on Group 2 or 3. Thus, the new combinations of names resulting from treating the subgenera of Group 2 at the generic level would not have a major effect in the literature.

Recognizing some subgenera of Group 2 at the generic rank while others as subgenera of *Megachile*, as in the second proposal, would still make *Megachile* highly heterogeneous morphologically and biologically, redering the genus difficult to diagnose as well as differentiate from remaining Megachilini. However, the third proposal allows a more efficient retrieval of information and significantly improves the recognition and diagnosability of *Megachile* when the genus only consists of groups that cut leaves and developed interdental laminae. For example, recognizing *Megachile* in a narrower sense than in the second proposal would highlight the differences in nesting behavior and morphology among groups. This division may also encourage faster taxonomic revisions and comparative biological studies that would in turn increase our understanding of Megachilini.

The multigeneric classification of the third proposal migth seem like an extreme change, but upon inspection, is not. First, authors have previously recognized several subgenera within Group 2 at the genus rank, and the need for a multigeneric classification in Megachilini has repeatedly been voiced (*e.g*., Mitchell, 1980; Engel and Baker, 2006; Michener, 2007; Trunz *et al*., 2016). The problem at the time had been in choosing the best approach to picking which taxa to recognize at the genus-rank in the absence of a robust phylogenetic hypothesis. Second, the morphological differences among the subgenera of Group 2 are comparable or even greater than that among other genera of bees, including other megachilid tribes. For example, the morphological differences between the stingless bee genera *Trigona* Jurine and *Partamona* Schwarz (Apidae: Meliponini), or between the wool carder bee genera *Anthidium* Fabricius and *Afranthidium* Michener (Megachilidae: Anthidiini), seem trivial when we compare that between *M*. (*Hackeriapis*) and *M*. (*Chalicodoma*). Such a difference in the breadth of generic concepts among bee groups might be a reflection of the levels of taxonomic, phylogenetic, and morphological knowledge within each group. Third, many bee taxa now widely accepted as genera today, were treated in the past as subgenera of much larger genera, just like in the case of *Megachile s.l*. For example, Michener (1944) and Schwarz (1948) treated the more than 20 genera of Meliponini recognized today as subgenera of *Trigona*. For these reasons, we follow the third proposal given its practical adavantages, its hierarchichal arragement, and congruence with modern generic concepts of bees.

In addition to elevating the status of subgenera of Group 2 to the genus level following the third proposal, we also proposed three new genera for species that rendered some taxa paraphyletic (see below), as well as new synonymies and taxonomic arrangements. The genus *Callomegachile* Michener, as herein recognized, will consist of some distinct species groups that we consider as subgenera, including *Alocanthedon* Engel and Gonzalez. The five species of *Carinula* Michener *et al*. are more related to *Hackeriapis* than *Callomegachile*. The presence of translucent distal margins in the male terga reinforces their affinity to *Hackeriapis* and their recognition at the genus level. The species placed in *Parachalicodoma* Pasteels, *Largella* Pasteels, and *Cestella* Pasteels (new synonym) showed a close relationship with *Pseudomegachile* Friese, rendering the latter paraphyletic in some analyses. We considered them as subgenera of *Pseudomegachile* (see below). *Dinavis* Pasteels and *Negletella* Pasteels did not cluster with *Pseudomegachile* and thus we treat them as separated genera. *Chelostomoides rugifrons* (Smith) rendered *Chelostomoides* paraphyletic in all analyses. However, the morphology of both sexes is highly variable among the species of this group and we were not able to find a single morphological feature that consistently separated *C. rugifrons* from the remaining *Chelostomoides*. Thus, we retained this species in *Chelostomoides* despite its position in the analyses.

In the genus *Megachile*, we resurrect the following subgenera: *Eurymella* Pasteels and *Digitella* Pasteels from *Eutricharaea* Thomson; *Phaenosarus* Mitchell and *Addendella* Mitchell from *Xanthosarus* Robertson; *Leptorachina* Mitchell from *Leptorachis* Mitchell; and *Chaetochile* Mitchell from *Dasymegachile* Mitchell. We also newly synonymize *M*. (*Schrottkyapis*) Mitchell under *M*. (*Sayapis*) Titus. While Trunz *et al*. (2016) already established some of the changes indicated above (*e.g*., *Parachalicodoma* as a synonym of *Pseudomegachile*), some other authors (*e.g*., Durante and Abrahamovich, 2006; Moure *et al*., 2007) never adopted Michener’s (2007) classification and still recognized some of the subgenera (*e.g*., *Leptorachina*, *Chaetochile*) that we recovered as independent lineages in our analyses.

Finally, Trunz *et al*. (2016) proposed to synonymize *Grosapis* Mitchell and *Eumegachile* Friese under *Megachile s.str*., and *Paracella* Michener under *Anodoneutricharaea* Tkalců, the order of the latter synonym corrected by Praz (2017). We do not follow these changes because *Eumegachile* only rendered *Megachile* s.*str*. in the morphological analysis under EW, and both groups resulted as the sister group to *Megachile s.str*. in the combined analysis. Furthermore, *Grosapis* and *Eumegachile* are morphologically distinctive and their inclusion in *Megachile s.str*. would make the latter difficult to diagnose. Trunz *et al*. (2016) suggested the synonymy of *Paracella* under *Anodoneutricharaea* based on the phylogenetic position of *M. villipes* Morawitz, a species assigned to *Anodoneutricharaea*, a subgenus synonymized under *Eutricharaea* by Michener (2007). However, neither these authors nor we were able to include the type species of *Anodoneutricharaea* in the analyses, thus the phylogenetic position of this species remains uncertain.

## Conclusions and future directions

Our total-evidence tip dating analyses favor the hypothesis of a recent origin (15–25 Ma) for LC bees (Figs. 8, 10). Eocene trace fossil excissions are therefore not likely to be the result of the activity of bees within this particular clade, although the limited number of fossils included in our analyses may have affected our divergence time estimates. Our observations on the mandibular morphology of Glyptapini and Ctenoplectrellini, extinct lineages from Eocene Baltic amber, also indicate these taxa were unlikely to cut leaves (Figs. 13E–G). Thus, the identity of the Eocene LC insects remains elusive. Instead, the traces may represent the activity of as-of-yet unidentified stem-group Megachilini or Osmiini, as interdental laminae are not necessary for cutting leaves and the behavior certainly predated the origin of cutting structures.

Interdental laminae, the most distinctive and taxonomically significant feature of LC bees, developed from two different structures in the female mandible (Figs. 3B–H). One type of lamina developed from the tooth (odontogenic laminae) while the other from the fimbrial ridge (ctenogenic laminae). Odontogenic laminae, a putative synapomorphy for all LC bees, evolved first and exhibited more phenotypic plasticity than ctenogenic laminae (Fig. 14).

Our phylogenetic results also provided robust solutions to long-standing issues in the systematics of Megachilidae, namely the non-monophyly of Osmiini, phylogenetic position of Dioxyini, and the internal relationships of Megachilini. Our preferred multigeneric classification of Megachilini is not only consistent with the phylogeny, but it is also congruent with modern classifications of bees in terms of its hierarchy and breadth of generic taxon concepts.

The IW analyses of the morphological dataset of Megachilini, using a range of constant *k*-values calculated for average character fits (Appendix S5), support the effectiveness of this weighting scheme in recovering topologies congruent with total evidence phylogenies. As in Reemer and Ståhls (2012), we found that tree topologies obtained with *k*-values calculated for character fits near 70% are highly congruent with the total-evidence phylogeny.

Finally, our study provides a solid framework to formulate and address novel and interesting evolutionary questions regarding the LC behavior in bees. For example, is there a phylogenetic pattern between the type of interdental laminae and the plants used by LC bees? Do mandibular shape and interdental laminae correlate with any leaf feature (*e.g*., toughness) or any particular cutting and handling process? Are interdental laminae stronger and more resistant to abrasion when compared with each other as well as with teeth? Do interdental laminae contain heavy metals and halogens to increase hardness as in the mandible of other insects? Certainly, plants vary in leaf traits and the variable morphology of the mandible of LC bees suggest mechanical solutions to some functional problems. In LC ants, for example, the mandible and the LC behavior of species that cut edicot leaves are different from those that cut grasses (*e.g*., Camargo *et al*., 2015). In addition, the mandibular teeth of LC ants have heavy metals (*e.g*., Schofield *et al*., 2002), which increase their hardness and influence their ability to cut leaves. These aspects are unknown for LC bees, although there seems to be great variation within and among species in the degree and manner of leaf use. For example, a few records indicate that some species of *M*. (*Litomegachile*), *M*. (*Megachiloides*), *Megachile s. str*., and *M*. (*Xanthosarus*) use small circular pieces of leaves to make the bottom of a brood cell (Williams *et al*., 1986; Krombein and Norden, 1995). In other subgenera, such as *M*. (*Eutricharaea*), bees make the bottom of the cell by bending the leaf pieces from the cell cup (Medler, 1965; Kim, 1992). However, the nesting biology of the vast majority of species of *Megachile s.l*. remains unknown.

## Descriptions of new taxa

Family Megachilidae Latreille

Subfamily Megachilinae Latreille

Pseudoheriadini Gonzalez & Engel, **trib. nov.**

Type genus: *Pseudoheriades* Peters

### Diagnosis

This tribe can be readily separated from all other tribes of Megachilinae by the following combination of features: small body size (4.0—8.5 mm in length); heriadiform; maxillary palpi two-segmented; propodeum with basal area not marked posteriorly by a strong carina, but if present, it does not extends laterally behind propodeal spiracle; outer surfaces of pro- and mesotibiae without a distinct notch on distal margin; arolia present; female T6 with wide apical hyaline rim; male T7 large, exposed, quadrately surrounded by T6; male S3 with gradulus projecting into thin, basal hyaline lamella; male S5 with capitate discal setae.

### Description

♀. Preoccipital carina present (laterally in *Pseudoheriades*, dorsally in *Afroheriades*); clypeus little to no overhanging labral base; labrum not elongate, margin without fringe or apical tuft of setae; maxillary palpi two-segmented; mesoscutellum flat or slightly convex, not overhanging metanotum; metepisternum with dorsal carina or lamella (weakly present in *Afroheriades*); T6 with wide apical hyaline rim; S6 without lateral or apical projection. ♂. Metasoma with two or three sterna visible; T7 large, exposed, quadrately surrounded by T6; S3 with gradulus projecting into thin, basal hyaline lamella; S5 with capitate discal setae.

### Comments

This tribe contains at least 15 species (Griswold and Gonzalez, 2011; Ascher and Pickering, 2018) grouped in two Eastern Hemisphere genera, *Afroheriades* and *Pseudoheriades*. The first genus is restricted to the Cape Province of South Africa whereas the second is more widespread, occurring across Africa, the Middle East, and India. Griswold (1985) provided detailed descriptions and diagnostic features of both genera, some of which Griswold and Gonzalez (2011) illustrated.

Ochreriadini Gonzalez & Engel, **trib. nov.**

Type genus: *Ochreriades* Mavromoustakis

### Diagnosis

This tribe is readily separated from all other tribes of Megachilinae by the following combination of features: body very long and with yellow or ivory integumental markings; pronotum distinctly elevated and surrounding mesoscutum anteriorly; very long mouthparts, reaching the tip of metasoma.

### Description

♀. Clypeus no overhanging labral base; labrum not elongate, margin without fringe or apical tuft of setae; maxillary palpi three-segmented; metepisternum with dorsal carina or lamella; pronotum enlarged and surrounding mesoscutum anteriorly, practically eliminating omaular surface of mesepisternum and anterior surface of mesoscutum; mesoscutellum flat, on the same plane with metanonum and propodeal base, as seen in profile; T6 without wide apical hyaline rim; S6 without lateral or apical projection. ♂. Metasoma with six sterna visible; S2 and S3, each with disc swollen; S4 with dense pubescence on disc; S5 not emarginate, with branched or simple discal setae; T7 exposed, inferiorly directed.

### Comments

This tribe contains a single genus, *Ochreriades*, which consists of two species. *Ochreriades fasciatus* (Friese) occurs in the Middle East whereas *O. rozeni* Griswold occurs in Namibia, Africa (Griswold, 1994; Ascher and Pickering, 2018).

Tribe Megachilini Latreille

*Cremnomegachile* Gonzalez & Engel, **gen. nov.**

Type species: *Megachile dolichosoma* Benoist, 1962.

### Diagnosis

This genus resembles *Stenomegachile* Pasteels in the elongate, shiny female mandible, female hypostomal area toothed, and male preapical carina of T6 bilobed. It can easily separated by the shape of mesoscutum, which is midanteriorly projected and truncate, thus forming an anterior-facing area.

### Description

Small to moderate sized-bees (10.0–12.0 mm in body length). Integument shiny, with punctures coarse and spaced. Preoccipital border strongly carinate on gena; ocelloccipital distance distinctly greater than ocellocular distance. ♀. Mandible without interdental laminae, elongate, outer surface shiny, with apex about as broad as base, four-toothed, Mt4 on upper margin and clearly separated from Mt1–3, which are on distal margin; clypeus not covering base of labrum; labrum elongate, triangular, with distinct preapical protuberance bearing long, stiff tuft of setae; hypostomal carina with posterior portion ending in a tooth. Pronotal lobe with transverse lamella; mesoscutum flat on disc, midanteriorly projected and truncate, thus forming an anterior-facing area; mesoscutellum flat, not overhanging metanotum in dorsal view (Fig. 4C). Metasoma narrow, parallel-sided, with white apical fasciae and distinct postgradular grooves on T2–T4; sterna without apical fasciae beneath scopa; T6 straight (vertical) in profile. ♂. Antennal flagellum unmodified, F1 shorter than F2; mandible tridentate, without basal projection or tooth on lower margin; hypostomal carina unmodified, area behind mandible unmodified, without a projection or concavity; procoxa aspinose; pro- and mesotibiae and tarsi unmodified; metabasitarsus elongate, about 4.0× longer than broad; mesotibial spur present, articulated to mesotibia, about as long as apical width of mesotibia. T6 vertical in profile, with deep concavity above broad, medially emarginate preapical carina, distal margin without a distinct tooth or projection. T7 with preapical carina broadly rounded; S4 exposed, with punctation and vestiture similar to those of preceding sterna; S8 with marginal setae. Genital capsule elongate, 1.9× longer than wide; gonostylus straight or nearly so in ventral view, apically simple (not bifid), much narrower than base in lateral view, with long setae along its medial margin; volsella present, apically truncate.

### Etymology

The new genus-group name is a combination of of the Greek word, *kremnos*, meaning “overhanging wall”, in reference to the projected and anterior-facing surface of the mesoscutum, and *Megachile*.

### Comments

The genus is known from the type species only, which occurs in southern Madagascar (Pauly *et al*., 2001). In addition to the features indicated in the diagnosis, the male of *Stenomegachile* differs from that of *Cremnomegachile* in the four-toothed mandible (tridentate in *Cremnomegachile*), the hypostomal area, behind the mandible, which strongly projected into a tooth (unmodified in *Cremnomegachile*), and the pro- and mesotarsi expanded (normal in *Cremnomegachile*). The genital morphology is very different, particularly in the shape of the volsella, which is a narrow and apically notched (see Pasteels, 1965: p. 513). In the female of *Stenomegachile*, the mandible is more elongate and apically curved, and the labrum is long but parallel-sided. The hypostomal projection of *Stenomegachile* might not be homologous to that of *Cremnomegachile* because it is not part of the posterior portion of the hypostomal carina as in the latter genus.

*Rozenapis* Gonzalez & Engel, **gen. nov.**

Type species: *Megachile ignita* Smith, 1853.

### Diagnosis

This genus superficially resembles some robust species of *Hackeriapis* Cockerell with the terminal terga reddish and thus contrasting with the preceeding black terga. The female shares with *Austrochile* Michener a large, conspicuous midapical spine on S1 (absent in *Hackeriapis*), but it differs in the mandible. In *Austrochile*, the transverse ridge is strong and extends basally to merge with the acetabular carina whereas in *Rozenapis* such a ridge is entirely absent. The male differs from *Austrochile* in the absence of the midapical spine of S1 and the shape of T6, which has four equally distant teeth on its distal margin and a preapical carina that extends almost across the entire width of the tergum. In *Austrochile*, the spine of S1 is present, the preapical carina of T6 is restricted to the median third, and the median projections of the distal margin are closer than the distance from one of them to a lateral tooth. The male of *Rozenapis* differ from *Hackeriapis* (*sensu* King, 1994) in the impunctate distal margins of T2–T4, which are narrow and nearly concolor with discal areas (broad, distinctive, and hyaline in *Hackeriapis*). It also differs in the pretarsal claws, which lack a basal tooth (present in *Hackeriapis*).

### Description

Moderate sized-bees (12.0–15.0 mm in body length). Integument shiny, with punctures coarse and nearly contigous. Preoccipital border rounded, not carinate; ocelloccipital distance slightly longer than ocellocular distance in the female, much longer in the male. ♀. Mandible without interdental laminae, short, outer surface dulled without transverse ridge, with apex about as broad as base, four-toothed; clypeus barely covering base of labrum; labrum rectangular. Pronotal lobe with transverse carina; mesoscutellum not overhanging metanotum in dorsal view. Metasoma robust, parallel-sided, with white apical fasciae laterally only and weak postgradular grooves on basal terga; S1 with long, distinct midapical projection; sterna without apical fasciae beneath scopa; T6 gently convex in profile, slightly concave preapically. ♂. Antennal flagellum unmodified, F1 shorter than F2; mandible tridentate, without basal projection or tooth on lower margin; hypostomal area behind mandible unmodified, without a projection or concavity; procoxal spine small; pro- and mesotibiae and tarsi slightly expanded; metabasitarsus elongate, about 4.0× longer than broad; mesotibial spur present, articulated to mesotibia, about as long as apical width of mesotibia. T6 vertical in profile, with deep concavity above broad, medially emarginate preapical carina, distal margin with four small, equidistant teeth or projections. T7 with preapical carina slightly projecting medially; S4 apically exposed, with punctation and vestiture similar to those of preceding sterna; S8 with marginal setae. Genital capsule elongate, 1.4× longer than wide; gonostylus straight or nearly so in ventral view, apically simple, truncate, much broader than base in lateral view, with short setae along its medial margin; volsella present, apically notched.

### Etymology

This new genus-group name is a patronymic honoring Dr. Jerome G. Rozen, Jr., American Museum of Natural History, for his significant contributions to biology and systematics of bees.

### Comments

This genus resulted as the sister group of *Austrochile* in our analyses. Only the type species is known, which Michener (1965) listed in *Hackeriapis* as a member of species of group A.

*Saucrochile* Gonzalez & Engel, **gen. nov.**

Type species: *Megachile heriadiformis* Smith, 1853.

### Diagnosis

This genus is most similar to *Hackeriapis* (*sensu* King, 1994). It differs in the pretarsal claws, which lack of a basal tooth, and in the distal margins of male T2–T4, which are punctate and concolor with discal areas. In *Hackeriapis*, the pretarsal claws have a distinct basal tooth and the distal margins of male T2–T4 are impunctate, broad, and hyaline. In addition, the pronotal lobe is distinctly carinate or lamellate, at least dorsally, in *Hackeriapis* (rounded in *Saucrochile*).

### Description

Small sized-bees (8.0–11.0 mm in body length). Integument shiny, with punctures coarse and spaced. Preoccipital border rounded, not carinate; ocelloccipital distance much longer than ocellocular distance. ♀. Mandible without interdental laminae, elongate, outer surface shiny, with sparse punctures, outer ridge weak, extending basally to acetabular carina, three teeth on distal margin; clypeus not covering base of labrum; labrum elongate, parallel-sided, without preapical protuberance. Pronotal lobe without transverse carina or lamella; mesoscutellum flat, not overhanging metanotum in dorsal view. Metasoma elongate, parallel-sided, with white apical fasciae and strong postgradular grooves on basal terga; sterna without apical fasciae beneath scopa; T6 gently convex in profile. ♂. Antennal flagellum unmodified, F1 shorter than F2; mandible tridentate, without basal projection or tooth on lower margin; hypostomal area behind mandible unmodified, without a projection or concavity; procoxal spine small; pro- and mesotibiae and tarsi unmodified; metabasitarsus elongate, about 4.0× longer than broad; mesotibial spur present, articulated to mesotibia, about as long as apical width of mesotibia. T6 vertical in profile, with weak concavity above narrow, medially emarginate preapical carina, distal margin with four small, equidistant teeth or projections. T7 with preapical carina slightly projecting medially; S4 hidden, with punctation and vestiture different to those of preceding sterna; S8 with marginal setae. Genital capsule elongate, about 2.0× longer than wide; gonostylus straight or nearly so in ventral view, slightly narrower basally in lateral view, apically simple, with short setae along its medial margin; volsella present, apically notched.

### Etymology

The new genus-group name is a combination of of the Greek words, *saukros*, meaning “graceful”, and in reference to the general elegant aspect of this group, and *chile*, meaning “tooth”.

### Comments

Only the type species is known, which Michener (1965) listed it in *Hackeriapis* as a member of species of group A.

### Key to the extant tribes of the Megachilinae

(Modified from Michener, 2007)

1. Metanotum with median spine or tubercle (except in *Allodioxys* and *Ensliniana*); mandible of female slender apically, bidentate, similar to that of male; pronotum (except in *Prodioxys*) with prominent obtuse or right-angular dorsolateral angle, below which a vertical ridge extends downward; sting and associated structures greatly reduced (scopa absent) …………………‥ Dioxyini
—. Metanotum without median spine or tubercle; mandible of female usually wider apically, with three or more teeth, except rarely bidentate when mandible is greatly enlarged and porrect and clypeus is also modified; pronotum with dorsolateral angle weak or absent (or produced to a tooth in some *Chelostoma* but without vertical ridge below it); sting and associated structures well developed …………………‥ 2
2(1). Pterostigma less than twice as long as broad, inner margin basal to vein r-rs usually little if any longer than width, rarely about 1.5 times width; pretarsal claws of female cleft or with an inner tooth (except in *Trachusoides*); body commonly with yellow or ivory integumental marks …………………‥ 3
—. Pterostigma over twice as long as broad, inner margin basal to vein r-rs longer than width; pretarsal claws of female simple (except in *Osmia* subgenus *Metalinella*, Palaearctic); body without yellow or white integumental marks, except in Ochreriadini …………………‥ 4
3(2). Outer surface of metatibia with long setae forming a distinct scopa; prestigma much more than twice as long as broad; preaxilla, below posterolateral angle of mesoscutum, sloping and with small patch of setae, these as long as those of adjacent sclerites …………………‥ Aspidosmiini
—. Outer surface of metatibia usually with abundant simple bristles, not forming a distinct scopa; prestigma commonly short, usually less than twice as long as broad; preaxilla vertical, smooth and shining, usually without setae …………………‥ Anthidiini
4(2). Body distinctly elongate with enlarged pronotum surrounding mesoscutum anteriorly, thus practically eliminating omaular surface of mesepisternum and anterior surface of mesoscutum; body with yellow or ivory integumental markings at least on metasoma …………………‥ Ochreriadini, **trib. nov.**
—. Body not as elongate and slender as above, pronotum not enlarged nor surrounding mesoscutum anteriorly, mesepisternum with distinct omaular surface, mesoscutum; body without yellow or white integumental marks …………………‥ 5
5(4). Outer surfaces of pro- and mesotibiae apically with an acute angle (usually produced into a spine) and distinct notch anteriorly; male T6 with preapical carina often present; arolia normally absent, except in a few tropical Old World taxa (*Noteriades*, *Matangapis* and *Heriadopsis*); body nonmetallic or nearly so …………………‥ Megachilini
—. Outer surfaces of pro- and mesotibiae apically without an acute angle or spine and lacking distinct notch anteriorly; male T6 without preapical carina; arolia present; body sometimes metallic green, blue, or brassy …………………‥ 6
6(5). Maxillary palpus with two palpomeres; propodeum with basal area not marked posteriorly by a strong carina, if present, it does not extend laterally behind propodeal spiracle (female: T6 with wide apical hyaline rim, S1 with slender, erect spine, posterolateral angle of mesoscutum with marginal ridge rounded or carinate, if rounded, with dense patch of long setae laterally); male T7 large, exposed, quadrately surrounded by T6; male S5 with modified discal setae …………………‥ Pseudoheriadini, **trib. nov.**
—. Maxillary palpus with at least with three palpomeres; propodeum with basal area not marked posteriorly by carina or if present, extending laterally behind propodeal spiracle; male T7 small, usually hidden, not quadrately surrounded by T6; male S5 with branched or simple discal setae …………………‥ Osmiini

## Acknowledgements

We thank the late Charles D. Michener for his many years of advice, encouragement, and comments on early drafts of this manuscript. Amy R. Comfort, Mabel Alvarado, Laura C.V. Breitkreuz, and two anonymous reviewers provided helpful comments and suggestions that improved this work. We also thank the curators, collection managers, and staff of the collections we visited or from which we borrowed specimens, and particularly Zachary H. Falin and Jennifer C. Thomas for their constant help throughout the project. The participation of V.H.G. was partially supported by National Science Foundation’s REU program (DBI-1560389), while that of G.T.G by a NIH IRACDA postdoctoral fellowship (5K12GM064651), and M.S.E. by NSF DBI-0096905 and DBI-1057366. This is a contribution of the Division of Entomology, University of Kansas Natural History Museum.

## References

Agnarsson, I., Coddington, J.A. 2008. Quantitative test of primary homology. Cladistics 24, 51–61.

Arcila, D., Pyron, R.A., Tyler, J.C., Ortí, G., Betancur-R, R. 2015. An evaluation of fossil tip-dating versus node-age calibration in tetraodontiform fishes (Teleostei: Percomorphaceae). Mol. Phylogenetics. Evol. 82, 131–145.

Ascher, J.S., Pickering, J. 2018. Discover Life bee species guide and world checklist (Hymenoptera: Apoidea: Anthophila). Draft 50. Available from http://www.discoverlife.org/mp/20q?guide=Apoidea_species (accessed March 31, 2018)

Baker, D.B., Engel, M.S. 2006. A new subgenus of Megachile from Borneo with arolia (Hymenoptera: Megachilidae). Am. Mus. Novit. 3505, 1–12.

Banaszak, J., Romasenko, L. 1998. Megachilid bees of Europe (Hymenoptera, Apoidea, Megachilidae). Bydgoszcz, Poland: Pedagogical University of Bydgoszcz. 239 pp.

Bänziger, H., Boongird, S., Sukumalanand, P., Bänziger, S. 2009. Bees (Hymenoptera: Apidae) that drink human tears. J. Kansas Entomol. Soc. 82, 135–150.

Brauns, H. 1926. V. Nachrag zu “Friese, Bienen Afrikas”. Zool. Jahrb. Abt. System. Geog. Biol. Tiere. 52, 187–230.

Camargo, R.S., Hastenreiter, I.N., Forti, L.C., Lopes, J.F.S. 2015. Relationship between mandible morphology and leaf preference in leaf-cutting ants (Hymenoptera: Formicidae). Rev. Colomb. Entomol. 41, 241–244.

Cane, J.H. 2004. Exotic nonsocial bees (Hymenoptera: Apiformes) in North America: Ecological implications, pp. 113–116. In: Strickler KL, Cane JH (Eds.). For non-native crops, whence pollinators of the future? Thomas Say Publications in Entomology, Lanham, MD.

Cardinal, S., Danforth, B.N. 2013. Bees diversified in the age of eudicots. Proc. R. Soc. Lond. [Biol]. 280, 20122686. https://doi.org/10.1098/rspb.2012.2686.

Castresana, J. 2000. Selection of conserved blocks from multiple alignments for their use in phylogenetic analysis. Mol. Biol. Evol. 17, 540–552.

Cockerell, T.D.A. 1907. Descriptions and records of bees–XV. Ann. Mag. Nat. Hist. 7, 59–68.

Cockerell, T.D.A. 1922. Descriptions and records of bees–XCV. Ann. Mag. Nat. Hist. 9, 265–269.

Dalla Torre, C.G. 1896. Catalogus hymenopterorum hucusque descriptorum systematicus et synonymicus. Vol. X. Apidae. Leipzig: Engelmann.

Dos Reis, M., Yang, Z. 2013. The unbearable uncertainty of Bayesian divergence time estimation. J. Syst. Evol. 51, 30–43.

Duckworth, R.A. 2009. The role of behavior in evolution: a search for mechanism. Evol. Ecol. 23, 513–531.

Durante, S., Abrahamovich, A. 2006. Redescription of Chaetochile as subgenus of Megachile (Hymenoptera, Megachilidae). T. Am. Entomol. Soc. 132, 103–109.

Durante, S., Cabrera, N. 2009. Cladistic analysis of Megachile (Chrysosarus) Mitchell and revalidation of Megachile (Dactylomegachile) Mitchell (Hymenoptera, Megachilidae). Zootaxa 2284, 48–62.

Eardley, C. 2012. A taxonomic revision of the southern African species of dauber bees in the genus Megachile Latreille (Apoidea: Megachilidae). Zootaxa 3460, 1–139.

Eardley, C. 2013. A taxonomic revision of the southern African leaf-cutter bees, Megachile Latreille sensu stricto and Heriadopsis Cockerell (Hymenoptera: Apoidea: Megachilidae). Zootaxa 3601, 1–133.

Eickwort, G.C., Matthews, R.W., Carpenter, J. 1981. Observations on the nesting behavior of Megachile rubi and M. texana with a discussion of the significance of soil nesting in the evolution of megachilid bees (Hymenoptera: Megachilidae). J. Kansas Entomol. Soc. 54, 557–570.

Engel, M.S. 1999. Megachile glaesaria, the first megachilid bee fossil from amber (Hymenoptera: Megachilidae). Am. Mus. Novit. 3276, 1–13.

Engel, M.S. 2000. A new interpretation of the oldest fossil bee (Hymenoptera: Apidae). Am. Mus. Novit. 3296, 1–11.

Engel, M.S. 2001. A monograph of the Baltic amber bees and evolution of the Apoidea (Hymenoptera). Bull. Am. Mus. Nat. Hist. 259, 1–192.

Engel, M.S. 2005. Family-group names for bees (Hymenoptera: Apoidea). Am. Mus. Novit. 3476, 1–36.

Engel, M.S. 2007. Ferdinand Anatole Meunier and the destruction of his Hymenoptera collections. Entomol. Gaz. 58, 183–184.

Engel, M.S., Baker, D.B. 2006. A remarkable new leaf-cutter bee from Thailand. Beitr. Entomol. 56, 69–74.

Engel, M.S., Gonzalez, V.H. 2011. Alocanthedon, a new subgenus of Chalicodoma from Southeast Asia (Hymenoptera, Megachilidae). ZooKeys 101, 51–8.

Engel, M.S, Perkovsky, E.E. 2006. An Eocene bee in Rovno amber, Ukraine (Hymenoptera: Megachilidae). Am. Mus. Novit. 3506, 1–12.

Friese, H. 1898. Species aliquot novae vel minus cognitae generis Megachile Latr. (et Chalicodoma Lep.). Természetrajzti Füz. 21, 198–202.

Friese, H. 1899. Die Bienen Europa’s (Apidae europaeae) nach ihren Gattungen, Arten und Varietäten auf vergleichend morphologisch-biologischer Grundlage. Theil V: Solitäre Apiden: Genus Lithurgus, Genus Megachile (Chalicodoma). C. Lampe, Innsbruck: Lampe. Vol. 3, 228 pp.

Friese, H. 1909. Die Bienen Afrikas nach dem Stande unserer heutigen Kenntnisse. In: Schultz L (Ed.), pp. 83–476, pls. ix-x, Zoologische und Anthropologische Ergebnisse einer Forschungsreise im westlichen und zentralen Südafrika ausgefuhrt in den Jahren 1903-1905, Band 2, Lieferung 1, X Insecta (ser. 3) [Jenaische Denkschriften Vol. 14]. Jena: Fischer.

Friese, H. 1911a. Apidae I. Megachilinae. Das Tierreich. Lieferung 28. Berlin, Germany: Friedländer.

Friese, H. 1911b. Nachtrag zu “Bienen Afrikas.” Zoologische Jahrbücher, Abteilung für Systematik, Geographie und Biologie der Tiere 30: 651–670.

Gerstaecker, A. 1869. Beiträge zur näheren Kenntniss einiger Bienen-Gattugen. Entomol. Z. 30, 315–367.

Goloboff, P.A. 1993. Estimating character weights during tree search. Cladistics 9, 83–91.

Goloboff, P.A. 2008. Calculating SPR distances between trees. Cladistics 24, 591–597.

Goloboff, P.A., Farris, J., Nixon, K. 2003a. T.N.T. Tree analysis using new technology. Program and documentation. Available at: http://www.zmuc.dk/public/phylogeny/tnt/

Goloboff, P.A., Farris, J.S., Källersjö, M., Oxelman, B., Ramírez, M.J., Szumik, C.A. 2003b. Improvements to resampling measures of group support. Cladistics 19, 324–332.

Goloboff, P.A., Torres, A., Arias, J.S. 2017. Weighted parsimony outperforms other methods of phylogenetic inference under models appropiate for morphology. Cladistics 0, 1–31. https://doi.org/10.1111/cla.12205

Gonzalez, V.H. 2008. Phylogeny and classification of the bee tribe Megachilini (Hymenoptera: Apoidea, Megachilidae), with emphasis on the genus Megachile. Doctoral dissertation. Lawrence, Kansas: University of Kansas. 274 pp.

Gonzalez, V.H. 2013. Taxonomic comments on Megachile subgenus Chrysosarus (Hymenoptera: Megachilidae). J. Melittol. 5, 1–6.

Gonzalez, V.H., Engel, M.S. 2012. African and Southeast Asian Chalicodoma (Hymenoptera: Megachilidae): new subgenus, new species, and notes on the composition of Pseudomegachile and Largella. Ann. Zool. 62, 599–617.

Gonzalez, V.H., Engel, M.S., Hinojosa, I.A. 2010. A new species of Megachile from Pakistan, with taxonomic notes on the subgenus Eutricharaea (Hymenoptera: Megachilidae). J. Kans. Entomol. Soc. 83, 58–67.

Gonzalez, V.H., Griswold, T. 2013. Wool carder bees of the genus Anthidium in the Western Hemisphere (Hymenoptera: Megachilidae): diversity, host plant associations, phylogeny, and biogeography. Zool. J. Linnean Soc. 168, 221–425.

Gonzalez, V.H., Griswold, T., Engel, M.S. 2013. Obtaining a better taxonomic understanding of native bees: where do we start?. Syst. Entomol. 38, 645–653.

Gonzalez, V.H., Griswold, T., Engel, M.S. 2018. South American leaf-cutter bees (genus Megachile) of the subgenera Rhyssomegachile and Zonomegachile, with two new subgenera (Hymenoptera: Megachilidae). Bull. Am. Mus. Nat. Hist. 425, 1–73.

Gonzalez, V.H., Griswold, T., Praz, C.J., Danforth, B.N. 2012. Phylogeny of the bee family Megachilidae (Hymenoptera: Apoidea) based on adult morphology. Syst. Entomol. 37, 261–286.

Griswold, T.L. 1985. A generic and subgeneric revision of the Heriades genus-group. Doctoral Dissertation. Logan, Utah: Utah State University. xii + 165 pp.

Griswold, T.L. 1994. A review of Ochreriades (Hymenoptera: Megachilidae: Osmiini). Pan-Pac. Entomol. 70, 318–321.

Griswold, T.L., Gonzalez, V.H. 2011. New species of the Eastern Hemisphere genera Afroheriades and Noteriades (Hymenoptera, Megachilidae), with keys to species of the former. ZooKeys 159, 65–80.

Heath, T.A., Huelsenbeck, J.P., Stadler, T. 2014. The fossilized birth-death process for coherent calibration of divergence-time estimates. Proc. Natl. Acad. Sci. U.S.A., 111, E2957–E2966. https://doi.org/10.1073/pnas.1319091111

Horne, M. 1995. Leaf area and toughness: effects on nesting material preference of Megachile rotundata (Hymenoptera: Megachilidae). Ann. Entomol. Soc. Am. 88: 868–875.

Huelsenbeck, J.P., Larget, B., Alfaro, M.E. 2004. Bayesian phylogenetic model selection using reversible jump Markov Chain Monte Carlo. Mol. Biol. Evol. 21, 1123–1133.

Kalyaanamoorthy, S., Minh, B.Q., Wong, T.K.F., von Haeseler, A., Jermiin, L.S. 2017. ModelFinder: fast model selection for accurate phylogenetic estimates. Nature Methods 14, 587–589. https://doi.org/10.1038/nmeth.4285

Kambli, S.S., Aiswarya, M.S., Manoj, K., Varma, S., Asha, G., Rajesh, T.P., Sinu, P.A. 2017. Leaf foraging sources of leafcutter bees in a tropical environment: implications for conservation. Apidologie 48, 473–482.

Katoh, K. Toh, H. 2008. Improved accuracy of multiple ncRNA alignment by incorporating structural information into a MAFFT-based framework. BMC Bioinformatics 9, 212. https://doi.org/10.1186/1471-2105-9-212

Katoh, K., Standley, D.M. 2013. MAFFT Multiple Sequence Alignment Software Version 7: Improvements in performance and usability. Mol. Biol. Evol. 30, 772–780. https://doi.org/10.1093/molbev/mst010

Kim, J. 1992. Nest dimensions of two leaf-cutter bees (Hymenoptera: Megachilidae). Ann. Entomol. Soc. Am. 85, 85–90.

Kim, Y-H., Ahn, K-J. 2016. Phylogeny of the Homalotina (Coleoptera: Staphylinidae: Aleocharinae) based on morphology. Syst. Entomol. 41, 323–338.

King, J. 1994. The bee family Megachilidae (Hymenoptera: Apoidea) in Australia, I. Morphology of the genus Chalicodoma Lepeletier, and a revision of the subgenus Hackeriapis Cockerell. Invertebr. Taxon. 8, 1373–1419.

Krombein, K.V., Norden, B.B. 1995. Notes on the behavior and taxonomy of Megachile (Xeromegachile) brimleyi Mitchell and its probable cleptoparasite, Coelioxys (Xerocoelioxys) galactiae Mitchell (Hymenoptera: Megachilidae). Proc. Entomol. Soc. Wash. 97, 86–89.

Lanfear, R., Calcott, B., Ho, S., Guindon, S. 2012. PartitionFinder: combined selection of partitioning schemes and substitution models for phylogenetic analysis. Mol. Biol. Evol. 29, 1695–1701. https://doi.org/whttp://dx.doi.org/10.1093/molbe/mss020

Lanfear, R., Frandsen, P.B., Wright, A.M., Senfeld, T. Calcott, B. 2016. PartitionFinder 2: new methods for selecting partitioned models of evolution for molecular and morphological phylogenetic analyses. Mol. Biol. Evol. 34, 772–773. https://doi.org/10.1093/molbev/msw260

Lapiedra, O., Sol, D., Carranza, S., Beaulieu, J.M. 2013. Behavioural changes and the adaptive diversification of pigeons and doves. Proc. R. Soc. B. 280, 20122893.

Laroca, S., Corbella, E., Varela, G. 1992. Biologia de Dactylomegachile affabilis (Hymenoptera, Apoidea): I. Descrição do ninho. Acta Biol. Paran. 21, 23–29.

Latreille, P.A. 1802. Histoire Naturelle des Fourmis, et recueil de memoires et d’observations sur les abeilles, les araignées, les faucheurs, et autres insectes. Paris: Crapelet, xvi+445 pp

Lepeletier de Saint-Fargeau, A.L.M. 1841. Histoire Naturelle des Insectes–Hyménoptères. Paris: Roret. Vol. 2, 1–680.

Litman, J.R., Danforth, B.N., Eardley, C.D., Praz, C.J. 2011. Why do leafcutter bees cut leaves? New insights into the early evolution of bees. Proc. R. Soc. Lond. [Biol] 278, 3593–3600.

MacIvor, J.S. 2016. DNA barcoding to identify leaf preference of leafcutting bees. R. Soc. Open Sci. 3, 150623.

Maddison, W.P. Maddison, D.R. 2018. Mesquite: a modular system for evolutionary analysis. Version 3.40. http://mesquiteproject.org [accessed on Mar. 1, 2018]

Marín, M.A., Peña, C., Uribe, S.I., Freitas, A.V.L. 2017. Morphology agrees with molecular data: phylogenetic affinities of Euptychiina butterflies (Nymphalidae: Satyrinae). Syst. Entomol. 42, 768–785.

Medler, J.T. 1965. A note on Megachile mendica Cresson Say in trap-nests in Wisconsin (Hymenoptera: Megachilidae). Proc. Entomol. Soc. Wash. 67, 113–116.

Meunier, F. 1888. Megachillidae [sic]. Nat. Sicil. 7, 152.

Michener, C.D. 1944. Comparative external morphology, phylogeny, and a classification of the bees. Bull. Am. Mus. Nat. Hist. 82, 151–326.

Michener, C.D. 1953. The biology of a leafcutter bee (Megachile brevis) and its associates. Univ. Kans. Sci. Bull. 35, 1659–1748.

Michener, C.D. 1962. Observations on the classification of the bees commonly placed in the genus Megachile (Hymenoptera: Apoidea). J. New York Entomol. Soc. 70, 17–29.

Michener, C.D. 1965. A classification of the bees of the Australian and South Pacific regions. Bull. Am. Mus. Nat. Hist. 130, 1–362.

Michener, C.D. 1983. The classification of the Lithurginae (Hymenoptera: Megachilidae). Pan-Pac. Entomol. 59, 176–187.

Michener, C.D. 1996. The first South African dioxyine bee and a generic review of the tribe Dioxyini. Contributions on Hymenoptera and Associated Insects Dedicated to Karl V. Krombein, Memoirs of the Entomological Society of Washington: Vol. 17 (ed. by B.B. Norden and A.S. Menke), pp. 142–152. Entomological Society of Washington, Washington, District of Columbia.

Michener, C.D. 2000. The Bees of the World. Baltimore, Maryland: Johns Hopkins University Press. 913 pp.

Michener, C.D. 2007. The Bees of the World. Baltimore, Maryland: Johns Hopkins University Press. 2nd Edition, 953 pp.

Michener, C.D., Fraser, A. 1978. A comparative anatomical study of mandibular structure in bees. Univ. Kans. Sci. Bull. 51, 463–482.

Michez, D., Vanderplanck, M., Engel, M.S. 2012. Fossil bees and their plant associates. Pp. 103–106. In Patiny S (ed)., Evolution of plant-pollinator relationships. Cambridge University Press; Cambridge, UK; xv+477+[6] pp.

Minckley, R.L. 1998. A cladistic analysis and classification of the subgenera and genera of the large carpenter bees, tribe Xylocopini (Hymenoptera: Apidae). Scientific Papers, Nat. Hist. Mus. Univ. Kansas. 9, 1–47.

Minh, B.Q., Nguyen, M.A.T. von Haeseler, A. 2013. Ultrafast approximation for phylogenetic bootstrap. Mol. Biol. Evol. 30, 1188–1195. https://doi.org/10.1093/molbev/mst024

Mirande, J.M. 2009. Weighted parsimony phylogeny of the family Characidae (Teleostei: Characiformes). Cladistics 25, 574–613.

Mitchell, T.B. 1924. New megachilid bees. J. Elisha Mitchell Sci. Soc. 40, 154–165.

Mitchell, T.B. 1934. A revision of the genus Megachile in the Nearctic Region. Part I. Classification and descriptions of new species (Hymenoptera: Megachilidae). Trans. Am. Entomol. Soc. 59, 295–360 (Pls. XX–XXI).

Mitchell, T.B. 1935a. A revision of the genus Megachile in the Nearctic Region. Part II. Morphology of the male sternites and genital armature and the taxonomy of the subgenera Litomegachile, Neomegachile and Cressoniella (Hymenoptera: Megachilidae). Trans. Am. Entomol. Soc. 61, 1–44 (Pl. I).

Mitchell, T.B. 1935b. A revision of the genus Megachile in the Nearctic region. Part III. Taxonomy of subgenera Anthemois and Delomegachile (Hymenoptera: Megachilidae). Trans. Am. Entomol. Soc. 61, 155–205 (Pls. VIII–IX).

Mitchell, T.B. 1936. A revision of the genus Megachile in the Nearctic region. Part IV. Taxonomy of subgenera Xanthosarus, Phaenosarus, Megachiloides and Derotropis (Hymenoptera: Megachilidae). Trans. Am. Entomol. Soc. 62, 117–165 (Pls. VIII–XI).

Mitchell, T.B. 1937a. A revision of the genus Megachile in the Nearctic region. Part VI. Taxonomy of subgenus Xeromegachile (Hymenoptera: Megachilidae). Trans. Am. Entomol. Soc. 62, 323–382 (Pls. XXII–XXVI).

Mitchell, T.B. 1937b. A revision of the genus Megachile in the Nearctic region. Part VI. Taxonomy of subgenera Argyropile, Leptorachis, Pseudocentron, Acentron and Melanosarus (Hymenoptera: Megachilidae). Trans. Am. Entomol. Soc. 63, 45–83 (Pls. V–VI).

Mitchell, T.B. 1937c. A revision of the genus Megachile in the Nearctic region. Part VII. Taxonomy of the subgenus Sayapis (Hymenoptera: Megachilidae). Trans. Am. Entomol. Soc. 63, 175–206 (Pls. XII, XIII).

Mitchell, T.B. 1937d. A revision of the genus Megachile in the Nearctic region. Part VIII. Taxonomy of the subgenus Chelostomoides, addenda and index (Hymenoptera: Megachilidae). Trans. Am. Entomol. Soc. 63, 381–425 (Pls. XXVI–XXIX).

Mitchell, T.B. 1943. On the classification of neotropical Megachile (Hymenoptera, Megachilidae). Ann. Entomol. Soc. Am. 36, 656–671.

Mitchell, T.B. 1980. A generic revision of the megachiline bees of the Western Hemisphere. Raleigh, North Carolina: North Carolina State University. [ii] + 95 pp.

Moure, J.S., Melo, G.A.R., DalMolin, A. 2007. Megachilini Latreille, 1802, pp. 917–1001, in

Moure, J. S., D. Urban, and Melo, G. A. R (eds.), Catalogue of Bees (Hymenoptera, Apoidea) in the Neotropical Region. Curitiba, Brazil: Sociedade Brasileira de Entomologia.

Nguyen, L.-T., Schmidt, H.A., von Haeseler, A. Minh, B.Q. 2015. IQ-TREE: a fast and effective stochastic algorithm for estimating maximum likelihood phylogenies. Mol. Biol. Evol. 32, 268–274. https://doi.org/10.1093/molbev/msu300

Nixon, K.C. 1999. WINCLADA, version 0.9.99tuc.13, beta. Cornell University, Ithaca, New York.

O’Reilly, J.E., Donoghue, P.C.J. 2016. Tips and nodes are complementary not competing approaches to the calibration of molecular clocks. Biol. Lett. 12, 637–650.

O’Reilly, J.E., Reis, dos M., Donoghue, P.C.J. 2015. Dating tips for divergence-time estimation. Trends Genet. 31, 637–650.

Ornosa, C., Ortiz-Sánchez, F., Torres, F. 2007. Catálogo de los Megachilidae del Mediterráneo occidental (Hymenoptera, Apoidea). II. Lithurgini y Megachilini. Graellsia 63, 111–134.

Orr, M.C., Portman, Z.M., Griswold, T. 2015. Megachile (Megachile) montivaga (Hymenoptera: Megachilidae) nesting in live thistle (Asteraceae: Cirsium). J. Melittol. 48, 1–6.

Packer, L. 2003. Comparative morphology of the skeletal parts of the sting apparatus of bees (Hymenoptera: Apoidea). Zool. J. Linn. Soc. 138, 1–38.

Packer, L. 2004. Morphological variation in the gastral sterna of female Apoidea (Insecta: Hymenoptera). Can. J. Zool. 82, 130–152.

Pasteels, J.J. 1965. Revision des Megachilidae (Hymenoptera Apoidea) de L’Afrique noire. I. Les genres Creightoniella [sic], Chalicodoma et Megachile (s. str.). Musee Royal de L’Afrique Centrale. Tervuren, Belgique Annales. Serie IN-8, Sciences Zoologiques 137: 1–579.

Pasteels, J.J., Pasteels, J.M. 1971. Etude au microscope électronique à balayage des plages glandulaires tergales chez des espèces du genre Megachile (Hymenoptera, Apoïda, Megachilidae). Comptes Rendus Acad. Sci., Paris, Série D 273, 1481–1483.

Pauly, A., Brooks, R.W., Nilsson, A., Pesenko, Y.A., Eardley, C.D., Terzo, M., Griswold, T., Schwarz, M., Patiny, S., Munzinger, J., Barbier, Y. 2001. Hymenoptera Apoidea de Madagascar et des îles voisines. Mus. Roy. Afr. Centr. Ann. Sci. Zool. 286, 1–390, p.16

Pitts-Singer, T.L., Cane, J.H. 2011. The alfalfa leafcutting bee, Megachile rotundata: the world’s most intensively managed solitary bee. Annu. Rev. Entomol. 56, 221–237.

Potts, S.G., Imperatriz-Fonseca, V., Ngo, H.T., Aizen, M.A., Biesmeijer, J.C., Breeze, T.D., Dicks, L.V., Garibaldi, L.A., Hill, R., Settele, J., Vanbergen, A.J. 2016. Safeguarding pollinators and their values to human well-being. Nature doi:10.1038/nature20588

Praz, C.J. 2017. Subgeneric classification and biology of leafcutter and dauber bees (genus Megachile Latreille) of the western Paleartic (Hymenoptera, Apoidea, Megachilidae). J. Hymenopt. Res. 55, 1–54.

Praz, C.J., Müller, A., Danforth, B.N., Griswold, T., Widmer, A., Dorn, S. 2008. Phylogeny and biogeography of bees of the tribe Osmiini (Hymenoptera: Megachilidae). Mol. Phylogenet. Evol. 49, 185–197.

Provancher, L. 1882. Faune Canadienne. Les Insectes Hyménoptères. Nat. Can. 13, 225–242.

Pyron, R.A. 2011. Divergence time estimation using fossils as terminal taxa and the origins of Lissamphibia. Syst. Biol. 60, 446–481.

Radoszkowsky, O.I.B. 1874[1873]. Supplément indispensable à l’article publié par M. Gerstaecker en 1869, sur quelques genres d’Hyménoptères. (Suite et Fin). Bull. Soc. Imp. Nat. Moscou 47[46], 131–151, 1 pl.

Rambaut, A., Suchard, M., Xie, D., Drummon, A.J. 2014. Tracer v1.6. http://tree.bio.ed.ac.uk/software/tracer/ [accessed on Mar. 1, 2018]

Rasmussen, C., Carríon, A.L., Castro-Urgal, R., Chamorro, S., Gonzalez, V.H., Griswold, T.L., Herrera, H.W., McMullen, C.K., Olesen, J.M., Traveset, A. 2012. Megachile timberlakei Cockerell (Hymenoptera: Megachilidae): Yet another adventive bee species to the Galápagos Archipelago. Pan-Pac. Entomol. 88, 98–102.

Raw, A. 2006. A new subgenus and three new species of leafcutter bees, Megachile (Austrosarus) (Hymenoptera, Megachilidae) from central Brazil. Zootaxa 1228, 25–34.

Reemer, M., Ståhls, G. 2012. Unravelling a hotchpotch: phylogeny and classification of the microdontinae (Diptera: Syrphidae). Doctoral Dissertation, Leiden University, Leiden.

Reemer, M., Ståhls, G. 2013. Phylogenetic relationships of Microdontinae (Diptera: Syrphidae) based on molecular and morphological characters. Syst. Entomol. 38, 661–688.

Rieppel, O., Kearney, M. 2002. Similarity. Biol. J. Linn. Soc. 75, 59–82.

Robertson, C. 1901. Some new or little-known bees. Can. Entomol. 33, 229–231.

Robertson, C. 1903. Synopsis of Megachilidae and Bombinae. Trans. Am. Entomol. Soc. 29, 163–178.

Rocha Filho, L.C.D., Packer, L. 2017. Phylogeny of the cleptoparasitic Megachilini genera Coelioxys and Radoszkowskiana, with the description of six new subgenera in Coelioxys (Hymenoptera: Megachilidae). Zool. J. Linn. Soc. 180, 354–413.

Roig-Alsina, A., Michener, C.D. 1993. Studies of the phylogeny and classification of long-tongued bees (Hymenoptera: Apoidea). Univ. Kans. Sci. Bull. 55, 124–162.

Ronquist, F., Klopfstein, S., Vilhelmsen, L., Schulmeister, S., Murray, D.L. Rasnitsyn, A.P. 2012a. A total-evidence approach to dating with fossils, applied to the early radiation of Hymenoptera. Syst. Biol. 61, 973–999.

Ronquist, F., Teslenko, M., van der Mark, P., Ayres, D.L., Darling, A., Höhna, S., Larget, B., Liu, L., Suchard, M.A., Huelsenbeck, J.P. 2012b. MrBayes version 3.2: efficient Bayesian phylogenetic inference and model choice, across a large model space. Syst. Biol. 61, 539–542.

Roubik, D.W. 1982. Obligate necrophagy in a social bee. Science 217, 1059–1060.

Rozen, J.G., Ascher, J.S., Kamel, S.M., Mohamed, K.M. 2016. Larval diversity in the bee genus Megachile (Hymenoptera: Apoidea: Megachilidae). American Museum Novitates 3863, 1–16.

Rozen, J.G., Kamel, S.M. 2007. Investigations on the biologies and immature stages of the cleptoparasitic bee genera Radoszkowskiana and Coelioxys and their Megachile host (Hymenoptera: Apoidea: Megachilidae: Megachilini). Am. Mus. Novit. 3573, 1–43.

Rozen, J.G., Özbek, H., Ascher, J.S., Sedivy, C., Praz, C., Monfared, A., Müller, A. 2010. Nest, petal usage, floral preferences, and immatures of Osmia (Ozbekosmia) avosetta (Megachilidae: Megachilinae: Osmiini), including biological comparisons with other osmiine bees. Am. Mus. Novit. 3680, 1–22.

Sarzetti, L.C., Labandeira, C.C., Genise, J.F. 2008. A leafcutter bee trace fossil from the middle Eocene of Patagonia, Argentina, and a review of megachilid (Hymenoptera) ichnology. Palaeontology 51, 933–941.

Schofield, R.M.S., Nesson, M.H., Richardson, K.A. 2002. Tooth hardness increases with zinc-content in mandibles of young adult leaf-cutter ants. Naturwissenschaften 89, 579–583.

Schultz, T.R., Brady, S.G. 2008. Major evolutionary transitions in ant agriculture. PNAS 105, 5435–5440.

Schwarz, H.F. 1948. Stingless bees (Meliponidae) of the Western Hemisphere. Bull. Am. Mus. Nat. Hist. 90, 1–546.

Sereno, P.C. 2007. Logical basis for morphological characters in phylogenetics. Cladistics 23, 565–587.

Sheffield, C.S., Westby, S.M. 2007. The male of Megachile nivalis Friese, with an updated key to members of the subgenus Megachile s. str. (Hymenoptera: Megachilidae) in North America. J. Hymenopt. Res. 16, 178–191.

Silveira, F.A., Melo, G.A., Almeida, E.A. 2002. Abelhas Brasileiras. Sistemática e Identificação. Belo Horizonte, Brazil: Fernando A. Silveira. 253 pp.

Sinu, P.A., Bronstein, J.L. 2018. Foraging preferences of leafcutter bees in three contrasting geographical zones. Divers. Distrib. https://doi.org/10.1111/ddi.12709

Slater, G.J., Harmon, L.J., Freckleton, R. 2013. Unifying fossils and phylogenies for comparative analyses of diversification and trait evolution. Methods Ecol. Evol. 4, 699–702.

Smith, F. 1865. Descriptions of some new species of hymenopterous insects belonging to the families Thynnidae, Masaridae and Apidae. Trans. Entomol. Soc. Lond. 3, 389–399, pl. 21.

Talavera, G. Castresana, J. 2007. Improvement of phylogenies after removing divergent and ambiguously aligned blocks from protein sequence alignments. Syst. Biol. 56, 564–577.

Taschenberg, E. 1883. Die gattungen der bienen (Anthophila). Berliner Entomol. Z. 27, 37–100.

Thomson, C.G. 1872. Skandinaviens Hymenoptera. 2: a Delen, Innehållande Slägtet Apis Lin. Lund, Sweden.: Berlingska Boktryckeriet. 286 pp.

Torretta, J.P., Durante, S.P., Basilio, A.M. 2014. Nesting ecology of Megachile (Chrysosarus) catamarcensis Schrottky (Hymenoptera: Megachilidae), a Prosopis-specialist bee. J. Apic. Res. 53, 590–598.

Trunz, V., Packer, L., Vieu, J., Arrigo, N., Praz, C.J. 2016. Comprehensive phylogeny, biogeography and new classification of the diverse bee tribe Megachilini: can we use DNA barcodes in phylogenies of large genera? Mol. Phylogenet. Evol. 103, 245–259.

Wedmann S, Wappler T, Engel MS. 2009. Direct and indirect fossil records of megachilid bees from the Paleogene of Central Europe (Hymenoptera: Megachilidae). Naturwissenschaften 96, 703–712.

West-Eberhard, M.J. 2003. Developmental Plasticity and Evolution. Oxford University Press, New York.

Whiting, M.F., Bradler, S., Maxwell, T. 2003. Loss and recovery of wings in stick insects. Nature 421, 264–267.

Whitlock, J.A., Wilson, J.A. 2013. Character distribution maps: a visualization method for comparative cladistics. Acta Zool. 94, 490–499.

Williams, H.J., Strand, M.R, Elzen, G.W., Vinson, S.B., Merritt, S.J. 1986. Nesting behavior, nest architecture, and use of Dufour’s gland lipids in nest provisioning by Megachile integra and M. mendica mendica (Hymenoptera: Megachilidae). J. Kansas Entomol. Soc. 59, 588–597.

Williams, N.M., Goodell, K. 2000. Association of mandible shape and nesting material in Osmia Panzer (Hymenoptera: Megachilidae): a morphometric analysis. Ann. Entomol. Soc. Am. 93, 318–325.

Winston, M.L. 1979. The proboscis of the long-tongued bees: A comparative study. Univ. Kans. Sci. Bull. 51, 631–667.

Wittmann, D., Blochtein, B. 1995. Why males of leafcutter bees hold the females’ antennae with their front legs during mating. Apidologie 26, 181–195.

Zhang, C., Stadler, T., Klopfstein, S., Heath, T.A. Ronquist, F. 2016. Total-evidence dating under the fossilized birth-death process. Syst. Biol. 65, 228–249. https://doi.org/10.1093/sysbio/syv080

Zillikens, A., Steiner, J. 2004. Nest architecture, life cycle and cleptoparasite of the Neotropical leaf-cutting bee Megachile (Chrysosarus) pseudanthidioides Moure (Hymenoptera: Megachilidae). J. Kansas Entomol. Soc. 77, 193–202.

