## Appendix S1 for "Evolution of leaf-cutter behavior in bees (Hymenoptera: Megachilidae) as inferred from total-evidence tip-dating analyses"

Running title: Systematics and evolution of leaf-cutter bees

**Appendix S1.** Summary of recent major proposals in the classification of the genus *Megachile s.l.*, as discussed in the text. Original names of subgenera are under each genus. Shaded cells include the leaf-cutter taxa. * = A few recently described taxa are included within this classificatory scheme. † = Extinct taxa. § = *Cesacongoa* is a replacement name for *Cuspidella*.

| MICHENER (1965); PASTEELS (1965) | MITCHELL (1980) | MICHENER (2000, 2007)* |
|---|---|---|
| **Genus *Chalicodoma*:**<br>*Archimegachile*, *Austrochile*, *Callomegachile*, *Carinella*, *Cestella*, *Chalicodoma*, *Chalicodomoides*, *Chelostomoda*, *Chelostomoides*, *Cuspidella*, *Digronoceras*, *Dinavis*, *Eumegachilana*, *Gronoceras*, *Hackeriapis*, *Largella*, *Maximegachile*, *Morphella*, *Neglectella*, *Pseudomegachile*, *Rhodomegachile*, *Schizomegachile*, *Stelodides*, *Stenomegachile*, *Thaumatosoma* | **Genus *Chalicodoma*:**<br>*Archimegachile*, *Austrochile*, *Callomegachile*, *Carinella*, *Cestella*, *Chalicodoma*, *Chalicodomoides*, *Chelostomoda*, *Chelostomoides*, *Cuspidella*, *Digronoceras*, *Dinavis*, *Eumegachilana*, *Gronoceras*, *Hackeriapis*, *Largella*, *Maximegachile*, *Morphella*, *Neglectella*, *Pseudomegachile*, *Rhodomegachile*, *Schizomegachile*, *Stelodides*, *Stenomegachile*, *Thaumatosoma* | **Genus *Megachile***<br>Group 2:<br>*Alecanthedon*, *Austrochile*, *Callomegachile*, §*Cesacongoa*, *Cestella*, *Chalicodoma*, *Chalicodomoides*, †*Chalicodomopsis*, *Chelostomoda*, *Chelostomoides*, *Gronoceras*, *Hackeriapis*, *Heriadopsis*, *Largella*, *Lophanthedon*, *Matangapis*, *Maximegachile*, *Megella*, *Mitchellapis*, *Neochalicodoma*, *Parachalicodoma*, *Pseudomegachile*, *Rhodomegachile*, *Schizomegachile*, *Stellenigris*, *Stenomegachile*, *Thaumatosoma* |
| **Genus *Creightonella***<br>**Genus *Megachile*:**<br>*Acentron*, *Amegachile*, *Argyropile*, *Austromegachile*, *Callochile*, *Chrysosarus*, *Cressoniella*, *Dactylomegachile*, *Dasymegachile*, *Delomegachile*, *Derotropis*, *Digitella*, *Eumegachile*, *Eurymella*, *Eutricharaea* *Holcomegachile*, *Leptorachis*, *Litomegachile*, *Megachile*, *Megachiloides*, *Megella*, *Melanosarus*, *Mitchellapis*, *Neomegachile*, *Paracella*, *Phaenosarus*, *Platysta*, *Pseudocentron*, *Ptilosarus*, *Sayapis*, *Tylomegachile*, *Xanthosarus*, *Xeromegachile* | **Genus *Creightonella***<br>**Genus *Chrysosarus***<br>*Chrysosarus*, *Dactylomegachile*, *Stelodides*, *Zonomegachile* | Group 3: *Creightonella*<br>Group 1:<br>*Acentron*, *Aethomegachile*, *Amegachile*, *Aporiochile*, *Argyropile*, *Austrosarus*, *Austromegachile*, *Chalepochile*, *Chrysosarus*, *Cressoniella*, *Dasymegachile*, *Eumegachile*, *Eutricharaea*, *Grosapis*, *Leptorachis*, *Litomegachile*, *Megachile*, *Megachiloides*, *Melanosarus*, *Moureapis*, *Neochelynia*, *Neocressoniella*, *Paracella*, *Platysta*, *Pseudocentron*, *Ptilosaroides*, *Ptilosarus*, *Rhyssomegachile*, *Sayapis*, *Schrottkyapis*, *Stelodides*, *Trichurochile*, *Tylomegachile*, *Xanthosarus*, *Zonomegachile* |
| | **Genus *Cressoniella***<br>*Austromegachile*, *Chaetochile*, *Cressoniella*, *Dasymegachile*, *Holcomegachile*, *Neomegachile*, *Ptilosaroides*, *Ptilosarus*, | |

*Rhyssomegachile*, *Trichurochile*, *Tylomegachile*
**Genus *Eumegachile*:**
*Eumegachile*, *Grosapis*, *Mitchellapis*, *Sayapis*, *Schrottkyapis*

**Genus *Megachile*:** *Addendella*, *Amegachile*, *Callochile*, *Delomegachile*, *Digitella*, *Eurymella*, *Eutricharaea Litomegachile*, *Macromegachile*, *Megachile*, *Megella*, *Paracella*, *Platysta*, *Xanthosarus*
**Genus *Megachiloides*:** *Argyropile*, *Derotropis*, *Megachiloides*, *Phaenosarus*, *Xeromegachile*
**Genus *Pseudocentron*:** *Acentron*, *Grafella*, *Leptorachina*, *Leptorachis*, *Melanosarus*, *Moureana*, *Pseudocentron*
