## Appendix S2 for "Evolution of leaf-cutter behavior in bees (Hymenoptera: Megachilidae) as inferred from total-evidence tip-dating analyses"

Running title: Systematics and evolution of leaf-cutter bees

**Appendix S2.** List of taxa included in the morphological and molecular analyses of the family Megachilidae and tribe Megachilini. We followed the classifications of Gonzalez *et al*. (2012) for the tribes of Megachilidae and that of Michener (2007) for the subgenera of *Megachile* s.l., which also includes recently described taxa. In the combined analyses, we used closely related species to those used in the morphological analyses when molecular data was not available for the same species. We referred to those chimeric taxa by their genus name, and sometimes subgenus, followed by a combination of the first three letters of both specific epithets in square brackets. For example, the name for the operational taxonomic unit (OTU) resulting from *Trichothurgus wagenknechti* and *T. herbsti* is referred herein as *Trichothurgus* [wag×her] (see text for explanation). GenBank accession numbers for specimens of included species in the analyses are in Appendix S3. * = Type species of the subgenus of *Megachile* s.l., as recognized by Michener (2007). † = Extinct species. – = Species not included in the analysis.

| PHYLOGENY OF MEGACHILIDAE | | |
|---|---|---|
| Morphological analysis | Molecular analysis | Combined analysis |
| **Outgroups** | | |
| MELITTIDAE | | |
| *Macropis* (*Macropis*) *nuda* (Provancher, 1882) [USA] | *Macropis nuda* | *Macropis nuda* |
| *Melitta* (*Melitta*) *leporina* (Panzer, 1799) [France, Spain, Iran] | *Melitta leporina* | *Melitta leporina* |
| APIDAE | | |
| *Apis mellifera* Linnaeus, 1758 [USA] | *Apis mellifera* | *Apis mellifera* |
| *Exomalopsis* (*Stilbomalopsis*) *solani* Cockerell, 1896 [USA, Mexico] | *Exomalopsis* sp. | *Exomalopsis* [sol×sp] |
| *Diadasia* (*Coquillettapis*) *australis* (Cresson, 1878) [USA] | *Diadasia bituberculata* (Cresson, 1878) | *Diadasia* [aus×bit] |
| *Nomada utahensis* Moalif, 1988 [USA] | *Nomada maculata* Cresson, 1863 | *Nomada* [uta×mac] |
| *Ceratina calcarata* Robertson, 1900 [USA] | *Ceratina calcarata* | *Ceratina calcarata* |
| **Ingroup** | | |
| MEGACHILIDAE | | |
| FIDELIINAE | | |
| *Fidelia* (*Parafidelia*) *pallidula* (Cockerell, 1935) [South Africa] | *Fidelia pallidula* | *Fidelia pallidula* |
| *F*. (*Fidelia*) *villosa* Brauns, 1902 [South Africa] | *F. villosa* | *F. villosa* |
| *F*. (*Fideliana*) *braunsiana* Friese, 1905 [South Africa] | *F. braunsiana* | *F. braunsiana* |
| *F*. (*Fideliopsis*) *major* Friese, 1911 [South Africa] | *F. major* | *F. major* |
| *Neofidelia profuga* Moure & Michener, 1955 [Chile] | *Neofidelia profuga* | *Neofidelia profuga* |
| PARARHOPHITINAE | | |
| *Pararhophites orobinus* (Morawitz, 1875) [Pakistan] | *Pararhophites orobinus* | *Pararhophites orobinus* |
| *P. quadratus* (Friese, 1898) [Egypt] | *P. quadratus* | *P. quadratus* |
| LITHURGINAE | | |

| | | |
|---|---|---|
| LITHURGINI | | |
| *Lithurgus* (*Lithurgopsis*) *apicalis* Cresson, 1875 [USA] | *Lithurgus chrysosarus* Fonscolombe, 1834 | *Lithurgus* [api×chr] |
| *Microthurge corumbae* (Cockerell, 1901) [Bolivia] | *Microthurge* sp. | *Microthurge* [cor×sp] |
| *Trichothurgus aterrimus* (Cockerell, 1914) [Chile] | *Trichothurgus herbstei* (Friese, 1905) | *Trichothurgus* [ate×her] |
| †PROTOLITHURGINI | | |
| †*Protolithurgus ditomeus* Engel, 2001 | | |
| MEGACHILINAE | | |
| ANTHIDIINI | | |
| *Afranthidium* (*Capanthidium*) *capicola* (Brauns, 1905) [South Africa] | *Afranthidium capicola* | *Afranthidium capicola* |
| *Anthidiellum* (*Loyolanthidium*) *robertsoni* (Cockerell, 1904) [USA] | *Anthidiellum robertsoni* | *Anthidiellum robertsoni* |
| *Anthidium* (*Anthidium*) *porterae* Cockerell, 1900 [USA] | *Anthidium porterae* | *Anthidium porterae* |
| *Anthodioctes* (*Anthodioctes*) *calcaratus* (Friese, 1921) [Costa Rica] | *Anthodioctes* (*Anthodioctes*) *mapirensis* (Cockerell, 1927) | *Anthodioctes* [cal×map] |
| *Aztecanthidium tenochtitlanicum* Snelling, 1987 [Mexico] | *Aztecanthidium tenochtitlanicum* | *Aztecanthidium tenochtitlanicum* |
| *Cyphanthidium intermedium* Pasteels, 1969 [Namibia] | *Cyphanthidium intermedium* | *Cyphanthidium intermedium* |
| *Dianthidium* (*Dianthidium*) *subparvum* Swenk, 1914 [USA] | *Dianthidium subparvum* | *Dianthidium subparvum* |
| *Duckeanthidium thielei* Michener [Costa Rica: Heredia] | *Duckeanthidium thielei* | *Duckeanthidium thielei* |
| *Eoanthidium* (*Clistanthidium*) *rothschildi* (Vachal, 1909) [South Africa] | *Eoanthidium* (*Clistanthidium*) *turnericum* (Mavromoustakis, 1934) | *Eoanthidium* [rot×tur] |
| *Epanthidium* (*Epanthidium*) *bicoloratum* (Smith, 1879) [Argentina] | *Epanthidium bicoloratum* | *Epanthidium bicoloratum* |
| *Euaspis abdominalis* (Fabricius, 1773) [Zambia] | *Euaspis abdominalis* | *Euaspis abdominalis* |
| *Hoplostelis* (*Hoplostelis*) *bivittata* (Cresson, 1878) [Costa Rica] | *Hoplostelis bivittata* | *Hoplostelis bivittata* |
| *Hypanthidioides* (*Michanthidium*) *ferrugineum* (Urban, 1992[1994]) [Argentina] | *Hypanthidioides* (*Saranthidium*) *marginata* (Moure & Urban, 1994) | *Hypanthidioides* [fer×mar] |
| *Hypanthidium* (*Hypanthidium*) *mexicanum* (Cresson, 1878) [Mexico] | *Hypanthidium* (*Hypanthidium*) *obscurius* Schrottky, 1908 | *Hypanthidium* [mex×obs] |
| *Icteranthidium ferrugineum* (Fabricius, 1787) [Egypt, Tunisia] | *Icteranthidium ferrugineum* | *Icteranthidium ferrugineum* |
| *Notanthidium* (*Notanthidium*) *steloides* (Spinola, 1851) [Chile] | *Notanthidium steloides* | *Notanthidium steloides* |
| *Pachyanthidium* (*Pachyanthidium*) *katangense* Cockerell, 1930 [Congo] | *Pachyanthidium* (*Pachyanthidium*) *cordatum* (Smith, 1854) | *Pachyanthidium* [kat×cor] |

| | | |
|---|---|---|
| *Plesianthidium* (*Spinanthidiellum*) *rufocaudatum* (Friese, 1909) [South Africa] | *Plesianthidium rufocaudatum* | *Plesianthidium rufocaudatum* |
| *Pseudoanthidium* (*Micranthidium*) *lanificum* (Smith, 1879) [Cameroon, Congo] | *Pseudoanthidium* sp. | *Pseudoanthidium* [lan×sp] |
| *Rhodanthidium* (*Rhodanthidium*) *septemdentatum* (Latreille, 1809) [Greece] | *Rhodanthidium septemdentatum* | *Rhodanthidium septemdentatum* |
| *Serapista rufipes* (Friese, 1904) [South Africa] | *Serapista rufipes* | *Serapista rufipes* |
| *Stelis* (*Stelis*) *linsleyi* Timberlake, 1941 [USA] | *Stelis* (*Stelis*) *lateralis* Cresson, 1864 | *Stelis* [lin×lat] |
| *Trachusa* (*Heteranthidium*) *larreae* (Cockerell, 1897) [USA] | *Trachusa larreae* | *Trachusa larreae* |
| ASPIDOSMIINI | | |
| *Aspidosmia arnoldi* (Brauns, 1926) [South Africa] | *Aspidosmia arnoldi* | *Aspidosmia arnoldi* |
| *Aspidosmia volkmanni* (Friese, 1909) [South Africa] | *A. volkmanni* | *A. volkmanni* |
| †CTENOPLECTRELLINI | | |
| †*Ctenoplectrella cockerelli* Engel, 2001 | | |
| †*C. grimaldii* Engel, 2001 | | |
| †*C. viridiceps* Cockerell, 1909 | | |
| †*Glaesosmia genalis* Engel, 2001 | | |
| DIOXYNI | | |
| *Aglaoapis tridentata* (Nylander, 1848) [Austria] | *Aglaoapis tridentata* | *Aglaoapis tridentata* |
| *Dioxys pomonae* Cockerell, 19010 [USA] | *Dioxys moesta* Costa, 1883 | *Dioxys* [pom×moe] |
| †GLYPTAPINI | | |
| †*Glyptapis densopunctata* Engel, 2001 | | |
| †*G. disareolata* Engel, 2001 | | |
| MEGACHILINI | | |
| *Coelioxys* (*Boreocoelioxys*) *octodentata* Say, 1824 [USA] | *Coelioxys octodentata* | *Coelioxys octodentata* |
| *Megachile* (*Chelostomoides*) *angelarum* Cockerell, 1902 [USA] | *Megachile* (*Chelostomoides*) *angelarum* | *Megachile* (*Chelostomoides*) *angelarum* |
| *M.* (*Creightonella*) *discolor* Smith, 1853 [South Africa] | *M.* (*Creightonella*) *albisecta* Klug, 1817 | *M.* (*Creightonella*) [dis×alb] |
| *M.* (*Sayapis*) *pugnata* Say, 1873 [USA] | *M.* (*Sayapis*) *pugnata* | *M.* (*Sayapis*) *pugnata* |
| *Noteriades spinosus* Griswold & Gonzalez, 2011 [Thailand] | *Noteriades* sp. | *Noteriades* [spi×sp] |
| *Radoszkowskiana rufiventris* (Spinola, 1838) [Egypt] | *Radoszkowskiana rufiventris* | *Radoszkowskiana rufiventris* |
| OSMIINI | | |
| *Afroheriades hyalinus* Griswold & Gonzalez, 2011 [South Africa] | *Afroheriades primus* (Peters, 1970) | *Afroheriades* [hya×pri] |

| | | |
|---|---|---|
| *Ashmeadiella* (*Ashmeadiella*) *aridula* Cockerell, 1910 [USA] | *Ashmeadiella aridula* | *Ashmeadiella aridula* |
| *Atoposmia* (*Atoposmia*) *abjecta* (Cresson, 1878) [USA] | *Atoposmia* (*Eremosmia*) *mirifica* (Michener, 1954) | *Atoposmia* [abj×mir] |
| *Chelostoma* (*Chelostoma*) *florisomne* (Linnaeus, 1758) [Hungry, Sweden] | *Chelostoma florisomne* | *Chelostoma florisomne* |
| *Haetosmia vechti* (Peters, 1974) [Israel, Pakistan] | *Haetosmia brachyura* (Morawitz, 1875) | *Haetosmia* [vec×bra] |
| *Heriades* (*Heriades*) *truncorum* (Linnaeus, 1758) [Austria, Sweden] | *Heriades crucifer* Cockerell, 1897 | *Heriades* [tru×cru] |
| *Hofferia schmiedeknechti* (Schletterer, 1889) [Bulgaria, Greece] | *Hofferia schmiedeknechti* | *Hofferia schmiedeknechti* |
| *Hoplitis* (*Monumetha*) *albifrons* (Kirby, 1873) [USA: Utah] | *Hoplitis* (*Hoplitis*) *adunca* (Panzer, 1798) | *Hoplitis* [alb×adu] |
| *H.* (*Stenosmia*) *flavicornis* (Morawitz, 1877) [Mongolia, Uzbekistan] | *H.* (*Stenosmia*) *minima* (Schulthess, 1924) | *Hoplitis* [fla×min] |
| *Ochreriades fasciatus* (Friese, 1899) [Israel] | *Ochreriades fasciatus* | *Ochreriades fasciatus* |
| *Osmia* (*Osmia*) *lignaria* Say, 1837 [USA] | *Osmia lignaria* | *Osmia lignaria* |
| *Othinosmia* (*Megaloheriades*) *globicola* (Stadelmann, 1892) [South Africa] | *Othinosmia globicola* | *Othinosmia globicola* |
| *Protosmia* (*Chelostomopsis*) *rubifloris* (Cockerell, 1898) [USA] | *Protosmia* (*Protosmia*) *humeralis* (Pérez, 1895) | *Protosmia* [rub×hum] |
| *Pseudoheriade moricei* (Friese, 1897) [Egypt] | *Pseudoheriade moricei* | *Pseudoheriade moricei* |
| *Stenoheriades asiaticus* (Friese, 1921) [Turkey] | *Stenoheriades asiaticus* | *Stenoheriades asiaticus* |
| *Wainia* (*Caposmia*) *elizabethae* (Friese, 1909) [South Africa] | *Wainia* (*Caposmia*) *eremoplana* (Mavromoustakis, 1949) | *Wainia* [eli×ere] |

PHYLOGENY OF MEGACHILINI

**Outgroups**

| | | |
|---|---|---|
| ANTHIDIINI | | |
| *Aztecanthidium tenochtitlanicum* Snelling, 1987 [Mexico] | *A. tenochtitlanicum* | *A. tenochtitlanicum* |
| *Trachusa mitchelli* (Michener, 1948) [Mexico] | *T. larreae* (Cockerell, 1897) | *Trachusa* [mit×lar] |
| ASPIDOSMIINI | | |
| *Aspidosmia volkmanni* (Friese, 1909) [South Africa] | *A. volkmanni* | *A. volkmanni* |
| DIOXYINI | | |
| *Dioxys producta* (Cresson, 1879) [USA] | *Dioxys moesta* Costa, 1883 | *Dioxys* [pro×moe] |
| LITHURGINI | | |
| *Microthurge friesei* (Ducke, 1907) [Argentina] | *Microthurge* sp. | *Microthurge* [fri×sp] |

| | | |
|---|---|---|
| *Trichothurgus wagenknechti* (Moure, 1949) [Chile] | *Trichothurgus herbsti* (Friese, 1905) | *Trichothurgus* [wag×her] |
| OSMIINI | | |
| *Chelostoma rapunculi* (Lepeletier, 1841) [USA] | *Chelostoma florisomne* (Linnaeus, 1758) | *Chelostoma* [rap×flo] |
| *Hoplitis biscutellae* (Cockerell, 1897) [USA] | *Hoplitis adunca* (Panzer, 1798) | *Hoplitis* [rap×flo] |
| **Ingroup** | | |
| MEGACHILINI | | |
| *Coelioxys* (*Rhinocoelioxys*) *zapoteca* Cresson, 1878 [Argentina, Bolivia, Brazil, Mexico] | — | — |
| *C.* (*Liothyrapis*) *decipiens* Spinola, 1838 [India] | *Coelioxys decipiens* | *Coelioxys decipiens* |
| *C.* (*Torridapis*) *torrida* Smith, 1854 [South Africa] | — | — |
| *Megachile* s. l. | | |
| GROUP 1 | | |
| *M. (*Acentron*) *albitarsis* Cresson, 1872 [USA] | *M.* (*Acentron*) sp. | *M.* (*Acentron*) [alb×sp] |
| *M.* (*Acentron*) *candida* Smith, 1879 [Costa Rica] | — | — |
| *M.* (*Aethomegachile*) *laticeps* Smith, 1853 [India] | *M.* (*Aethomegachile*) *conjuncta* Smith, 1853 | *M.* (*Aethomegachile*) [lat×con] |
| *M. (*Aethomegachile*) *trichorhytisma* Engel, 2006 [Thailand] | — | — |
| *M. (*Amegachile*) *bituberculata* Ritsema, 1880 [Cameroom] | *M.* (*Amegachile*) c.f. *bituberculata* | *M.* (*Amegachile*) *bituberculata* |
| *M.* (*Amegachile*) *ustulatiformis* Cockerell, 1910 [Australia] | — | — |
| *M. (*Argyropile*) *parallela* Smith, 1853 [USA] | *M.* (*Argyropile*) *parallela* | *M.* (*Argyropile*) *parallela* |
| *M.* (*Argyropile*) *sabinensis* Mitchell, 1934 [USA] | — | — |
| *M.* (*Austromegachile*) *exaltata* Smith, 1853 [Brazil] | — | — |
| *M. (*Austromegachile*) *montezuma* Cresson, 1878 [Brazil] | *M.* (*Austromegachile*) sp. | *M.* (*Austromegachile*) [mon×sp] |
| *M. (*Chrysosarus*) *guaranitica* Schrottky, 1908 [Paraguay] | *M.* (*Chrysosarus*) sp. | *M.* (*Chrysosarus*) [pse×sp] |
| *M.* (*Chrysosarus*) *parsonsiae* Schrottky, 1914 [Argentina] | — | — |
| *M.* (*Chrysosarus*) *pseudanthidioides* Moure, 1943 [Brazil] | — | — |
| *M. (*Cressoniella*) *zapoteca* Cresson, 1878 [Mexico] | *M.* (*Cressoniella*) *zapoteca* | *M.* (*Cressoniella*) *zapoteca* |
| *M.* (*Dasymegachile*) *mitchelli* Raw, 2004 [Argentina, Peru] | — | — |
| *M. (*Dasymegachile*) *saulcyi* Guérin, 1845 [Chile] | *M.* (*Dasymegachile*) sp. | *M.* (*Dasymegachile*) [sau×sp] |
| *M. (*Eumegachile*) *bombycina* Radoszkowski, 1874 [Finland] | *M.* (*Eumegachile*) *bombycina* | *M.* (*Eumegachile*) *bombycina* |
| *M. (*Eutricharaea*) *argentata* Fabricius, 1793 [USA] | — | — |
| *M.* (*Eutricharaea*) *digiticauda* Cockerell, 1937 [Zimbabwe] | — | — |
| *M.* (*Eutricharaea*) *eurymera* Smith, 1854 [Kenya, Nigeria] | *M.* (*Eutricharaea*) aff. *eurymera* | *M.* (*Eutricharaea*) *eurymera* |
| *M.* (*Eutricharaea*) *femorata* Smith, 1853 [India] | — | — |
| *M.* (*Eutricharaea*) *leachella* Curtis, 1828 [Slovakia] | — | — |

| | | |
|---|---|---|
| *M.* (*Eutricharaea*) *rotundata* Fabricius, 1787 [USA] | *M.* (*Eutricharaea*) *rotundata* | *M.* (*Eutricharaea*) *rotundata* |
| *M.* (*Eutricharaea*) *submetallica* Benoist, 1954 [Madagascar] | — | — |
| **M.* (*Grosapis*) *cockerelli* Rohwer, 1923 [Mexico] | *M.* (*Grosapis*) *cockerelli* | *M.* (*Grosapis*) *cockerelli* |
| *M.* (*Leptorachis*) *crotalariae* Schwimmer, 1980 [Brazil] | — | — |
| *M.* (*Leptorachis*) *laeta* Smith, 1853 [Brazil] | — | — |
| **M.* (*Leptorachis*) *petulans* Cresson, 1878 [USA] | *M.* (*Leptorachis*) *petulans* | *M.* (*Leptorachis*) *petulans* |
| **M.* (*Litomegachile*) *brevis* Say, 1837 [USA] | *M.* (*Litomegachile*) *texana* Cresson, 1878 | *M.* (*Litomegachile*) [bre×tex] |
| *M.* (*Litomegachile*) *gentilis* Cresson, 1872 [USA] | — | — |
| **M.* (*Megachile*) *centuncularis* Linnaeus, 1758 [USA] | — | — |
| *M.* (*Megachile*) *montivaga* Cresson, 1878 [USA] | — | — |
| *M.* (*Megachiloides*) *integra* Cresson, 1878 [USA] | *M.* (*Megachiloides*) *nevadensis* Cresson, 1879 | *M.* (*Megachiloides*) [int×nev] |
| **M.* (*Megachiloides*) *oenotherae* Mitchell, 1924 [USA] | — | — |
| *M.* (*Megachiloides*) *pascoensis* Mitchell, 1934 [USA] | — | — |
| *M.* (*Melanosarus*) *nigripennis* Spinola, 1841 [Brazil] | — | — |
| **M.* (*Melanosarus*) *xylocopoides* Smith, 1853 [USA] | *M.* (*Melanosarus*) sp. | *M.* (*Melanosarus*) [xyl×sp] |
| **M.* (*Moureapis*) *anthidioides* Radoszkowski, 1874 [Brazil] | — | — |
| *M.* (*Neochelynia*) *chichimeca* Cresson, 1878 [Mexico] | *M.* (*Neochelynia*?) sp. | *M.* (*Neochelynia*) [chi×sp] |
| **M.* (*Neochelynia*) *paulista* Schrottky, 1920 [Brazil] | — | — |
| **M.* (*Neocressoniella*) *carbonaria* Smith, 1853 [India] | — | — |
| *M.* (*Paracella*) *curtula* Gerstaecker, 1857 [Uganda] | — | — |
| **M.* (*Paracella*) *semivenusta* Cockerell, 1931 [Malawi] | *M.* (*Paracella*) sp. | *M.* (*Paracella*) [sem×sp] |
| **M.* (*Platysta*) *platystoma* Pasteels, 1965 [Congo] | — | — |
| *M.* (*Pseudocentron*) *poeyi* Guérin, 1845 [Cuba] | — | — |
| **M.* (*Pseudocentron*) *pruina* Smith, 1853 [USA] | *M.* (*Pseudocentron*) sp. | *M.* (*Pseudocentron*) [pru×sp] |
| **M.* (*Ptilosaroides*) *neoxanthoptera* Cockerell, 1933 [Panama] | — | — |
| *M.* (*Ptilosarus*) *microsoma* Cockerell, 1912 [Trinidad and Tobago] | *M.* (*Ptilosarus*) *microsoma* | *M.* (*Ptilosarus*) *microsoma* |
| **M.* (*Rhyssomegachile*) *simillima* Smith, 1853 [Brazil] | — | — |
| *M.* (*Sayapis*) *coelioxyformis* Schrottky, 1910 [Brazil, Paraguay] | — | — |
| **M.* (*Sayapis*) *pugnata* Say, 1837 [USA] | *M.* (*Sayapis*) *pugnata* | *M.* (*Sayapis*) *pugnata* |
| **M.* (*Schrottkyapis*) *assumptionis* Schrottky, 1908 [Brazil] | — | — |
| **M.* (*Steloides*) *euzona* Pérez, 1899 [Chile] | *M.* (*Steloides*) *euzona* | *M.* (*Steloides*) *euzona* |
| **M.* (*Trichurochile*) *thygaterella* Schrottky, 1913 [Peru, Brazil] | — | — |
| **M.* (*Tylomegachile*) *orba* Schrottky, 1913 [Mexico] | *M.* (*Tylomegachile*) sp. | *M.* (*Tylomegachile*) [orb×sp] |

| | | |
|---|---|---|
| *M.* (*Tylomegachile*) *simplicipes* Friese, 1921 [Mexico] | — | — |
| *M.* (*Xanthosarus*) *addenda* Cresson, 1878 [USA] | — | — |
| *M.* (*Xanthosarus*) *fortis* Cresson, 1872 [USA] | *M.* (*Xanthosarus*) *fortis* | *M.* (*Xanthosarus*) *fortis* |
| *M.* (*Xanthosarus*) *lagopoda* Linnaeus, 1761 [Spain] | *M.* (*Xanthosarus*) *lagopoda* | *M.* (*Xanthosarus*) *lagopoda* |
| **M.* (*Xanthosarus*) *latimanus* Say, 1823 [USA] | — | — |
| **M.* (*Zonomegachile*) *moderata* Smith, 1879 [Brazil] | — | — |
| GROUP 2 | | |
| *M.* (*Alocanthedon*) *memecylonae* Engel, 2011 [Malaysia] | *M.* (*Alocanthedon*) sp. | *M.* (*Alocanthedon*) [orb×sp] |
| **M.* (*Austrochile*) *resinifera* Meade-Waldo, 1915 [Australia] | *M.* (*Austrochile*) sp. | *M.* (*Austrochile*) [res×sp] |
| *M.* (*Callomegachile*) *biseta* Vachal, 1903 [Gabon] | — | — |
| *M.* (*Callomegachile*) *clotho* Smith, 1861 [NE. Sulawesi] | — | — |
| *M.* (*Callomegachile*) *decemsignata* Radoszkowski, 1881 [Uganda] | *M.* (*Callomegachile*) *decemsignata* | *M.* (*Callomegachile*) *decemsignata* |
| **M.* (*Callomegachile*) *mystaceana* Michener, 1962 [Australia] | — | — |
| *M.* (*Callomegachile*) *sculpturalis* Smith, 1853 [Japan, USA] | *M.* (*Callomegachile*) *sculpturalis* | *M.* (*Callomegachile*) *sculpturalis* |
| *M.* (*Callomegachile*) *torrida* Smith, 1853 [Uganda] | — | — |
| **M.* (*Cesacongoa*) *quadraticauda* Pasteels, 1965 [Congo] | *M.* (*Cesacongoa*) sp. | *M.* (*Cesacongoa*) [qua×sp] |
| **M.* (*Cestella*) *cestifera* Benoist, 1954 [Madagascar] | — | — |
| *M.* (*Chalicodoma*) *asiatica* Morawitz, 1875 [Turkey] | — | — |
| *M.* (*Chalicodoma*) *lefebvrei* Lepeletier, 1841 [Greece, Italy] | *M.* (*Chalicodoma*) *lefebvrei* | *M.* (*Chalicodoma*) *lefebvrei* |
| *M.* (*Chalicodoma*) *manicata* Giraud, 1861 [Kazakhstan] | *M.* (*Chalicodoma*) *manicata* | *M.* (*Chalicodoma*) *manicata* |
| **M.* (*Chalicodoma*) *parietina* Geoffroy, 1785 [Spain] | *M.* (*Chalicodoma*) *parietina* | *M.* (*Chalicodoma*) *parietina* |
| **M.* (*Chalicodomoides*) *aethiops* Smith, 1853 [Australia] | *M.* (*Chalicodomoides*) *aethiops* | *M.* (*Chalicodomoides*) *aethiops* |
| †**M.* (*Chalicodomopsis*) *glaesaria* Engel, 1999 [Dominican Republic] | | |
| **M.* (*Chelostomoda*) *spissula* Cockerell, 1911 [China] | *M.* (*Chelostomoda*) sp. | *M.* (*Chelostomoda*) [spi×sp] |
| *M.* (*Chelostomoda*) *ulrica* Nurse, 1901 [India] | — | — |
| | *M.* (*Chelostomoides*) *angelarum* Cockerell, 1902 | *M.* (*Chelostomoides*) [cam×ang] |
| *M.* (*Chelostomoides*) *campanulae* Robertson, 1903 [USA] | | |
| *M.* (*Chelostomoides*) *georgica* Cresson, 1878 [USA] | — | — |
| **M.* (*Chelostomoides*) *rugifrons* Smith, 1854 [USA] | — | — |
| *M.* (*Chelostomoides*) *spinotulata* Mitchell, 1934 [USA] | *M.* (*Chelostomoides*) *spinotulata* | *M.* (*Chelostomoides*) *spinotulata* |
| **M.* (*Gronoceras*) *bombiformis* Gerstaecker, 1857 [Tanzania] | *M.* (*Gronoceras*) *bombiformis* | *M.* (*Gronoceras*) *bombiformis* |
| *M.* (*Gronoceras*) *cincta combusta* (Smith, 1853) [Tanzania] | — | — |
| *M.* (*Hackeriapis*) *ferox* Smith, 1879 [Australia] | — | — |
| *M.* (*Hackeriapis*) *heriadiformis* Smith, 1853 [Australia] | — | — |

| | | |
|---|---|---|
| *M.* (*Hackeriapis*) *ignita* Smith, 1853 [Australia] | — | — |
| **M.* (*Hackeriapis*) *rhodura* Cockerell, 1906 [Australia] | *M.* (*Hackeriapis*) sp. 1 | *M.* (*Hackeriapis*) [rho×sp] |
| **M.* (*Heriadopsis*) *striatulus* Cockerell, 1931 [Zimbabwe] | *M.* (*Heriadopsis*) sp. | *M.* (*Heriadopsis*) [str×sp] |
| **M.* (*Largella*) *semivestita* Smith, 1853 [C. Java] | *M.* (*Largella*) *floralis* (Fabricius, 1804) | *M.* (*Largella*) [sem×flo] |
| *M.* (*Lophanthedon*) *dimidiata* Smith, 1853 [Malaysia] | *M.* (*Lophanthedon*) *dimidiata* | *M.* (*Lophanthedon*) *dimidiata* |
| **M.* (*Matangapis*) *alticola* Cameron, 1902 [Borneo] | *M.* (*Matangapis*) *alticola* | *M.* (*Matangapis*) *alticola* |
| **M.* (*Maximegachile*) *maxillosa* Guérin, 1845 [Kenya, Natal, Tanzania] | *M.* (*Maximegachile*) *maxillosa* | *M.* (*Maximegachile*) *maxillosa* |
| **M.* (*Megella*) *malimbana* Strand, 1911 [Zaire] | — | — |
| *M.* (*Megella*) *pseudomonticola* Hedicke, 1925 [Japan] | *M.* (*Megella*) *pseudomonticola* | *M.* (*Megella*) *pseudomonticola* |
| **M.* (*Mitchellapis*) *fabricator* Smith, 1868 [Australia] | *M.* (*Mitchellapis*) *fabricator* | *M.* (*Mitchellapis*) *fabricator* |
| **M.* (*Parachalicodoma*) *incana* Friese, 1898 [Egypt] | *M.* (*Parachalicodoma*) sp. | *M.* (*Parachalicodoma*) [inc×sp] |
| *M.* (*Pseudomegachile*) *albocincta* Radoszkowski, 1874 [Egypt] | — | — |
| *M.* (*Pseudomegachile*) *armatipes* Friese, 1909 [Natal] | *M.* (*Pseudomegachile*) *laminata* Friese, 1903 | *M.* (*Pseudomegachile*) [arm×lam] |
| **M.* (*Pseudomegachile*) *ericetorum* Lepeletier, 1841 [Spain] | *M.* (*Pseudomegachile*) *ericetorum* | *M.* (*Pseudomegachile*) *ericetorum* |
| *M.* (*Pseudomegachile*) *flavipes* Spinola, 1838 [India] | — | — |
| *M.* (*Pseudomegachile*) *muansae* Friese, 1911 [Tanzania] | *M.* (*Pseudomegachile*) *leucospilura* Cockerell, 1937 | *M.* (*Pseudomegachile*) [mua×leu] |
| **M.* (*Rhodomegachile*) *abdominalis* Smith, 1853 [Australia] | *M.* (*Rhodomegachile*) sp. | *M.* (*Rhodomegachile*) [abd×sp] |
| **M.* (*Schizomegachile*) *monstrosa* Smith, 1868 [Australia] | — | — |
| **M.* (*Stenomegachile*) *chelostomoides* Gribodo, 1894 [Zaire] | *M.* (*Stenomegachile*) *chelostomoides* | *M.* (*Stenomegachile*) *chelostomoides* |
| *M.* (*Stenomegachile*) *dolichosoma* Benoist, 1962 [Madagascar] | — | — |
| **M.* (*Thaumatosoma*) *duboulaii* Smith, 1865 [Australia] | *M.* (*Thaumatosoma*) *remeata* Cockerell, 1913 | *M.* (*Thaumatosoma*) [dub×rem] |
| GROUP 3 | | |
| *M.* (*Creightonella*) *albisecta* Klug, 1817 [Slovakia] | *M.* (*Creightonella*) *albisecta* | *M.* (*Creightonella*) *albisecta* |
| **M.* (*Creightonella*) *cognata* Smith, 1853 [Uganda] | *M.* (*Creightonella*) *cornigera* Friese, 1904 | *M.* (*Creightonella*) [cog×cor] |
| *Noteriades jenniferae* Griswold & Gonzalez, 2011 [Thailand, Myanmar] | *Noteriades* sp. | *Noteriades* [jen×sp] |
| *Radoszkowskiana rufiventris* Spinola, 1838 [Egypt] | *R. rufiventris* | *R. rufiventris* |
