## Appendix S4 for "Evolution of leaf-cutter behavior in bees (Hymenoptera: Megachilidae) as inferred from total-evidence tip-dating analyses"

Running title: Systematics and evolution of leaf-cutter bees

**Appendix S4.** GenBank accession numbers for sequences used in this study.

| Phylogeny of Megachilidae | | | | | |
|---|---|---|---|---|---|
| *Taxa* | EF1a | Opsin | CAD | NAK | 28S |
| *Macropis nuda* | AY585155 | DQ116686 | DQ067171 | HQ995917 | HQ996008 |
| *Melitta leporina* | AY585158 | DQ116688 | DQ067174 | EF646394 | AY654529 |
| *Apis mellifera* | AF015267 | AMU26026 | DQ067178 | XM_623142 | AY703551 |
| *Exomalopsis* sp. | GU244989 | HM211835 | – | GU245110 | GU244802 |
| *Diadasia bituberculata* | GU244927 | AF344594 | – | GU245074 | GU244768 |
| *Nomada maculata* | GU245030 | AF344609 | – | GU245206 | GU244890 |
| *Ceratina calcarata* | AY585108 | AF344620 | DQ067190 | GU245213 | HQ996011 |
| *Fidelia pallidula* | HQ995686 | HQ995756 | HQ995831 | HQ995929 | HQ996025 |
| *Fidelia villosa* | HQ995682 | HQ995752 | HQ995827 | HQ995925 | HQ996021 |
| *Fidelia braunsiana* | HQ995683 | HQ995753 | HQ995828 | HQ995926 | HQ996022 |
| *Fideliopsis major* | DQ141113 | EU851628 | HQ995833 | HQ995931 | HQ996027 |
| *Fidelia profuga* | GU244990 | HQ995760 | HQ995836 | GU245151 | HQ996030 |
| *Pararhophites orobinus* | HQ995679 | HQ995749 | HQ995823 | HQ995922 | HQ996018 |
| *Pararhophites quadratus* | EU851522 | EU851627 | HQ995824 | GU245153 | GU244841 |
| *Afranthidium* (*Capanthidium*) *capicola* | KX060937 | KX060813 | KX060879 | KU976158 | KU976218 |
| *Aspidosmia arnoldi* | HQ995701 | HQ995773 | HQ995850 | HQ995945 | HQ996042 |
| *Aspidosmia volkmanni* | HQ995702 | HQ995774 | HQ995851 | HQ995946 | HQ996043 |
| *Anthidiellum* (*Loyolanthidium*) *robertsoni* | KX060952 | KX060830 | KX060894 | KU976175 | KU976235 |
| *Anthidium* (*Anthidium*) *porterae* | GU244996 | AF344619 | – | GU245158 | GU244846 |
| *Anthodioctes* (*Anthodioctes*) *mapirensis* | HQ995700 | HQ995772 | HQ995849 | HQ995944 | HQ996041 |
| *Dianthidium* (*Dianthidium*) *subparvum* | GU244993 | KX060847 | KX060909 | GU245155 | GU244843 |
| *Euaspis abdominalis* | JX869703 | JX869735 | JX869627 | JX869769 | JX869662 |
| *Hoplostelis bivittata* | JX869705 | JX869737 | – | JX869771 | JX869671 |
| *Pachyanthidium (Pachyanthidium) cordatum* | KX060971 | KX060852 | KX060915 | KU976196 | KU976257 |
| *Serapista rufipes* | HQ995716 | HQ995789 | HQ995866 | HQ995960 | HQ996057 |
| *Stelis lateralis* | | | | | |
| *Trachusa larreae* | HQ995719 | HQ995791 | HQ995868 | GU245154 | GU244842 |
| *Notanthidium steloides* | HQ995712 | HQ995784 | HQ995861 | HQ995956 | HQ996053 |
| *Plesianthidium (Spinanthidiellum) rufocaudatum* | KX060976 | KX060857 | KX060920 | KU976201 | KU976262 |
| *Icteranthidium ferrugineum flavum* | HQ995711 | HQ995783 | HQ995860 | HQ995955 | HQ996052 |
| *Hypanthidium obscurius* | HQ995710 | HQ995782 | HQ995859 | HQ995954 | HQ996051 |

| | | | | | |
|---|---|---|---|---|---|
| *Hypanthidioides marginata* | HQ995709 | HQ995781 | HQ995858 | HQ995953 | HQ996050 |
| *Eoanthidium turnericum* | HQ995707 | HQ995779 | HQ995856 | HQ995951 | HQ996048 |
| *Rhodanthidium septemdentatum* | HQ995715 | HQ995788 | HQ995865 | HQ995959 | HQ996056 |
| *Epanthidium bicoloratum* | HQ995708 | HQ995780 | HQ995857 | HQ995952 | HQ996049 |
| *Pseudoanthidium (Micranthidium) sp.* | KX060982 | KX060862 | KX060926 | KU976206 | KU976268 |
| *Aztecanthidium tenochtitlanicum* | – | KX060844 | KX060906 | KU976189 | KU976249 |
| *Duckeanthidium thielei* | HQ995706 | HQ995778 | HQ995855 | HQ995950 | HQ996047 |
| *Cyphanthidium intermedium* | KX060966 | KX060845 | KX060907 | KU976190 | KU976250 |
| *Aglaoapis tridentata* | EU851524 | EU851630 | HQ995844 | HQ995939 | HQ996036 |
| *Dioxys moesta* | HQ995696 | HQ995768 | HQ995845 | HQ995940 | HQ996037 |
| *Lithurgus chrysurus* | EU851523 | EU851629 | HQ995837 | HQ995934 | HQ996031 |
| *Microthurge sp* | HQ995694 | HQ995766 | HQ995842 | GU245161 | GU244849 |
| *Trichothurgus herbsti* | HQ995695 | HQ995767 | HQ995843 | GU245160 | GU244848 |
| *Afroheriades primus* | EU851532 | EU851638 | HQ995902 | HQ995995 | HQ996092 |
| *Ashmeadiella aridula* | EU851535 | EU851641 | HQ995903 | GU245171 | GU244858 |
| *Atoposmia mirifica* | EU851541 | EU851647 | HQ995904 | HQ995996 | HQ996093 |
| *Chelostoma florisomne* | EU851546 | EU851652 | HQ995905 | HQ995997 | HQ996094 |
| *Haetosmia brachyura* | HQ995748 | HQ995822 | HQ995906 | HQ995998 | HQ996095 |
| *Heriades crucifer* | EU851555 | EU851661 | DQ067194 | GU245168 | GU244855 |
| *Hofferia schmiedeknechti* | EU851556 | EU851662 | HQ995907 | HQ995999 | HQ996096 |
| *Hoplitis adunca* | EU851572 | EU851678 | HQ995908 | HQ996000 | HQ996097 |
| *Ochreriades fasciatus* | EU851590 | EU851696 | HQ995909 | HQ996001 | HQ996098 |
| *Osmia lignaria* | EU851610 | EU851715 | HQ995910 | GU245169 | GU244856 |
| *Othinosmia globicola* | EU851616 | EU851721 | HQ995911 | HQ996002 | HQ996099 |
| *Protosmia humeralis* | EU851621 | EU851726 | HQ995913 | HQ996004 | HQ996101 |
| *Pseudoheriades moricei* | EU851622 | EU851727 | HQ995914 | HQ996005 | HQ996102 |
| *Stenoheriades asiaticus* | EU851623 | EU851728 | HQ995915 | HQ996006 | HQ996103 |
| *Hoplitis minima* | EU851625 | EU851730 | EU851520 | – | – |
| *Wainia eremoplana* | EU851626 | EU851731 | HQ995916 | HQ996007 | HQ996104 |
| *Noteriades sp* | EU851589 | EU851695 | HQ995900 | HQ995993 | HQ996090 |
| *Coelioxys octodentata* | KX428310 | KX428056 | KX428226 | KX428394 | KX428151 |
| *Megachile pugnata* | AY585147 | HQ995818 | DQ067196 | HQ995990 | HQ996087 |
| *Megachile angelarum* | HQ995727 | HQ995800 | HQ995878 | GU245163 | GU244851 |
| *Megachile albisecta* | EU851529 | EU851635 | HQ995881 | HQ995974 | HQ996071 |
| *Radoszkowskiana rufiventris* | HQ995747 | HQ995821 | HQ995901 | HQ995994 | HQ996091 |

| | | | | | |
|---|---|---|---|---|---|
| *Trichothurgus herbsti* | HQ995695 | HQ995767 | HQ995843 | GU245160 | GU244848 |
| *Microthurge sp* | HQ995694 | HQ995766 | HQ995842 | GU245161 | GU244849 |
| *Aspidosmia volkmanni* | HQ995702 | HQ995774 | HQ995851 | HQ995946 | HQ996043 |
| *Trachusa larreae* | HQ995719 | HQ995791 | HQ995868 | GU245154 | GU244842 |
| *Hoplitis adunca* | EU851572 | EU851678 | HQ995908 | HQ996000 | HQ996097 |
| *Chelostoma florisomne* | EU851546 | EU851652 | HQ995905 | HQ995997 | HQ996094 |
| *Dioxys moesta* | HQ995696 | HQ995768 | HQ995845 | HQ995940 | HQ996037 |
| *Aztecanthidium tenochtitlanicum* | – | KX060844 | KX060906 | KU976189 | KU976249 |
| *Radoszkowskiana rufiventris* | HQ995747 | HQ995821 | HQ995901 | HQ995994 | HQ996091 |
| *Coelioxys decipiens* | KX428313 | KX428059 | KX428229 | KX428397 | KX428154 |
| *Noteriades sp* | EU851589 | EU851695 | HQ995900 | HQ995993 | HQ996090 |
| *M. (Rhodomegachile) sp.* | HQ995744 | HQ995817 | HQ995897 | HQ995989 | HQ996086 |
| *M. (Matangapis) alticola* | – | – | – | – | KX580315 |
| M (Chelostomoda) sp | HQ995726 | HQ995799 | Missing | HQ995971 | HQ996068 |
| M (Heriadopsis) sp | – | – | – | – | KX580316 |
| M (Hackeriapis) sp 1 | KX428356 | KX428102 | KX428272 | KX428438 | KX428195 |
| M (Chelostomoidella) spinotulata | HQ995728 | HQ995801 | HQ995879 | HQ995972 | HQ996069 |
| M (Chelostomoides) angelarum | HQ995727 | HQ995800 | HQ995878 | GU245163 | GU244851 |
| M (Thaumatosoma) remeata | HQ995745 | HQ995819 | HQ995898 | HQ995991 | HQ996088 |
| M (Stenomegachile) chelostomoides | KX428383 | KX428129 | KX428299 | KX428461 | KX428218 |
| M (Maximegachile) maxillosa | HQ995737 | HQ995810 | HQ995890 | HQ995983 | HQ996080 |
| M (Chalicodomoides) aethiops | HQ995725 | HQ995798 | HQ995877 | HQ995970 | HQ996067 |
| M (Carinula) decemsignata | KX428325 | KX428071 | KX428242 | KX428409 | KX428166 |
| M (Callomegachile) sculpturalis | HQ995724 | HQ995797 | HQ995875 | HQ995968 | HQ996065 |
| M (Alocanthedon) sp | KX428318 | KX428064 | KX428235 | KX428402 | KX428159 |
| M (Gronoceras) bombiformis | HQ995733 | HQ995806 | HQ995886 | HQ995979 | HQ996076 |
| M (Lophanthedon) dimidiata | KX428359 | KX428105 | KX428275 | KX428441 | KX428198 |
| M (Austrochile) sp | HQ995723 | HQ995796 | HQ995874 | HQ995967 | HQ996064 |
| M (Parachalicodoma) sp | KX428370 | KX428116 | KX428286 | KX428448 | KX428205 |
| M manicata | KX428330 | KX428076 | KX428247 | KX428413 | KX428170 |
| M (Chalicodoma) parietina | EU851530 | EU851636 | HQ995876 | HQ995969 | HQ996066 |
| M (Allochalicodoma) lefebvrei | KX428329 | KX428075 | KX428246 | KX428412 | KX428169 |

| | | | | | |
|---|---|---|---|---|---|
| M (Largella) floralis | HQ995735 | HQ995808 | HQ995888 | HQ995981 | HQ996078 |
| M (Pseudomegachile) ericetorum | HQ995742 | HQ995815 | HQ995895 | GU245165 | GU244853 |
| M (Neglectella) laminata | KX428366 | KX428112 | KX428282 | KX428444 | KX428201 |
| M (Dinavis) leucospilura | KX428336 | KX428082 | KX428252 | KX428418 | KX428175 |
| M (Cesacongoa) sp | KX428328 | KX428074 | KX428245 | KX428411 | |
| M (Creightonella) albisecta | EU851529 | EU851635 | HQ995881 | HQ995974 | HQ996071 |
| M (Creightonella) cornigera | KX428334 | KX428080 | KX428250 | KX428416 | KX428173 |
| M (Mitchellapis) fabricator | HQ995740 | HQ995813 | HQ995893 | HQ995986 | HQ996083 |
| *Megachile pugnata* | AY585147 | HQ995818 | DQ067196 | HQ995990 | HQ996087 |
| M (Grosapis) cockerelli | KX428355 | KX428101 | KX428271 | KX428437 | KX428194 |
| M (Eumegachile) bombycina | KX428337 | KX428083 | KX428253 | KX428419 | KX428176 |
| M (Litomegachile) texana | HQ995736 | HQ995809 | HQ995889 | HQ995982 | HQ996079 |
| M (Neoeutricharaea) rotundata | XM_003705302 | XM_003705921 | XM_012287299 | XM_012290255 | – |
| M (Eurymella) aff. eurymera | KX428338 | KX428084 | KX428254 | KX428420 | KX428177 |
| M (Argyropile) parallela | HQ995722 | HQ995795 | HQ995873 | HQ995966 | HQ996063 |
| M (Amegachile) cf. bituberculata | KX428319 | KX428065 | KX428236 | KX428403 | KX428160 |
| M (Xeromegachile) nevadensis | HQ995739 | HQ995812 | HQ995892 | HQ995985 | HQ996082 |
| M (Phaenosarus) fortis | KX428371 | KX428117 | KX428287 | KX428449 | KX428206 |
| M (Xanthosarus) lagopoda | KX428390 | KX428136 | KX428306 | – | – |
| M (Leptorachis) petulans | KX428357 | KX428103 | KX428273 | KX428439 | KX428196 |
| M (Pseudocentron) sp | KX428372 | KX428118 | KX428288 | KX428450 | KX428207 |
| M (Melanosarus) sp | KX428365 | KX428111 | KX428281 | KX428443 | KX428200 |
| M (Acentron) sp | KX428316 | KX428062 | KX428233 | KX428400 | KX428157 |
| M (Paracella) sp | KX428368 | KX428114 | KX428284 | KX428446 | KX428203 |
| M (Tylomegachile?) sp | KX428384 | KX428130 | KX428300 | KX428462 | KX428219 |
| M (Aethomegachile) conjuncta | HQ995720 | HQ995793 | HQ995871 | HQ995964 | HQ996061 |
| M (Megella) pseudomonticola | KX428364 | KX428110 | KX428280 | KX428442 | KX428199 |
| M (Dasymegachile) sp | KX428335 | KX428081 | KX428251 | KX428417 | KX428174 |
| M (Cressoniella) zapoteca | HQ995730 | HQ995803 | HQ995882 | HQ995975 | HQ996072 |
| M (Austromegachile) sp | KX428322 | KX428068 | KX428239 | KX428406 | KX428163 |
| M (Ptilosarus) microsoma | HQ995743 | HQ995816 | HQ995896 | HQ995988 | HQ996085 |
| M (Neochelynia?) sp | KX428367 | KX428113 | KX428283 | KX428445 | KX428202 |
| M (Stelodides) euzona | KX428382 | KX428128 | KX428298 | KX428460 | KX428217 |
| M (Chrysosarus) sp | HQ995729 | HQ995802 | HQ995880 | HQ995973 | HQ996070 |
