## Appendix S5 for "Evolution of leaf-cutter behavior in bees (Hymenoptera: Megachilidae) as inferred from total-evidence tip-dating analyses"

Running title: Systematics and evolution of leaf-cutter bees

CONTENTS

**S5A.** Quantitative descriptors of trees obtained from implied weighting (IW) analyses……......2
**S5B.** Tree obtained from IW analysis using K value of 8.81 (F: 54%)………………………….3
**S5C.** Tree obtained from IW analysis using K value of 12.2 (F: 62%)………………………….4
**S5D.** Tree obtained from IW analysis using K value of 14.6. (F: 66%)…………………………5
**S5E.** Tree obtained from IW analysis using K value of 17.5 (F: 70%)………………………….6

**Appendix S5A.** Quantitative descriptors of trees obtained from implied weighting (IW) analyses. F = total fit of characters to tree (Goloboff, 1993); K = concavity factor determining weighting strength; MPT = number of most parsimonious trees; L = tree length; RI = retention index; GC Observed = average GC frequency-difference, as calculated from displayed support at each node in the resulting tree. Average value followed, in parentheses, by median, standard deviation, and number of nodes. Top highest values are in boldface. * = a single node collapsed in the consensus tree. In all analyses, the Consistency Index was 13. Avg. SPR-dist. EW consensus = SPR distance between the resulting topology of each IW analysis and the topology obtained from the consensus tree of the equal weighting analysis. Avg. SPR-dist. Total evidence = SPR distance between the resulting topology of each IW analysis and the topology obtained from the Bayesian inference analysis of the full dataset. High values in bold face.

| F | K | MPT | L | RI | GC Observed | Avg. SPR-dist EW consensus | Avg. SPR-dist Total Evidence |
|---|---|---|---|---|---|---|---|
| 0.5 | 7.50 | 1 | 2434 | 56 | **50.24** (48.0, ± 35.11, *n* = 63) | 0.5966 | 0.4538 |
| 0.54 | 8.81 | 1 | 2421 | 56 | **50.97** (54.5, ± 35.36, *n* = 60) | 0.6134 | **0.4874** |
| 0.58 | 10.40 | 1 | 2415 | 56 | **51.03** (55.0, ± 35.87, *n* = 62) | 0.6387 | 0.4622 |
| 0.62 | 12.20 | 1 | 2397 | 57 | 49.31 (42.0, ± 36.27, *n* = 65) | 0.6891 | **0.4874** |
| 0.66 | 14.60 | 2* | 2393 | 57 | **50.19** (46.5, ± 36.08, *n* = 64) | 0.6807 | **0.4874** |
| 0.7 | 17.50 | 2* | 2389 | 57 | 48.15 (40.0, ± 36.37, *n* = 67) | 0.6639 | **0.4874** |
| 0.74 | 21.40 | 1 | 2378 | 57 | 48.03 (40.0, ± 35.79, *n* = 68) | 0.7815 | 0.4790 |
| 0.78 | 26.60 | 1 | 2377 | 57 | 48.73 (41.0, ± 35.18, *n* = 67) | 0.7899 | 0.4538 |
| 0.82 | 34.20 | 1 | 2377 | 57 | 47.38 (38.0, ± 35.43, *n* = 69) | 0.7899 | 0.4538 |
| 0.86 | 46.10 | 1 | 2372 | 57 | 48.27 (38.5, ± 35.33, *n* = 66) | 0.7563 | 0.4286 |
| 0.9 | 67.50 | 1 | 2366 | 57 | 49.67 (41.0, ± 34.88, *n* = 64) | **0.8824** | 0.3782 |

**Appendix S5B.** Tree obtained from IW analysis using K value of 8.81 (F: 54%)

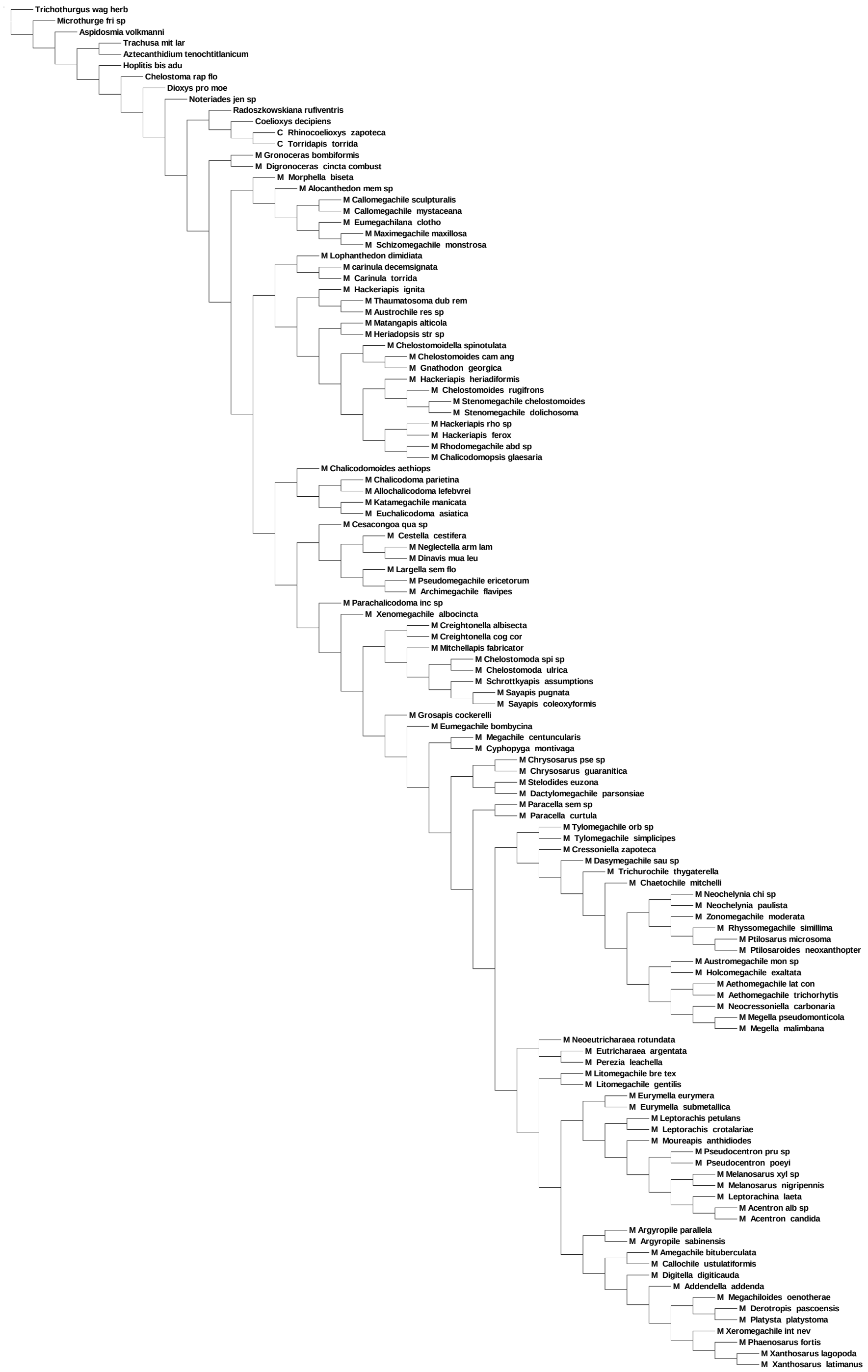

**Appendix S5C.** Tree obtained from IW analysis using K value of 12.2 (F: 62%)

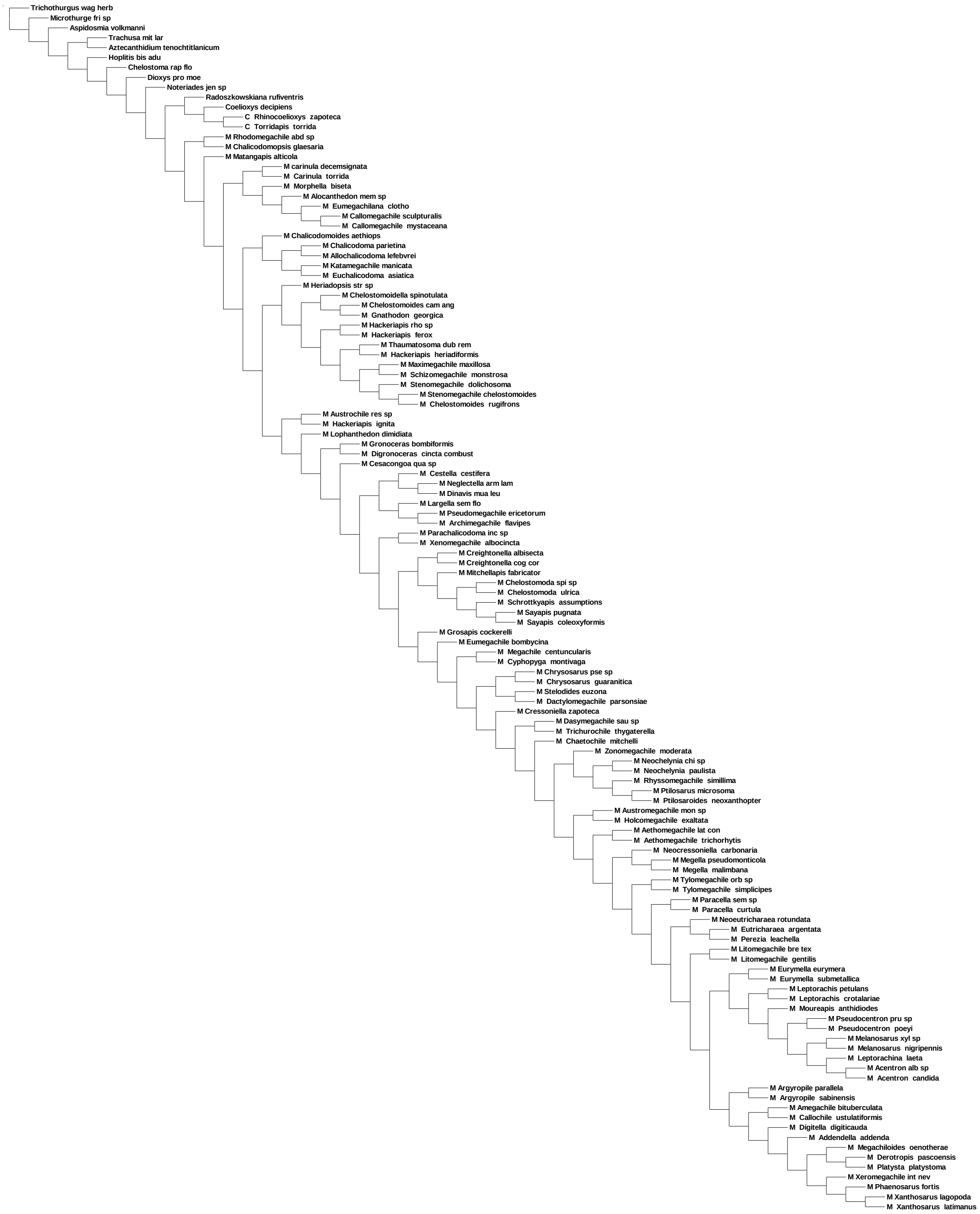

**Appendix S5D.** Tree obtained from IW analysis using K value of 14.6. (F: 66%)

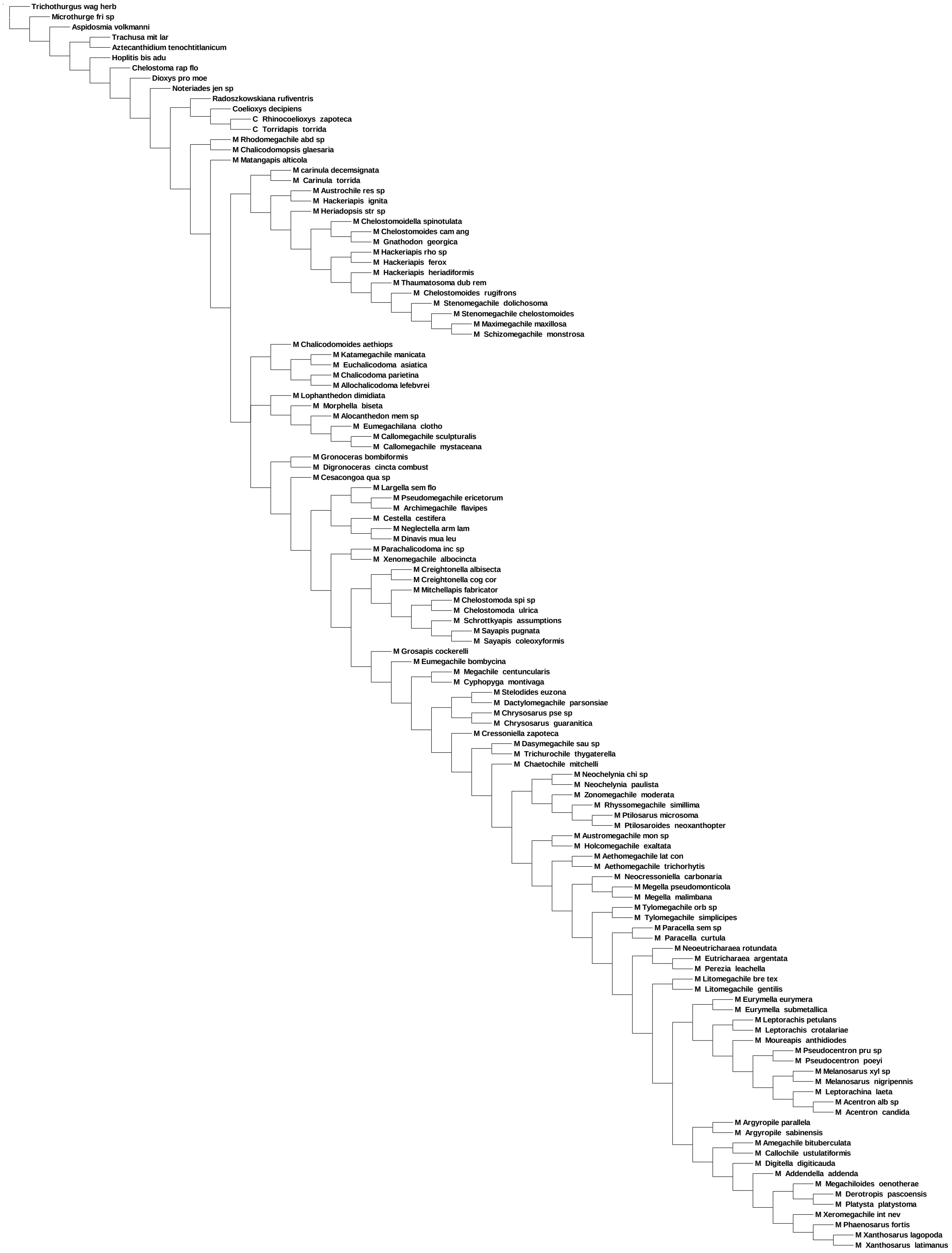

**Appendix S5E.** Tree obtained from IW analysis using K value of 17.5 (F: 70%)

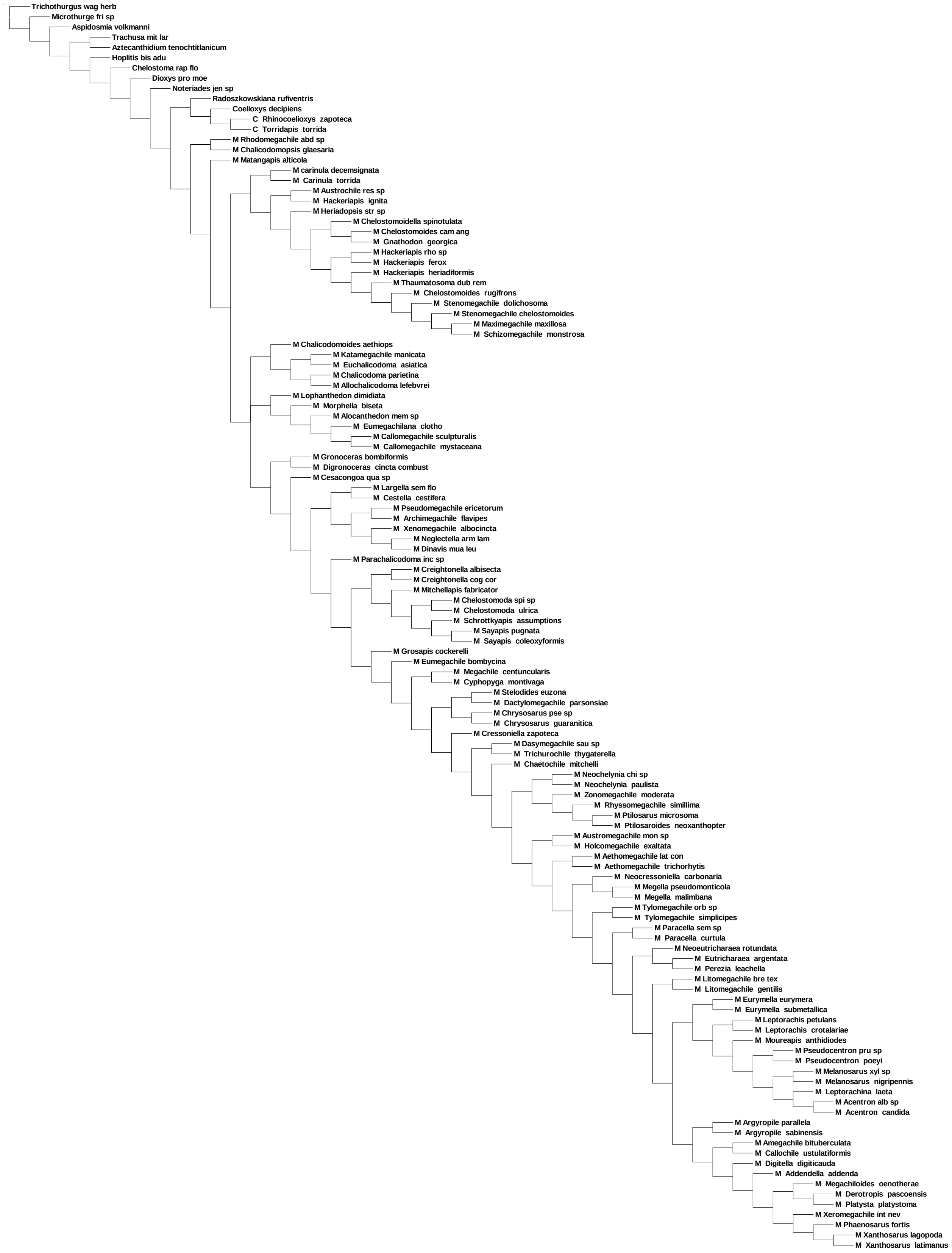
