## Appendix S6 for "Evolution of leaf-cutter behavior in bees (Hymenoptera: Megachilidae) as inferred from total-evidence tip-dating analyses"

Running title: Systematics and evolution of leaf-cutter bees

**Appendix S6.** Maximum likelihood (ML) and Bayesian inference (BI) analyses.

Tribal-level phylogeny:
**S6A.** ML analysis of molecular dataset with introns........................................................2
**S6B.** ML analysis of molecular dataset without introns....................................................3
**S6C.** ML total evidence analysis using molecular dataset without introns........................4
**S6D.** BI time free analysis..................................................................................................5
**S6E.** BI tip-dated analysis using MCMC generation set to 200 million............................6

Generic-level phylogeny:
**S6F.** ML analysis of molecular dataset with introns........................................................7
**S6G.** ML analysis of dataset without introns....................................................................8
**S6H.** ML total evidence analysis using molecular dataset with introns............................9
**S6I.** BI time free analysis..................................................................................................10
**S6J.** BI tip-dated analysis constraining *M. glaesaria* near *M.* (*Thaumatosoma*).............11
**S6K.** BI tip-dated analysis constraining *M. glaesaria* near *M.* (*Matangapis*)..................12

**S6A. Maximum likelihood analysis of the bee family Megachilidae using dataset with intronsS6C.**

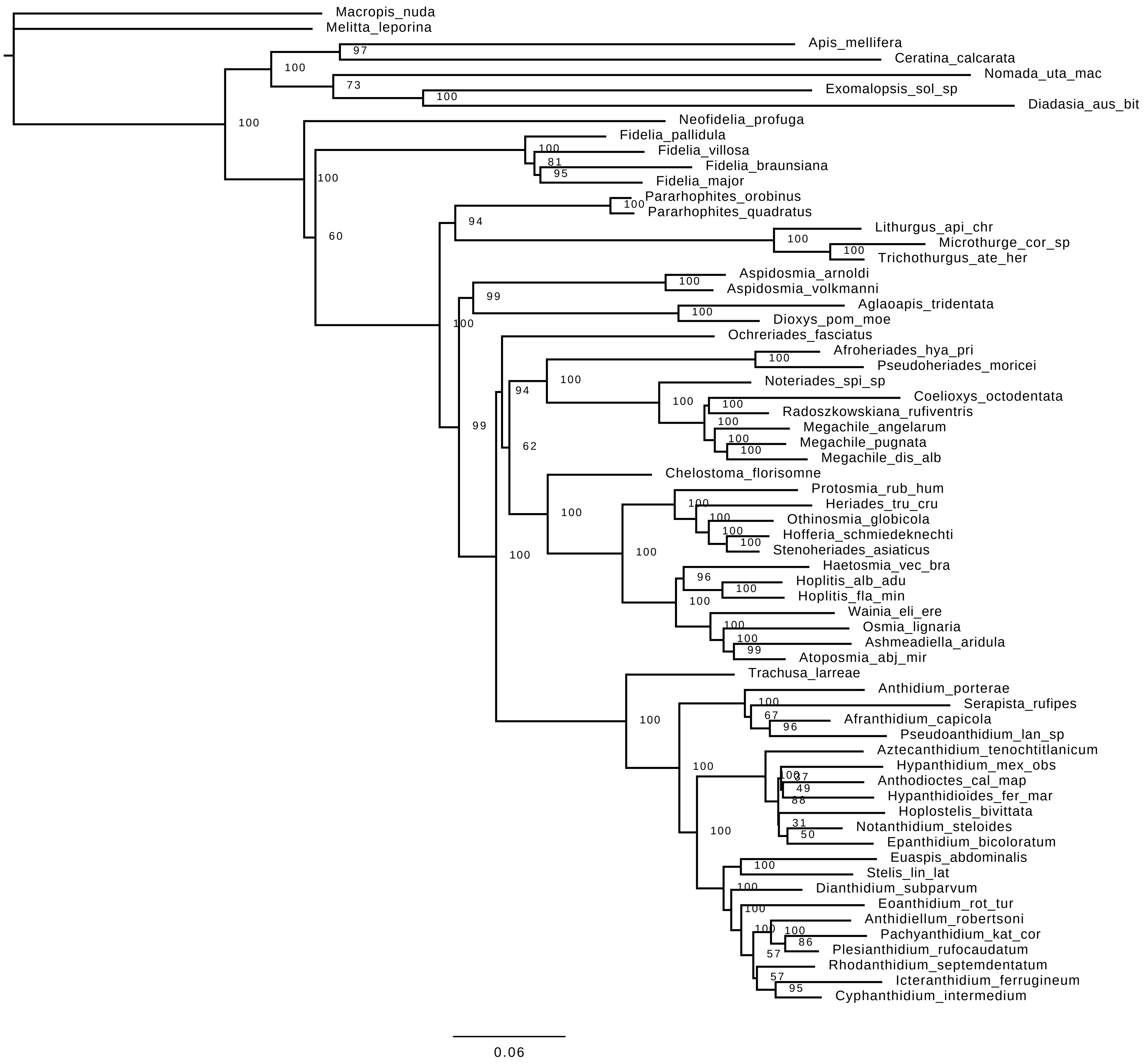

**S6B. Maximum likelihood analysis of the bee family Megachilidae using dataset without introns**

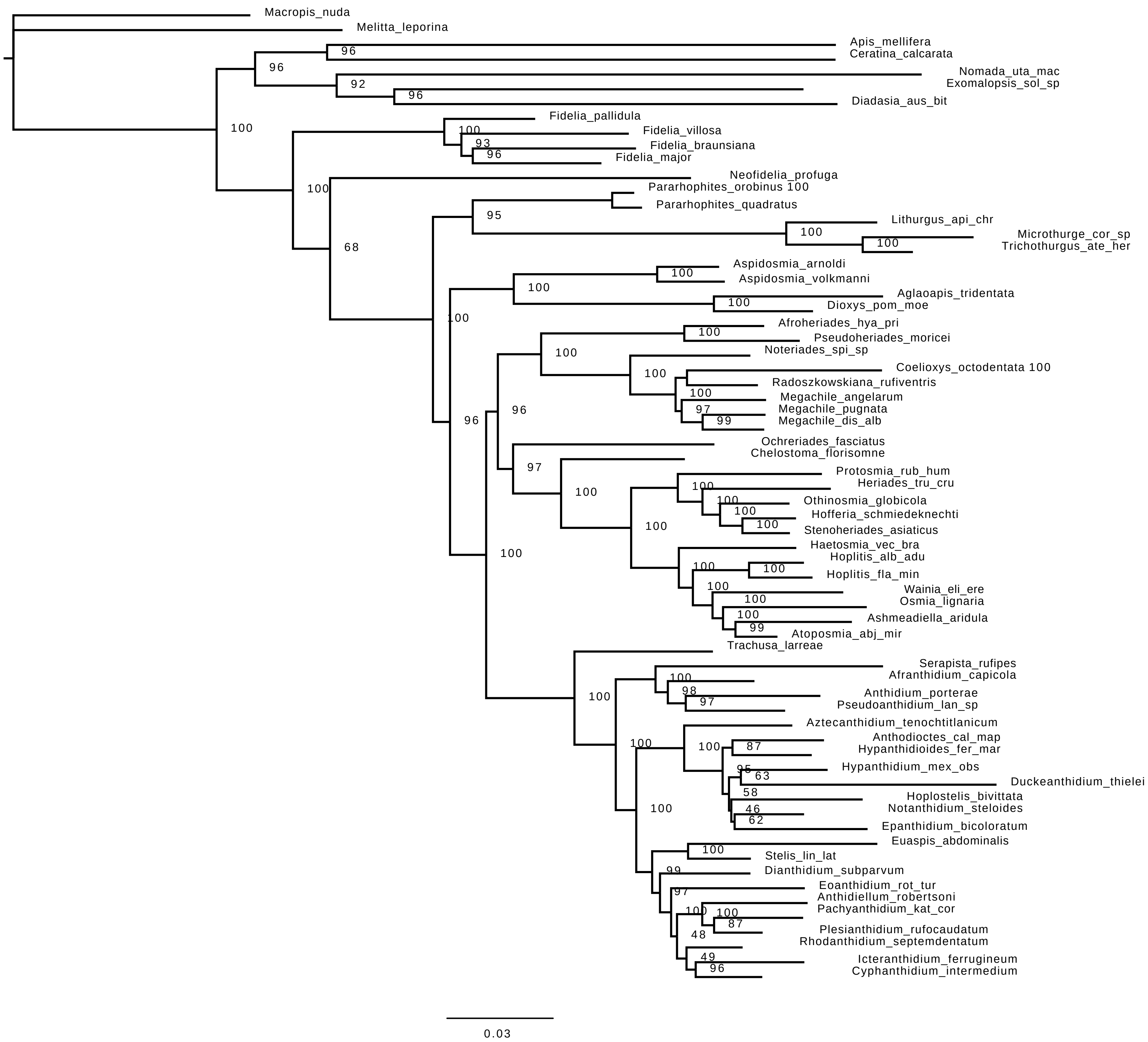

**S6C. ML total evidence analysis of Megachilidae using molecular dataset without introns**

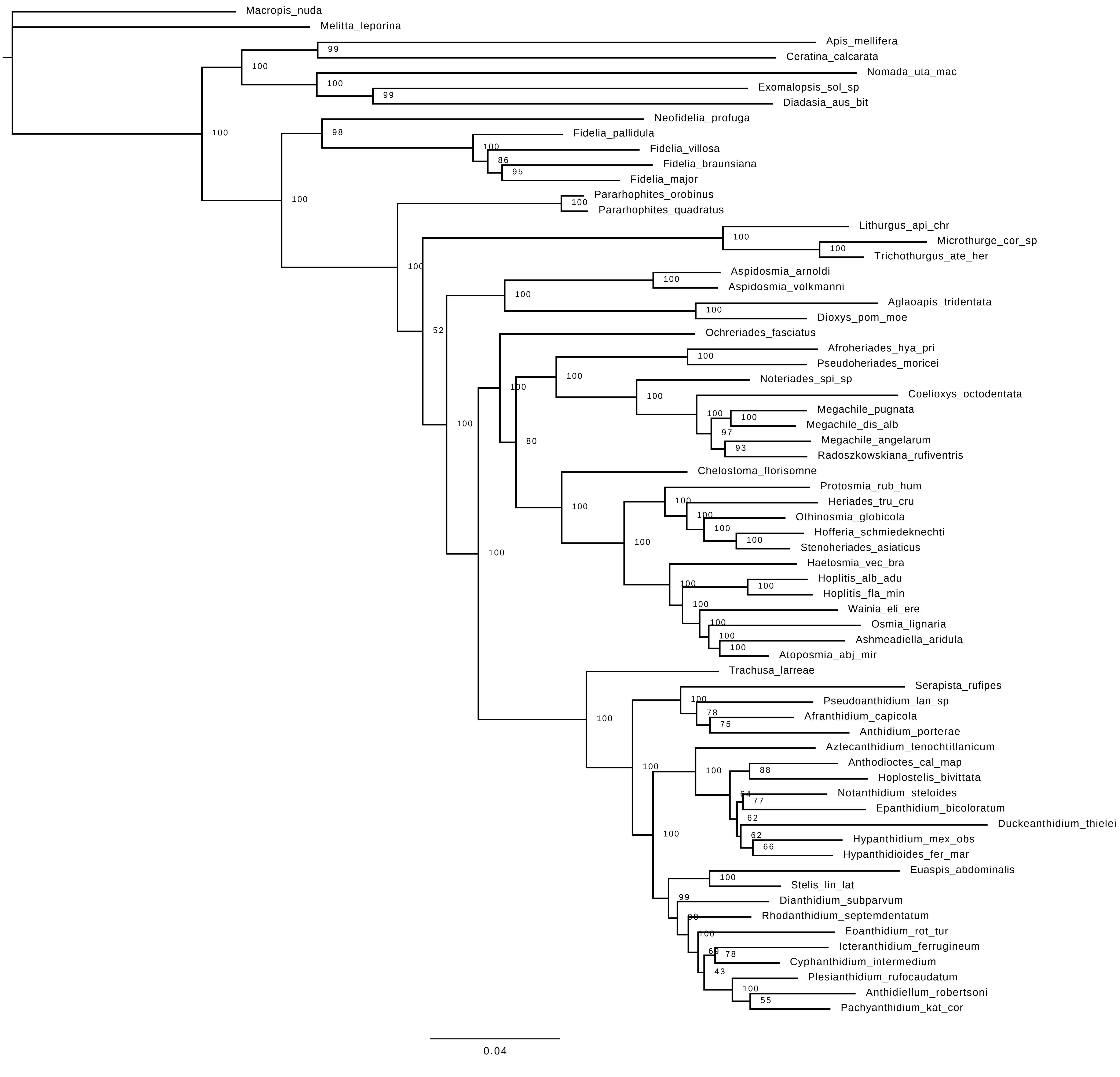

### S6D. BI total evidence time free analysis of Megachilidae

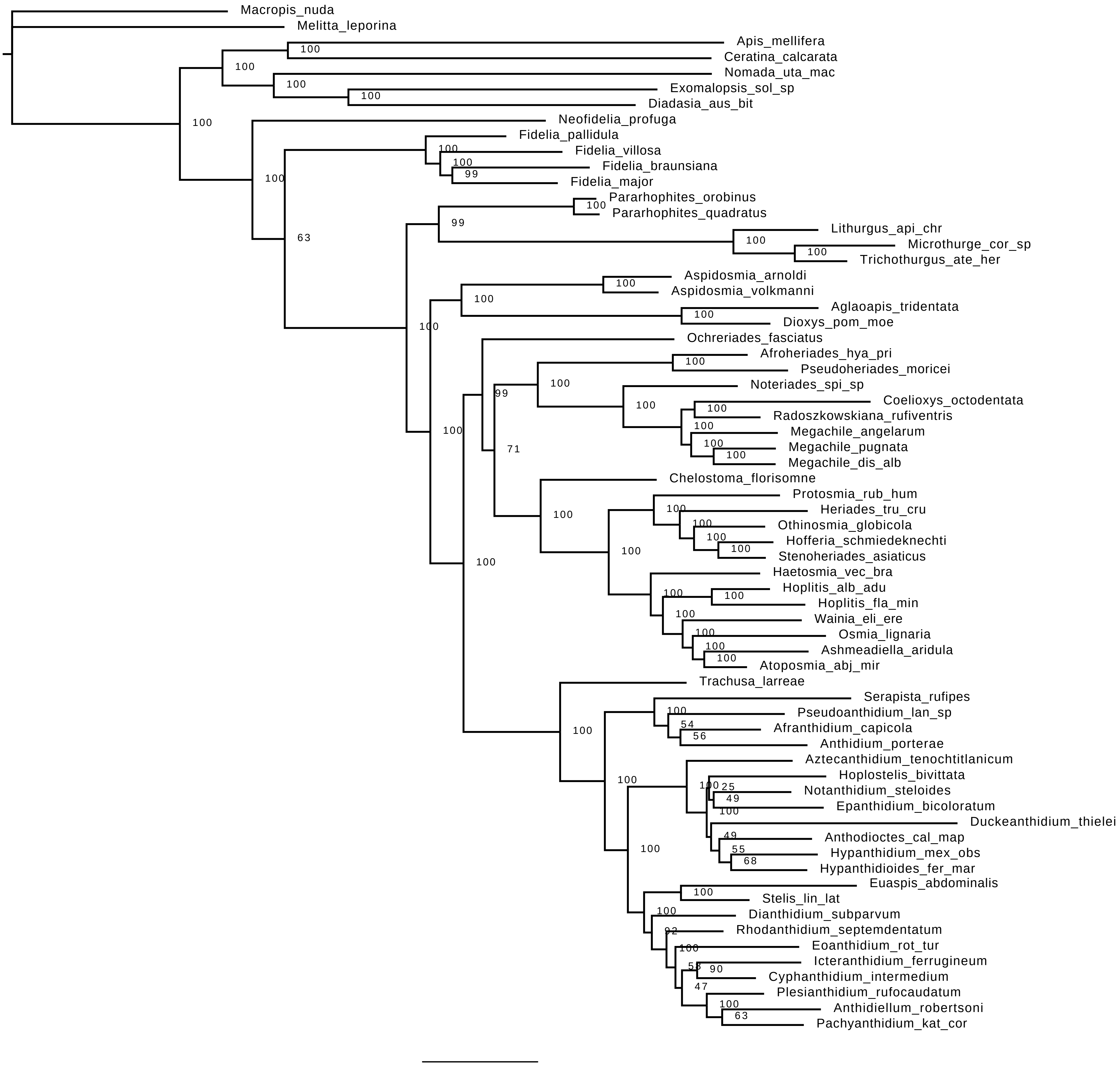

### S6E. BI tip-dated analysis of Megachilidae using 200 million MCMC generations

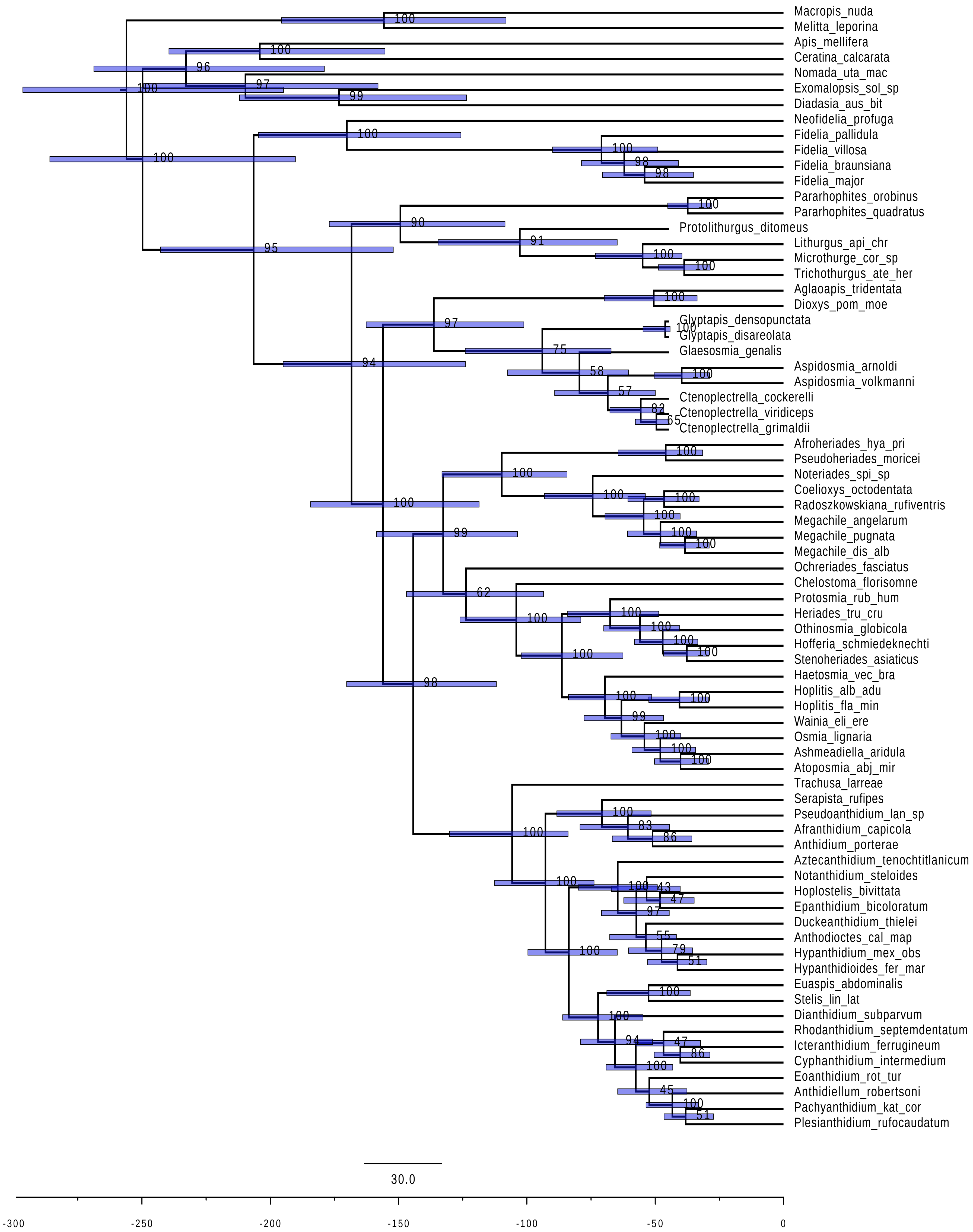

**S6F. Maximum likelihood analysis of the bee tribe Megachilini using dataset with introns**

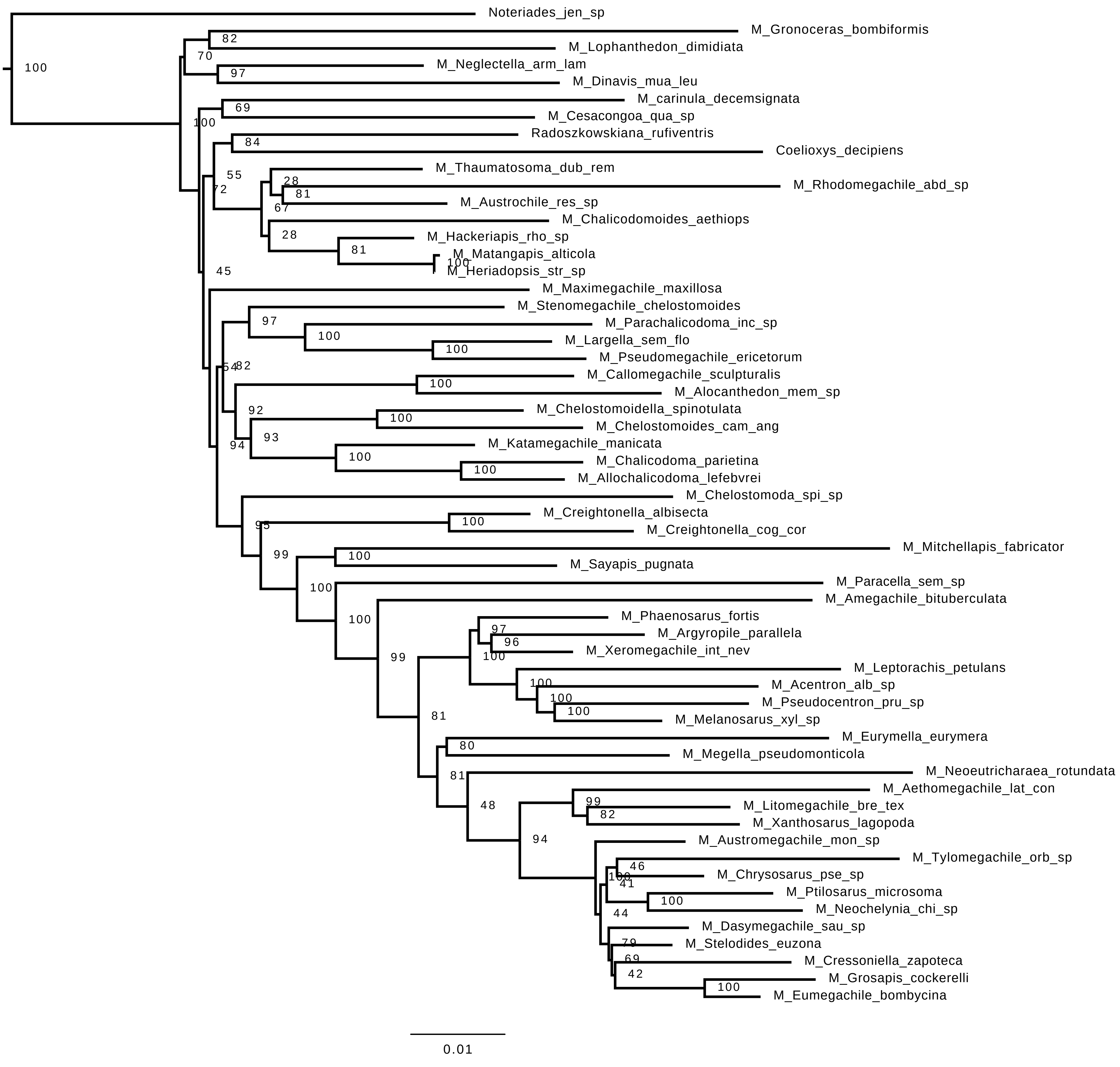

**S6G. Maximum likelihood analysis of the bee tribe Megachilini using dataset without introns**

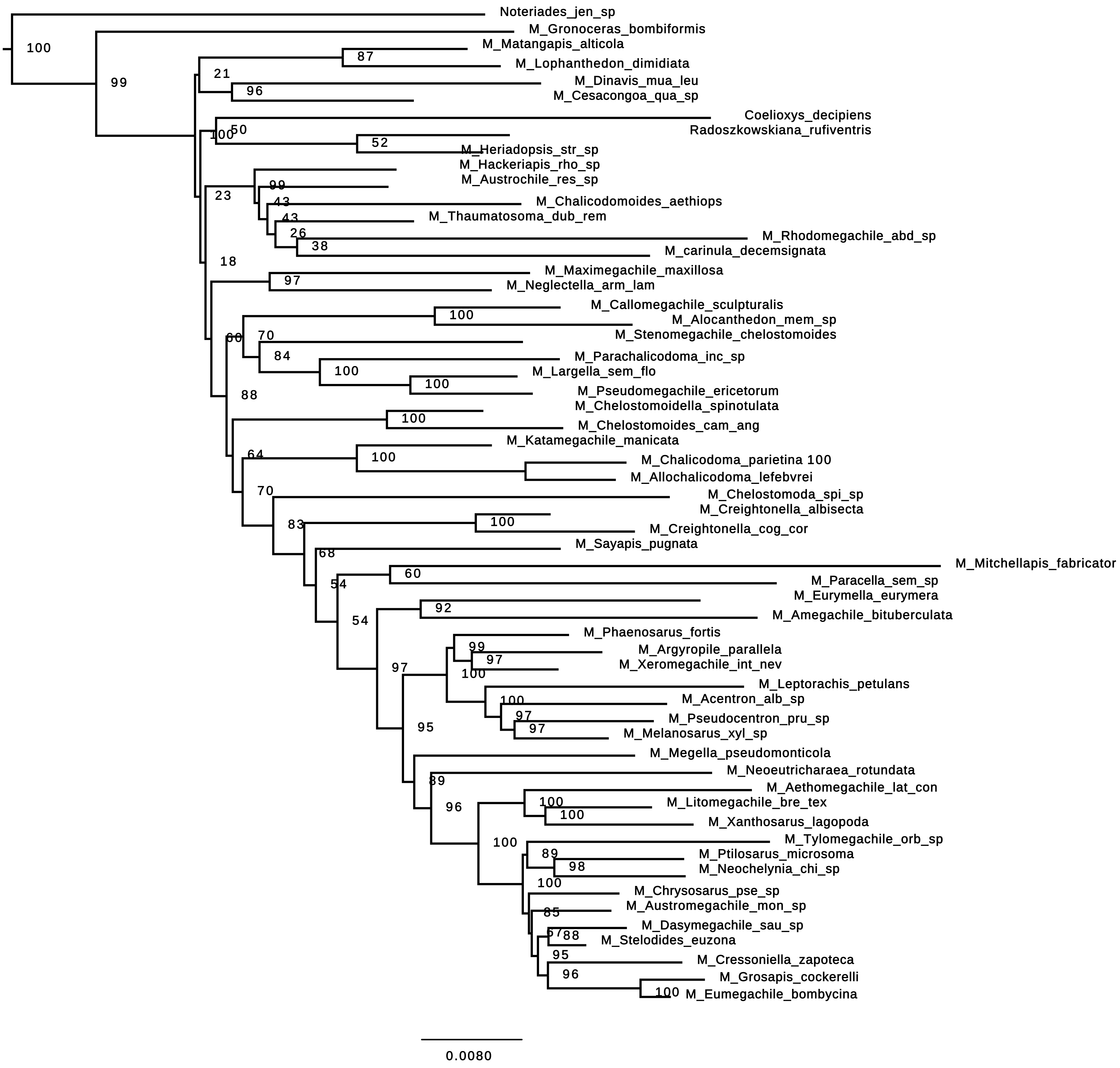

#### S6H. ML total evidence analysis of Megachilini using molecular dataset with introns

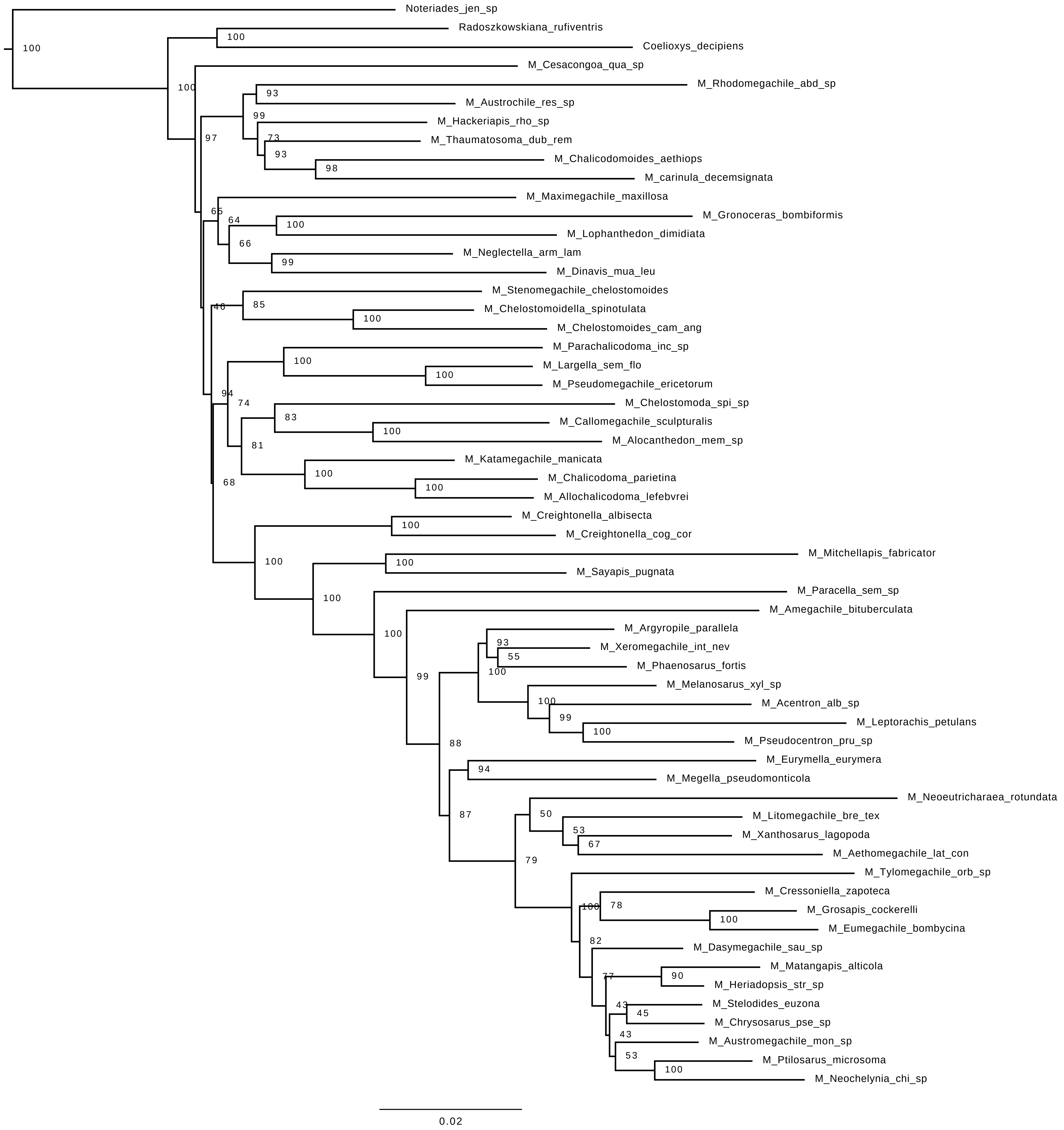

#### S6I. BI total evidence time free analysis of Megachilini

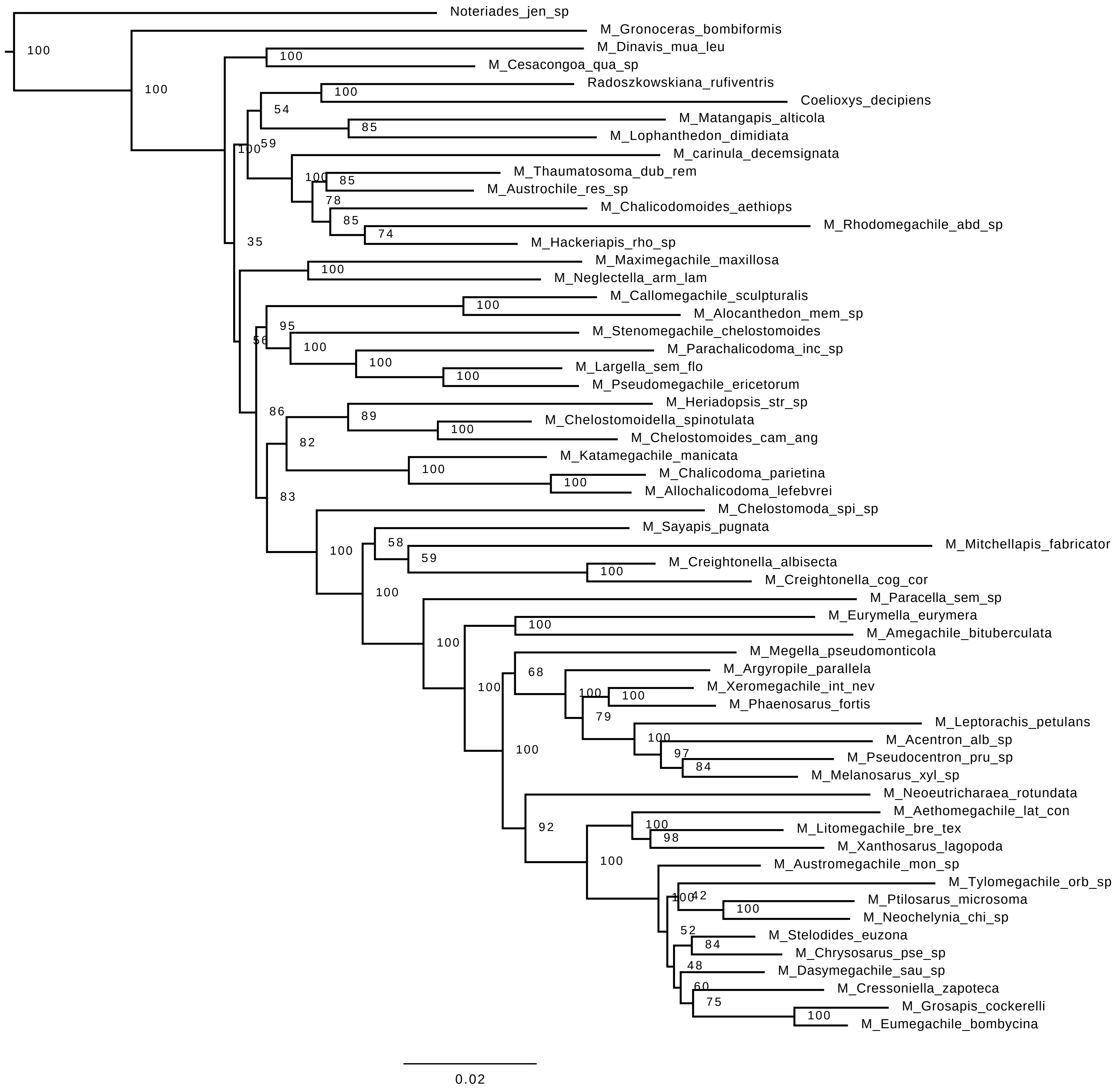

#### S6J. BI tip-dated analysis of Megachilini constraining *M. glaesaria* near *M.* (*Thaumatosoma*)

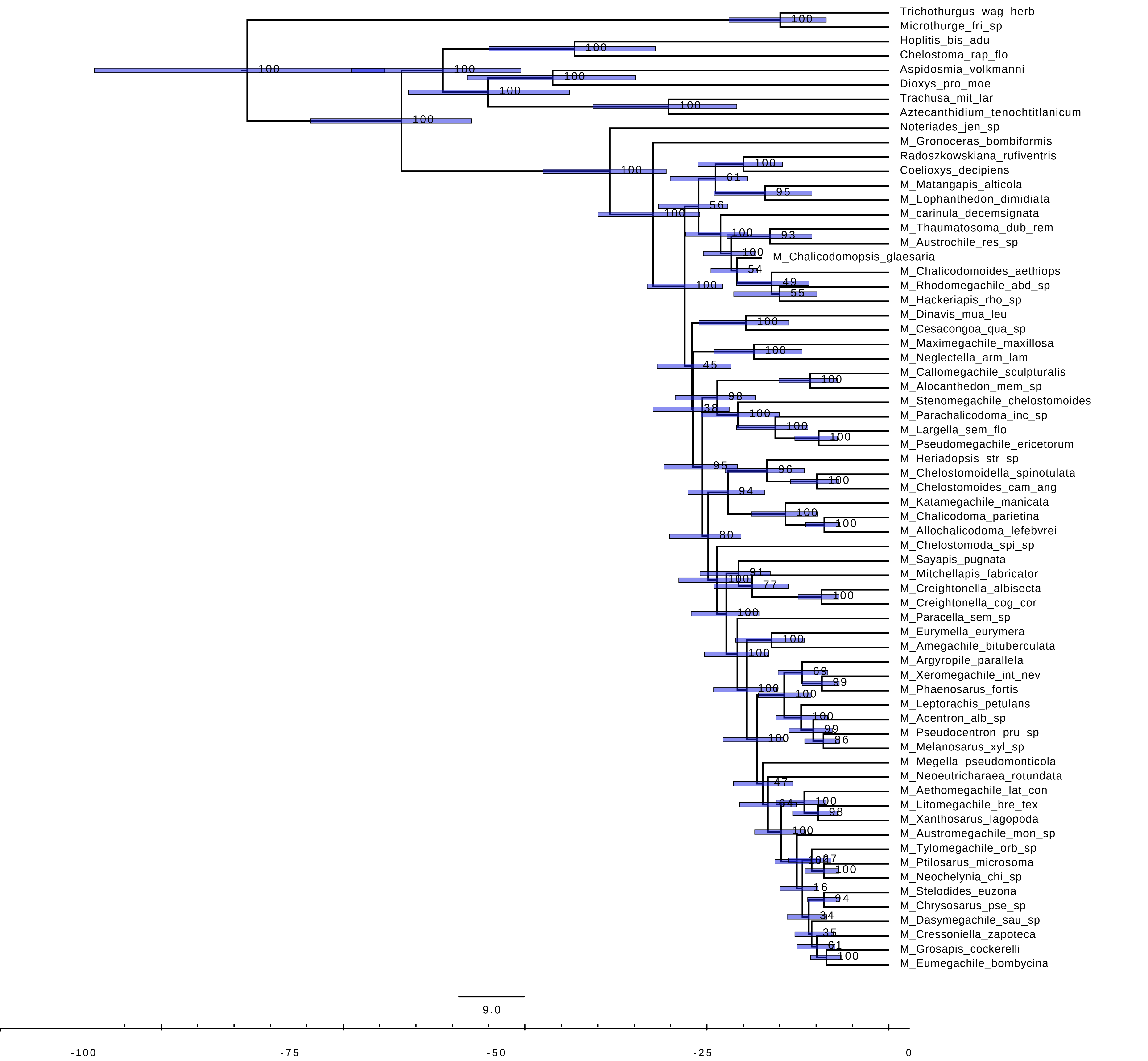

#### S6K. BI tip-dated analysis of Megachilini constraining *M. glaesaria* near *M.* (*Matangapis*)

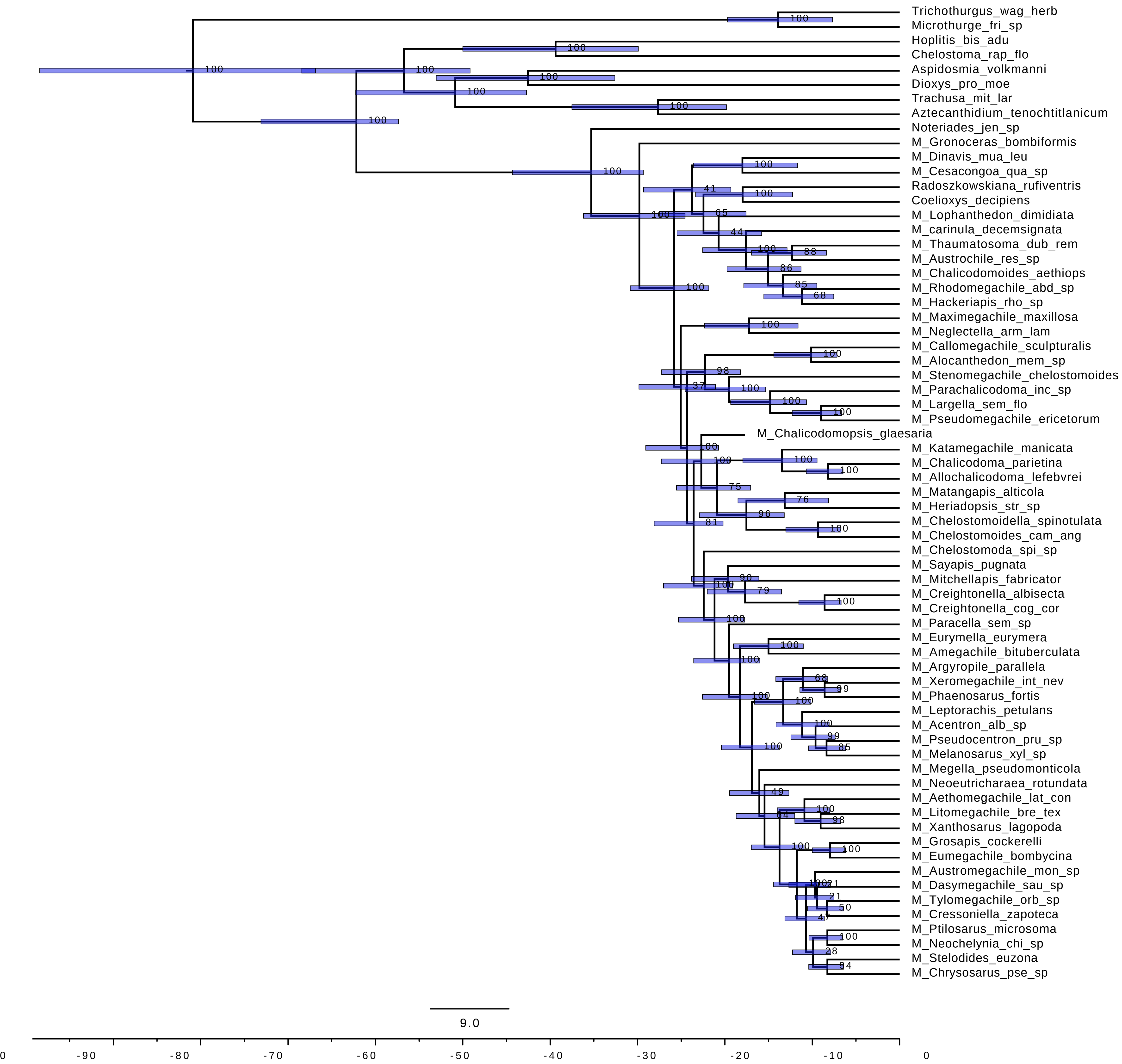
